# Fructose Dendrimer-Based Poly Ionic Complexes for Glut5-Specific Intracellular Drug Delivery in Murine Brain

**DOI:** 10.64898/2026.09.24.754190

**Authors:** P.S. Woods, G. Aldhahri, E. Hanchar, S. Dhulekar, Espinoza R. Ramos, A. Piñeiro, J.P Vincent, T.M. O’Shea

## Abstract

Glial cells play essential roles in maintaining neural tissue homeostasis and limiting the spread of injury-induced neuroinflammation, yet their dysfunction can also exacerbate neurodegenerative, neuroimmune, and neurodevelopmental disorders. Therapeutic targeting of specific glial cell populations would open up new treatment possibilities, but currently available viral and non-viral delivery vehicles are limited by carrier toxicity as well as poor delivery efficiency and specificity. To work towards new glia-targeted, non-viral delivery vehicles, we performed an *in silico* analysis on neural cell specific RNA-Seq datasets in mice to identify surface molecules differentially enriched on neurons, astrocytes, and microglia. We identified *Slc2a5* (Glut5) as a microglia-enriched transporter and validated its microglia specific expression in healthy murine neural tissue using RNA in situ hybridization. By employing a sequential thiol-ene and enzymatic catalyzed reaction scheme, we synthesized polyelectrolytic dendrimers with multivalent presentation of fructose, the natural ligand for Glut5, and formulated nano-sized polyionic complexes (PICs) loaded with protein or nucleic acid cargo by mixing oppositely charged dendrimers. Fructose functionalized PICs lacking cationic surface charge enabled fructose-dependent intracellular protein delivery to Glut5-expressing human breast cancer cells (MCF-7) and mouse fibroblasts, while restricting uptake by Glut5-negative mouse astrocytes, confirming transporter-dependent selectivity *in vitro*. Fructose functionalized PICs resulted in minimal neuroinflammation when injected into mouse striatum at 1 mg/ml but elicited concentration-dependent neural tissue toxicity at higher doses. Fructose-PICs failed to achieve microglia specific intracellular delivery of functional proteins when injected into healthy neural tissue but aided in directing protein delivery to microglia and astrocytes at ischemic stroke lesions. Together, our findings demonstrate proof-of-principle for using in silico RNA-seq screening workflows to identify cell specific surfaceome targets and incorporating that insight into a modular biomaterial platform for targeted drug delivery.

## Introduction

Central nervous system (CNS) parenchymal tissue is composed of a collection of glia which tile the entire volume of the brain in a contiguous, non-overlapping manner to support neuronal functions and maintain healthy brain homeostasis^1–3^. Glia dysfunction evoked by environmental factors or genetic mutations can compromise neural tissue function, and is implicated in the cause and exacerbation of numerous neurodegenerative diseases^4^ including Parkinson’s (PD)^5^, Huntington’s (HD)^6^ and Alzheimer’s (AD)^7^ as well as gliomas^8^, neuroimmune^9^ and neurodevelopmental^10^ disorders. Microglia, a specific class of glia responsible for innate immunity in the CNS^2^, have emerged as important regulators and drivers of neuroinflammation in neurodegenerative diseases^11,12^. In these diseases, microglia adopt reactive states early on in disease progression and may contribute to neuronal damage rather than simply responding to it^11,13,14^. While healthy microglia can clear misfolded or damaged proteins, when microglia functions are compromised or there is a high disease burden, the resulting accumulation and aggregation of dysfunctional proteins and intensified microglia reactivity in response, further amplifies neuroinflammation and neurotoxicity^13,15–18^. Depleting microglia chronically to attenuate their reactivity responses may mitigate neurodegenerative disease progression^19^ but could also exacerbate it^19,20^, while repopulation after ablation may improve outcomes^21^. Therefore, intervening in microglial dysfunction while maintaining viable populations of these cells may offer a mechanism to prevent neurodegeneration in early disease stages.

Despite the contributions of microglia to CNS disorder progression, neuropharmacology as a field has mostly focused on developing drugs which act on conventional neuronal targets such as ion channels and neurotransmitter receptors. We currently have limited approaches to specifically target microglia for therapy. Additionally, developing therapies that act only on local populations of dysfunctional microglia will likely be important for realizing new effective treatments for a variety of CNS disorders. However, targeting select neuroanatomically defined populations of microglia for therapy is non-trivial. CNS tissue barriers, including the blood brain barrier (BBB) and glia limitans, pose obstacles for delivering systematically administered drugs to the CNS^22^. Furthermore, since many CNS disorders involve tissue region specific pathology, widespread CNS delivery of therapies that overcome these tissue barriers may not be ideal for many disorders. intraparenchymal injection of viral vectors is currently routine in preclinical CNS models for inducing localized, cell specific expression of proteins or gene silencing, but there are numerous limitations with these technologies including immunogenicity, cytotoxicity, inability to target certain brain cells effectively, and restrictions on the types of therapies that can be delivered^23^. Microglia are particularly difficult to transfect *in vivo* via viral methods, and even viruses that have high transfection efficiency for microglia *in vitro* have poor performance *in vivo*^24^. Thus, there is a need to develop new, modular approaches for localized targeting of specific microglia populations to advance therapies for CNS disorders.

Non-viral nanocarriers, including polymeric nanoparticles and lipid micelles, may be a safer and more versatile approach for drug delivery compared to viral modalities. These systems can be designed for cell specific delivery through facile modification with multivalent targeting moieties presented on the nanocarrier exterior^25,26^. Small molecule ligands with high affinity for receptors and transporters that are highly expressed on the surface of particular cells in a tissue^26,27^ or at solid tumors^28–31^ have been incorporated into nanocarriers and prodrugs to improve targeting of therapy. Small molecule ligands already explored for this purpose include glucose (for Glut1 (*Slc2a1*))^32^, fructose (for Glut5 (*Slc2a5*))^33,34^, neutral amino acids – e.g. l-tyrosine, tryptophan (for Lat1 (*Slc7a5*))^35–37^, folic acid (for folate receptor (*Folr1*))^38^, Ascorbate (for Svct2 (*Slc23a2*))^35^, and for L-carnitine (for Octn2 (*Slc22a5*))^39^ . Dendrimers, a special type of highly branched macromolecule, are particularly well suited as nanocarriers for targeted delivery due to the potential to achieve high multivalent presentation of targeting ligands^40^. For example, folic acid decorated dendrimers improved cellular uptake and tissue retention of drug cargo in a mouse model of squamous cell carcinomas^41^. There has been recent interest in using dendrimers as nanocarriers to deliver therapies to the CNS. For example, polyamidoamine (PAMAM) dendrimers have been applied to the CNS as treatment carriers for traumatic brain injury, where uptake of dendrimers was not reliant on ligand interactions, but rather was a result of phagocytosis by cells in a reactive state due to their inflammation response^42^, and glioblastoma, where ligand presentation of CREKA allowed for targeted interaction with tumor vasculature ^42^. Histidine-maltose shell (G4HisMal) and PAMAM dendrimers have also been used in models of AD to modulate the synthesis of soluble amyloid beta and the formation of amyloid beta fibrils by recruiting the protein to the particle for microglial clearance^42,43^. Similarly, PAMAM dendrimers have been used in PD models to sequester alpha synuclein protein and prevent formation of lewy bodies in dopaminergic neurons, ionically interacting with the protein aggregates to disrupt large deposits and assist degradation^42^. However, targeted interactions with microglia via established dendrimer structures remains elusive, as use of PAMAM dendrimers functionalized with mannose receptors designed to target CD206 on microglia via systemic injection did not alter uptake or targeting^44^. Additionally, the use of mannose to target microglia^43,44^ facilitates BBB penetration which can be useful for systemic treatment but can also result in vascular clearance of the treatment resulting in poor efficiency of treatment. Despite the promise of targeted delivery, use of these strategies in localized intraparenchymal injections into the CNS is very limited, and most are used to target CNS vasculature and traverse the BBB following systemic administration. There is a need to develop nanocarriers that target microglia and can be applied locally for improved tissue specificity and treatment retention. This will require identifying viable cell surface targets that can provide specificity for microglia.

We hypothesized that cell surface proteins highly expressed on microglia could be leveraged for targeted delivery following local neural parenchyma injection of a ligand-decorated non-viral dendrimer carrier. In this work, we first performed an *in silico* analysis of neuron and glia specific transcriptomics datasets to identify unique and highly expressed surfaceome genes on the different cell types that may be amenable to therapeutic targeting. We confirmed expression of some targets in tissue by immunohistochemistry and in situ RNA hybridization. As part of proof-of-principle studies, we selected a cell surface transporter uniquely and highly expressed by microglia, *Slc2a5* (encoding Glut5 transporter protein), for nanocarrier targeting. To target this transporter, we synthesized dendrimers by thiol-ene and enzymatic catalyzed methods. We then employed polyionic complexation (PIC) of oppositely charged polyelectrolyte dendrimers to afford multivalent presentation of the transporter’s ligand, fructose, on the exterior of the nanocarrier. Neutral fructose functionalized PIC nanoparticles permitted fructose-dependent intracellular protein but not functional mRNA delivery to Glut5 expressing MCF-7 cells *in vitro*. Efficacy of neutral fructose-functionalized PIC nanoparticles was reduced with soluble fructose competition, and Fructose-PICs did not deliver protein cargo to cultured astrocytes that had negligible expression of Glut5. Fructose-PICs injected into the mouse striatum were well tolerated up to 1 mg/ml and were retained locally at the injection site. Neutral fructose functionalized nanoparticles permitted delivery of Cre protein to Glut5-expressing reactive glia in the context of an ischemic stroke injury and facilitated delivery to neurons in an equivalent manner to the protein solution in healthy tissue where there was no reactivity induced upregulation of Glut5. Our findings demonstrate the feasibility of using *in silico* analyses of RNA-seq datasets for identifying cell specific targets as well as the potential of fructose-functionalized, dendrimer-based PIC nanoparticles to selectively target therapies to Glut5 expressing cells.

## Results

### Neural cell type specific surfaceome targets can be identified by *in silico* analysis of RNA-Seq datasets

To identify neural cell type specific surfaceome targets we employed a bespoke *in silico* analysis of published RNA-seq datasets derived from bulk murine brain tissue (**Fig 1a**). We used datasets where RNA was collected specifically from brain neurons^45^, microglia^46^, and astrocytes^47^ by neural cell Cre-line based RiboTag immunoprecipitation methods as well as through the dataset of Zhang et al. that isolated the same three neural cells using a sequential immunopanning technique^48^. These bulk-sequenced, cell specific datasets were opted for here since they had sufficient read depth for identifying lowly expressed but unique transcripts that may otherwise be missed by current single cell methods. We normalized the raw counts derived from these different datasets by calculating FPKM values and computing z-scores, before using a curated *in silico* human surfaceome database^49^ to narrow the list of applicable genes for analysis to include only genes encoding for proteins expressed on the exterior surface of the cell membrane. We detected measurable expression of 1764 surfaceome genes across both the RiboTag and immunopanned data sets and a further 718 genes that were unique to RiboTag samples (**Fig 1d**). Principal Component Analysis (PCA) for dimensionality reduction of all surfaceome expressed genes, showed pronounced differences for the various neural cell types for both the RiboTag and immunopanned data (**Fig 1b,c**). Using a 2-dimensional PCA space, we quantified the gene expression specificity for the various neural cell types using a cosine similarity approach that involved taking the cosine of the angle between the gene factor loading vector and the cell type vector, setting a threshold for cell type specificity at a cosine similarity value of greater than or equal to 0.9. Using this approach we identified 374, 339, and 659 uniquely expressed surfaceome genes for astrocytes, microglia, and neurons respectively from the RiboTag dataset, while 267, 361, and 399 uniquely expressed surfaceome genes for the same cell types were identified from the immunopanned data. To further improve the robustness of the analysis, we condensed the gene lists by only including genes that were identified to be unique to a given cell type on both the RiboTag and immunopanned datasets, resulting in 133 genes unique to astrocytes, 188 genes to microglia, and 178 genes to neurons (**Fig 1d**).

**Figure 1:**
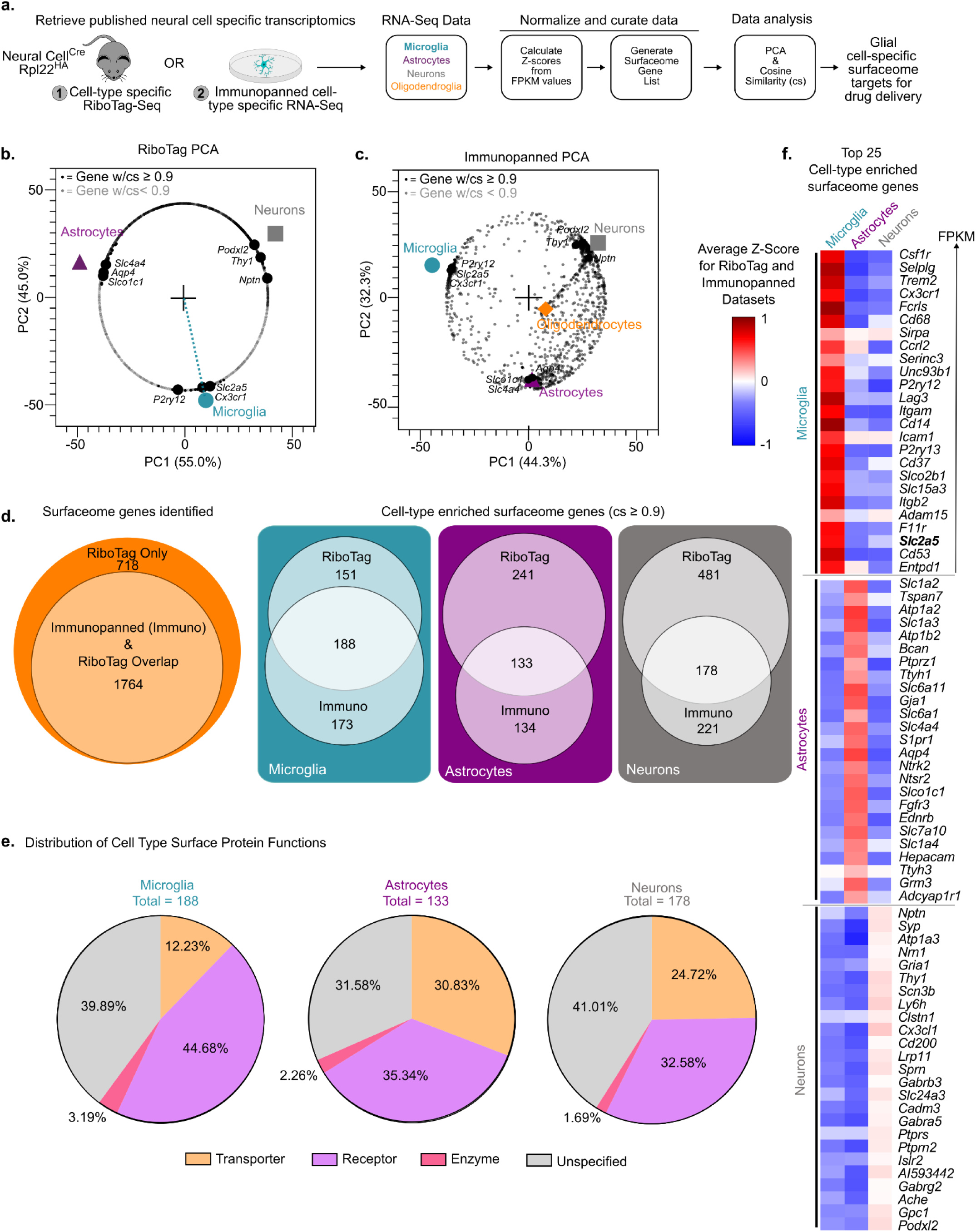
*In Silico* RNA-Seq Analysis of Neural Surfaceomes. (a) Work flow scheme for processing of published bulk tissue RiboTag and immunopanned RNA sequencing data from murine brain. (b) PCA of all surfaceome genes identified in the RiboTag dataset (grey), with genes that fell within the cell specificity cosine similarity threshold of 0.9 or higher shown in black. Indicators of microglia, astrocytes, and neurons are shown in the PCA space, with examples of three canonical genes that fell within our cell specificity threshold labeled for each cell type respectively. (c) PCA of surfaceome genes identified in the immunopanned dataset, similar to panel b. (d) Venn diagrams illustrating that all immunopanned dataset genes fell within the larger dataset of RiboTag genes, and showing the amount of overlap between the two datasets within the constraints of cell specificity threshold for microglia, astrocytes, and neurons. (e) Pie charts illustrating the protein functions associated with the cell specific surfaceome genes that occur in the overlap between the RiboTag and immunopanned datasets for microglia, astrocytes, and neurons. (f) Heat map of gene z-scores, showing twenty five genes with the highest FPKM scores for microglia, astrocytes, and neurons, providing an indicator of each gene’s z-score in all three cell types respectively.

Since we wanted to identify neural cell specific surfaceome targets for drug delivery, we curated the list of surfaceome genes by investigating the function of the encoded protein, prioritizing proteins with established ligands that could potentially be used for driving cell specific targeting interactions. We categorized the genes into transporter, receptor, enzyme, or unspecified functions, with receptors followed by transporters being the predominant known functional classifications across the three cell types (**Fig 1e**). We then sorted the receptor or transporter gene lists for each cell type based on high to low FPKM values, since highly expressed surfaceome proteins would likely improve target accessibility, and assembled the top twenty-five most highly expressed genes for each of the three cell types (**Fig 1f**). The top twenty-five genes for each neural cell type included many known canonical genes for that specific cell type, validating the analysis approach. For example, microglia showed high unique expression of: (i) *Csfr1,* a cytokine receptor that is critical for growth, survival, and function of microglia and which is accessed pharmacologically by small molecular inhibitors to deplete microglia^50^; (ii) *Trem2, a* transmembrane protein for anionic ligands that has a validated role in coordinating phagocytic processes and synaptic pruning^51^; and (iii) *P2ry12,* a purinergic G protein-coupled receptor for ADP which is established to be uniquely expressed by microglia^52^. Astrocytes showed uniquely high expression of its characteristic glutamate transporter *Slc1a2*, the aquaporin *Aq4*, the gap junction connexin *Gja1*, as well as *Slco1c1* which is a transporter that mediates uptake of sodium-independent thyroid hormone^53^. Neurons selectively expressed synapse-associated or synaptic regulatory genes *Thy1*, *Nptn*, and *Syp* as well as *Podxl2* which is a transmembrane receptor of sialic acid involved in regulating neurite branching and axon guidance^54^. Notably, comparing z-scores across the three cell types revealed that the unique surfaceome genes for microglia were some of the most highly expressed genes out of the entire population of expressed genes by microglia, whereas unique surfaceome genes for neurons were only at modest expression levels within the distribution of all neuron genes, suggesting that surfaceome genes may be more essential to overall microglia function than other neural cell types. Of particular note, *Slc2a5* was revealed to be uniquely and highly expressed by microglia and is a transporter that mediates uptake of fructose. For proof of concept, we selected *Slc2a5* for the nanoparticle-based targeting investigations explored throughout the rest of the work described here for several reasons including: (i) the ease of accessibility to its associated fructose ligand, (ii) its previous use for targeted delivery to other organs and tissues^33,34^; and (iii) since its unique expression in microglia was independently corroborated in multiple single cell RNA-seq databases (**Fig S1**).

### *Slc2a5* and encoded Glut5 are differentially expressed on microglia and astrocytes in mouse brain

To corroborate the *in silico* findings we next assessed the neural cell type specific expression of Glut5 in healthy and injured tissue, by performing immunohistochemistry (IHC) on murine brain tissue sections taken from healthy and ischemic stroke injured brains. Given limited available literature on Glut5 protein expression in the brain as assessed by IHC, we screened five unique Glut5 antibodies purchased from different reputable vendors. Surprisingly, the detected staining for Glut5 in healthy neural tissue varied widely across the five different antibodies. One antibody detected Glut5 expression around neuronal bodies, another colocalized with Glut1 expression on neurovasculature, another seemingly detected both of these features, and a fourth showed only diffuse non-specific staining (**Fig S3**). One antibody from Proteintech showed distinctly higher colocalization of Glut5 staining with Pu.1-positive microglia and Sox9-positive astrocytes compared to all the others (**Fig S3**). We further validated this particular antibody for detecting Glut5 by staining sections of mouse small intestine taken from the jejunum, a tissue region established to have a high expression of Glut5 along the apical brush-border membrane of the epithelium. We found that the Proteintech antibody readily detected Glut5 expression on the epithelium consistent with previous published studies^55,56^, whereas the others did not (**Fig S4**). Therefore, for the remainder of this study we chose to use the Proteintech antibody for Glut5 identification by IHC as well as western blotting protein analysis.

Glut5 expression was broadly and equally dispersed throughout the brain (**Fig 2a**), appearing as punctuated halos around Sox9-positive astrocytes and Pu.1-positive microglia at higher magnification (**Fig 2b**). Quantification of Glut5 expression in the striatum revealed that 70% of astrocytes had halos of Glut5-positive puncta around their Sox9-positive nuclei whereas 40% of microglia had similar patterns of expression around their Pu.1-positive nuclei, representing a significant reduction in the proportion of Glut5-positive cells between the two glial cell types (**Fig 2c**). Curious as to whether Glut5 expression was dependent on brain region, we compared Glut5 staining and glial cell colocalization in the cortex and hippocampus and found that the prevalence of Glut5 expression on astrocytes and microglia was conserved across all regions (**Fig S2**). We next evaluated Glut5 expression at ischemic stroke lesions induced via striatal injection of (L-N5-(1-iminoethyl)ornithine) (L-NIO), an established vasoconstrictive agent for inducing stroke, at two days after injury. During this acute stage of injury, L-NIO lesions are defined by distinct glial cell compartmentalization where a zone of migrating reactive microglia are separated from a forming astrocyte border and a more distal region of reactive but preserved neural tissue^57–60^ (**Fig 2d**). At two days post-stroke, there was a dramatic reduction in detection of Glut5 levels in the lesion core but globally preserved levels in regions of reactive neural tissue.

**Figure 2:**
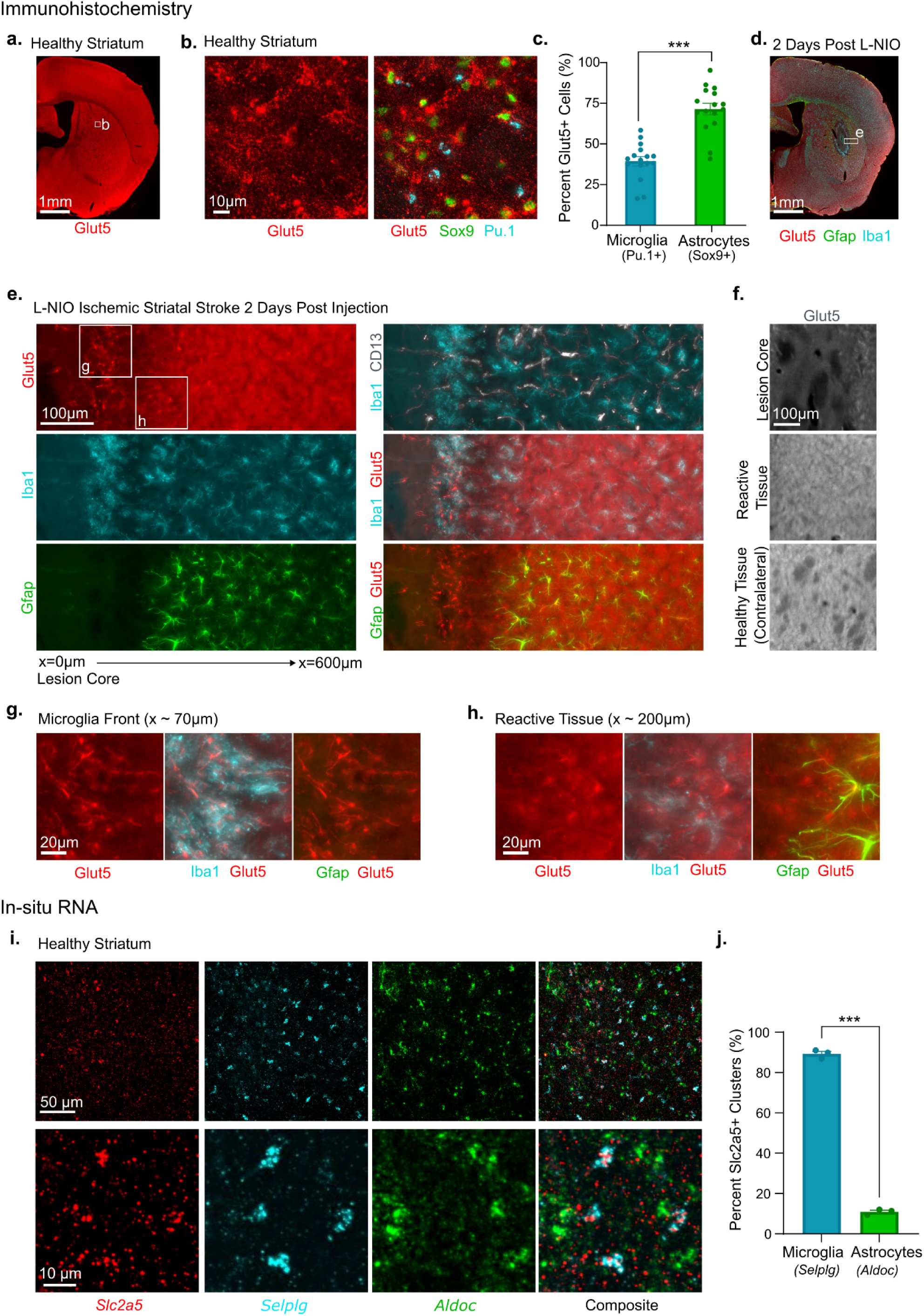
Slc2a5 and encoded Glut5 protein are differentially expressed on microglia and astrocytes in healthy and injured CNS. (a) Survey image of Glut5 stained in healthy mouse striatum. (b) Detailed image showing Glut5 (red) expression in healthy mouse striatum costained with Pu.1 (cyan) a nuclear marker for microglia and Sox9 (green) as a nuclear marker for astrocytes. (c) Quantification of the percentage of Pu1 positive cells and Sox9 positive cells that colocalize with high Glut5 staining intensity in healthy striatum tissue. (d) Survey image of Glut5, Gfap, and Iba1 stained in mouse striatum 2 days post L-NIO induced ischemic stroke. (e) 20x images of Glut5 (red), Iba1 (cyan), Gfap (green), and CD13 (white), across a gradient of reactivity induced by striatal injection of LNIO, where the left boundary of the image is showing the lesion core and the right boundary of the image shows the less affected tissue. (f) Glut5 staining across three different stages of tissue injury, the lesion core of the ischemic stroke, the reactive tissue adjacent to the stroke border, and healthy tissue on the contralateral side of the section. (g) Cropped 20x images of Glut5 (red), Iba1 (cyan), and Gfap (green) at the reactive microglia border, approximately 73µm from the left boundary of the image in panel d. (h) Cropped 20x images of Glut5 (red), Iba1 (cyan), and Gfap (green) at the reactive astrocyte region, approximately 255µm from the left boundary of the image in panel d. (i) Fluorescent in situ RNA probes of Slc2a5 (red), canonical microglia gene Selplg (cyan), and canonical astrocyte gene Aldoc (green) in healthy mouse striatum. (j) Quantification of colocalization between Slc2a5 and Selplg or Aldoc, demonstrating high occurrence on microglia compared to astrocytes. Quantifications, panels c and h, show mean +/- SEM (standard error of mean) and were analyzed with unpaired t-tests, ***p-value <0.001.

Both the reactive microglial front immediately adjacent to the L-NIO lesion core and the forming astrocyte border showed increased expression of Glut5 on individual cells at 2 days post-stroke compared to healthy tissue (**Fig 2e-h**), whereas the adjacent reactive tissue showed Glut5 expression restricted to Gfap-positive astrocytes. The noted increased expression of Glut5 in reactive glia is in agreement with previous studies showing upregulation of *Slc2a5* transcripts in both microglia and astrocytes in states of injury and neuroinflammation ^61^.

The detection of Glut5 expression in healthy astrocytes was contradictory to both the RNA-seq data from the *in silico* analysis as well as prior single cell sequencing results^62–71^, so we decided to incorporate RNA *in situ* hybridization to compare protein versus RNA expression levels of *Slc2a5*/Glut5 in healthy brain tissue. We found that clusters of fluorescence from *Slc2a5* probes had a significantly high colocalization with microglia-specific *Selplg* probe expression but not astrocyte-specific *Aldoc* probes. Quantification revealed that 85% of microglia expressed *Slc2a5* whereas 10% of astrocytes expressed *Slc2a5* (**Fig 2i,j**). By contrast, detection of an astrocyte specific transporter identified in our *in silico* analysis, *Slc1a2*, was restricted to astrocytes with no expression noted in microglia or *Snap25/Atp1b1*-positive neurons. Similarly, one of the neuron specific genes identified in our *in silico* analysis, *Syp*, which encodes for synaptic vesicle proteins^72,73^, was highly colocalized with *Snap25/Atp1b1* positive neurons, but not with either of the glial cell probes (**Fig S5**). Thus, these data show that the RNA in situ hybridization results were congruent with the *in silico* RNA-seq findings, confirming that *Slc2a5* (Glut5) is a microglia specific target. However, our data also show that protein expression of Glut5 in both healthy and injured tissue may occur in both microglia and astrocytes. This RNA and protein expression discrepancy could be technical such as due to unanticipated antibody cross-reactivity with other similar Glut family transporters or perhaps biological such as a result of exosome mediated transfer between these different glial cell types^74^. Irrespective of the source of this discrepancy, the capacity to potentially target microglia and astrocytes via Glut5 was still appealing, so we moved forward with developing a non-viral nanoparticle formulation that can incorporate fructose as a ligand to target these Glut5-expressing cells.

### Synthesis of fructose functionalized dendrimers

Dendrimers are highly branched macromolecules that allow for multivalent presentation of specific terminal functional groups^40,75,76^. We devised a dendrimer formulation that would allow for the multivalent presentation of fructose to target Glut5 to increase the potential for drug carrier engagement with Glut5 expressing cells such as microglia in the CNS. We synthesized a dendrimer scaffold using a series of sequential thiol-ene radical and enzymatically catalyzed reactions under solvent-free conditions (**Fig 3a**). Starting from a core composed of pentaerythritol tetrakis (3-mercaptopropinate) (PTE3MP), each successive generation of dendrimer was synthesized via two sequential reactions. The first reaction involved a solvent free, UV radical initiated reaction between thiol groups on the PTE3MP and the vinyl group on trimethylolpropane allyl ether (TMPAE). This reaction proceeded to full completion and doubled the number of branches on the molecule, as TMPAE contains two hydroxyl groups on which to add further to in the subsequent reaction. The second reaction involved an enzymatically catalyzed transesterification of these two free primary hydroxyls with excess ethyl 3-mercaptopropionate (E3MP), catalyzed by Candida antarctica Lipase B (CALB)-immobilized on acrylic resin, that re-introduced thiol groups onto the end of each dendrimer branch for further generation growth or end cap functionalization via thiol-ene reactions. Each dendrimer generation was purified by precipitating and washing in hexanes repeatedly until thin layer chromatography (TLC) indicated no remaining unreacted E3MP was present in the resultant product pellet. Trace amounts of solvent were removed from the dendrimer product by rotary evaporation before proceeding to synthesize the next generation. We grew dendrimer scaffolds from a G1 with a maximum of 8 branches to a G5 dendrimer with a maximum of 128 branches. Reaction progress was monitored and confirmed by TLC and proton NMR whereby consumption of the allyl group from the TMPAE (5.2, 5.8ppm), the presence of hydroxyl protons (4.2ppm) in G1a, and subsequent loss of these detectable hydroxyl protons as well as positive Ellman’s TLC indicated successful G1 synthesis (**Fig 3b, Fig S6**). Essentially all proton chemical shifts were conserved across dendrimer generations from G1-G5 but peaks broadened with increasing generations, reflecting greater conformational heterogeneity with increasing dendrimer size (**Fig3b, Fig S6**). Increasing dendrimer size across generations was also confirmed by cellulose TLC (**Fig S7**).

**Figure 3:**
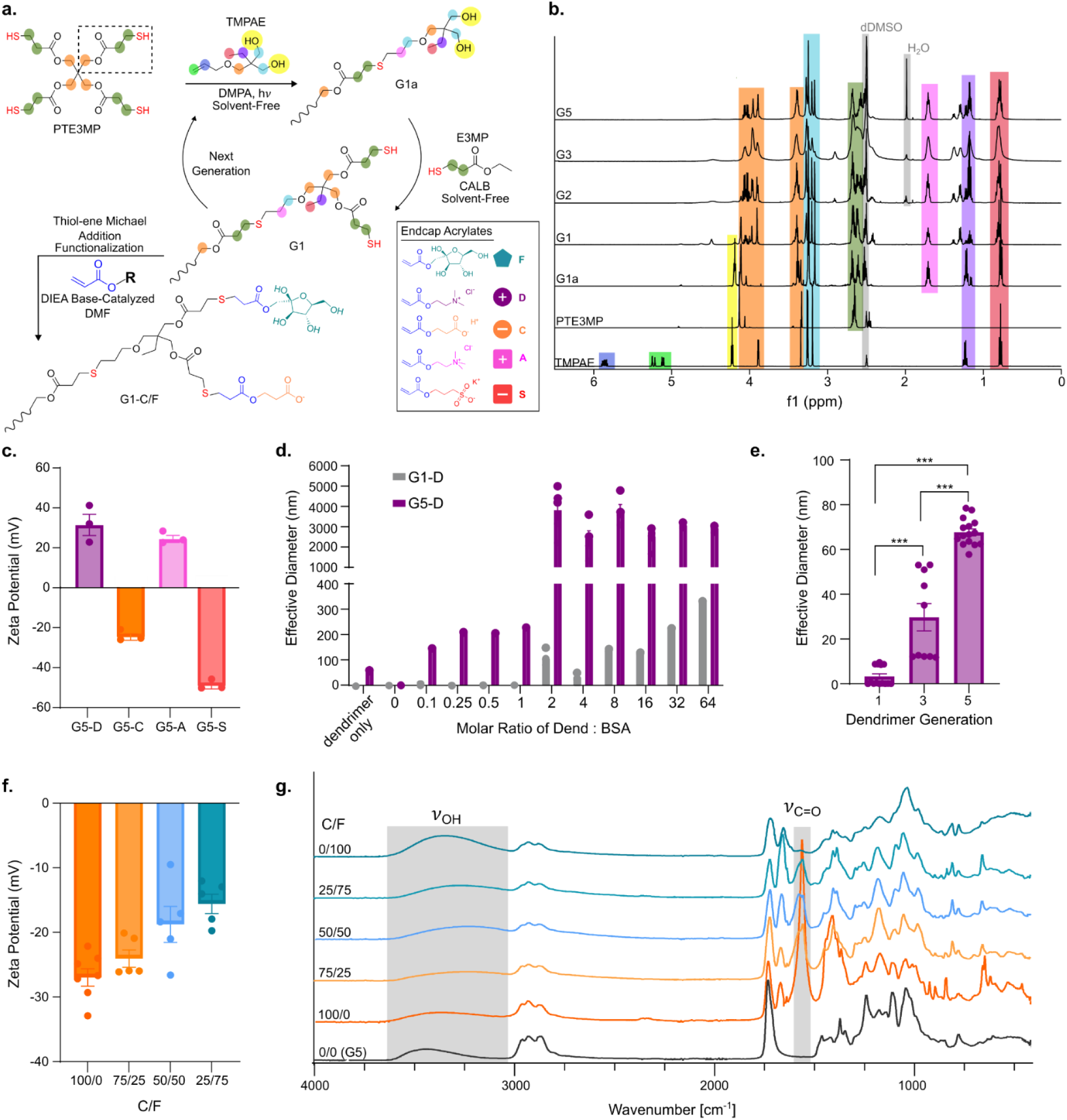
Synthesis and characterization of dendrimers. (a) Reaction scheme for dendrimer scaffold synthesis and endcap functionalization shown for a single branch of a G1 dendrimer with protons highlighted to correspond to the highlighted shifts in panel b. (b) Proton NMR spectra of dendrimer scaffold reactants and three different generations of synthesized scaffolds (G1,G2,G3) with shifts highlighted to correspond with highlighted protons on the structure drawing in a, all spectra normalized to the CH3 shift occurring at 0.83 ppm highlighted in red. (c) Zeta potential measurements of charge of G5 dendrimers functionalized with either ionizable cationic D endcap, ionizable anionic C endcap, constitutively charged cationic A endcap, or constitutively charged anionic S endcap, demonstrating success in altering molecule surface charge via endcap decoration. (d) Increase in effective diameter as measured with DLS as G1 and G5 D functionalized dendrimers complex a negatively charged BSA protein, demonstrating that complexation occurs at a lower molar ratio of dendrimer to protein when the dendrimer is of a larger generation. (e) Effective diameter of D functionalized dendrimers of three different generations in solution in PBS at 10mg/mL. (f) Zeta potential measurements of charge of G5 dendrimers functionalized with different ratios of anionic C endcap and fructose F endcap in solution in PBS at 10mg/mL. (g) FTIR spectra of G5 dendrimers functionalized with different ratios of anionic C endcap and fructose F endcap, showing correlation between the stoichiometric ratio of endcap added and the area under the curve of the corresponding FTIR stretch. Quantifications in panels c,d,e,f,h,i are mean +/-SEM (standard error of mean). Panel e was analyzed with one way ANOVA and Tukey’s test, ***p-value <0.001.

We next modified the branch ending thiol groups on dendrimer scaffolds with various mono-acrylate functionalized groups via a based catalyzed Michael addition reaction to terminally endcap the dendrimer. We demonstrated successful dendrimer endcapping with five different functional groups: (i) constitutively cationically charged quaternary amine groups using [2-(Acryloyloxy)ethyl] trimethylammonium chloride (A), (ii) constitutively anionically charged sulfate groups using 3-Sulfopropyl acrylate (S), (iii) ionizable cationically charged tertiary amine groups with 2-(Dimethylamino)ethyl acrylate (D), (iv) ionizable anionic groups with 2-Carboxyethyl acrylate (C), which were all previously used by our group in the development of ionically complexed trehalose coacervates^77^, and (v) fructose acrylate (F) that was synthesized in-house using enzymatic catalyzed reactions (**Fig S8**). Successful dendrimer functionalization was determined through chemical characterization by FTIR (**Fig S9**) and NMR (**Fig S10,S11**), as well as physical characterization by zeta potential measurements, which demonstrated that A and D end-capped dendrimers had positive zeta potentials while A and S endcapped dendrimers had negative zeta potentials, consistent with chemical characteristics of the end-capping groups (**Fig 3c**).

Due to the nature of the repeating structure present across all generations and branches of dendrimer molecules, it was not possible to readily determine and quantify an absolute number of total branches by NMR. Instead, we evaluated complexation capacity of cationic D endcapped dendrimers with anionic bovine serum albumin (BSA, pI = 4.5–5.0) under physiological pH conditions to profile multivalency across different dendrimer generations. D endcapped dendrimers across generations had equivalent pkAs by fluorescent 6-(p-toluidino)-2-naphthalenesulfonic acid (TNS) assay, indicating that any complexation differences would not be due to differences in relative ionization, but in total number of present endcaps. G5-D dendrimers, likely due to the greater number of branches and a higher multivalency presentation of cationic charges compared to G1-D dendrimers, formed 100nm-plus complexes at a 0.1 molar ratio of G5-D to BSA, requiring a 20-fold lower molar concentration of dendrimer material to complex BSA than G1-D dendrimers (**Fig 3d, FigS7**). As another way to assess increased branching and size in higher generation dendrimers, we used Matrix-Assisted Laser Desorption/Ionization Time-of-Flight (MALDI-TOF) mass spectrometry to evaluate the smallest (G1) and largest (G5) dendrimer generations synthesized. G5 dendrimers not only had a greater relative abundance of higher molecular weight fragments compared to the G1, but also generated more ionized fragments, consistent with greater multivalency (**Fig S7**). Finally, we detected increasing size of dendrimers in aqueous solution as a function of generation by dynamic light scattering (DLS), with D1, D3 and D5 dendrimers having a mean effective diameter of 4nm, 30nm, and 68nm respectively, representing a near 17-fold increase in size as dendrimers were grown from G1 to G5 (**Fig 3e**).

Since fully fructose (F) functionalized dendrimers were sparingly soluble in aqueous buffers, we synthesized partially fructose functionalized dendrimers and added proportions of ionizable groups to improve dendrimer solubility, permit electrostatic interactions with charged biomolecular drugs, or facilitate polyionic complex (PIC) formation through the mixing of oppositely charged dendrimers. We employed the highest generation dendrimer scaffold synthesized, G5, for the remainder of the experiments since it affords the greatest multivalency. Varying the stoichiometric ratio of ionizable C acrylate to F acrylate within the endcapping Micahel addition reaction, yielded functionalized dendrimers with essentially the same equivalent proportion of endcap species resulting in facile modulation of total charge (**Fig 3f**) and fructose content (**Fig 3g,h**, **Fig S11**) on the dendrimer as assessed by Zeta potential, NMR, and FTIR. All synthesized C/F functionalized dendrimers were water soluble at concentrations up to 10mg/ml (**Fig S12**). Initially, we also attempted to synthesize dendrimers using stoichiometric mixtures of D and F acrylate endcaps to incorporate cationic charge and fructose ligands into a single dendrimer, which would be particularly relevant for nucleic acid delivery. However, we detected a likely D endcap induced base-catalyzed degradation of the fructose on the dendrimers prepared under these synthesis conditions indicated by a white to dark brown change in the dendrimer during lyophilization and complete loss of aqueous solubility upon attempted reconstitution in water or buffered solutions (**Fig S12**). Consequently, since we could only prepare anionic dendrimers functionalized with fructose, we pursued combining these dendrimers with a fully D functionalized dendrimer to form a PIC as the most appropriate way to prepare a versatile fructose-based nanocarrier. We envisioned this approach would allow us to still load dendrimers with nucleic acid or any other anionically charged cargo and improve prospects for endosomal escape upon delivery of these carriers into cells.

### Mixing of oppositely charged dendrimers forms polyionic complexes (PICs)

Polyionic complexes (PICs) are formulated by combining oppositely charged polyelectrolytes that self-assemble into larger complexes due to electrostatic interactions and can be used to load and deliver drug cargo^78^. To test whether mixing oppositely charged dendrimers formed PICs, we started by combining anionic dendrimers with and without fructose (C, S, or C/F) and cationic dendrimers (D and A) under physiologically buffered conditions. While all dendrimers had excellent aqueous solubility in solution alone (**Fig S13**), upon mixing two oppositely charged dendrimers under the same buffered conditions at 10 mg/ml, we observed rapid liquid-liquid phase separation that resulted in significantly increased turbidity and formation of discrete, suspended PIC droplets (**Fig S13**). PICs were formed with both ionizable (C+D) and constitutively charged (A+S) chemistries at 10 mg/ml with no apparent differences in concentration or size (**Fig S13**). The size of PICs could however be readily tuned by modulating the formulation concentration. For example, while PICs prepared at 10 mg/ml generated microscopic droplets visible under brightfield microscopy, preparations made at lower concentrations (0.1 mg/ml) generated nanosized complexes detectable only by DLS (**Fig 4a, S13**).

**Figure 4:**
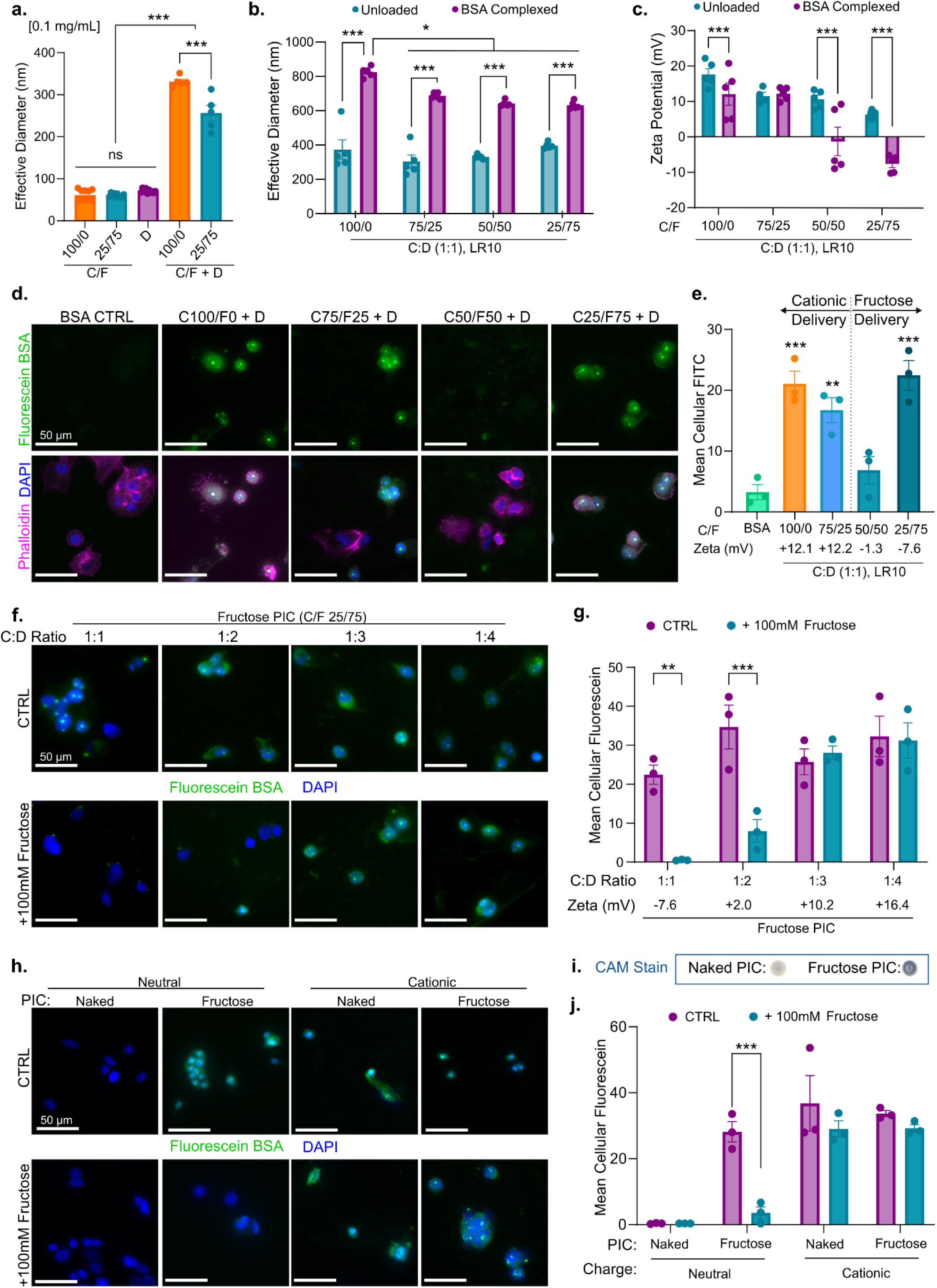
PIC facilitated protein delivery *in vitro* to MCF-7 cells. (a) Effective diameter measured with DLS at 0.1mg/ml demonstrating that PICs are significantly larger in size than dendrimers of a single charge alone. Quantifications, panels c and e, show mean +/- SEM (standard error of mean) and were analyzed with one way ANOVA and Tukey’s test, ***p-value <0.001. (b) DLS of the effective diameter of PICs at 0.1mg/ml in 1X PBS, with and without BSA protein loaded, n=5. (c) Zeta potential of PICs with varying amounts of fructose presentation, with and without BSA protein loaded, all at a loading ratio of 10 and PIC concentration of 10mg/ml in 1X PBS, n=5. All cell culture experiments in this figure (panels d-h,j) were MCF-7 cells treated with PICs at 0.1mg/mL, LR 10 of Fluoroscein BSA protein. (d) Representative images of cytochemical staining of PIC treatments at 1:1 for C:D ratio, with varying levels of fructose presentation, in serum free media for 24 hours. (e) Quantification of fluorescein signal measured within phalloidin stained cell boundaries for the groups in panel d, along with the zeta potential measurement of that PIC after protein is loaded, n=3. (f) Representative images of cytochemical staining of Fructose-PICs (C25/F75 + D) treatment across anionic:cationic C:D dendrimer ratios, with and without competition of 100mM free fructose media. (g) Quantification of fluorescein signal measured within phalloidin stained cell boundaries for the groups in panel f, along with the zeta potential measurement of that PIC after protein is loaded, n=3. (h) Representative images of cytochemical staining of treatment with PICs (C100/F0 + D) and Fructose-PICs (C25/F75 + D), that are either neutral (0-2mV) or cationic (12-15mV) in charge, with and without competition of 100mM free fructose media. (i) CAM stain of formulated Fructose-PIC and PIC, demonstrating fructose presentation is maintained after formulation for treatment. (j) Quantification of fluorescein signal measured within phalloidin stained cell boundaries for the groups in panel h, n=3. Quantifications, panels b, c, e, g, j, show mean +/- SEM (standard error of mean) and b, c, g, j, were analyzed with two way ANOVA and Tukey’s test, ***p-value <0.001.

Notably, even at dilute concentrations, formulated PIC sizes ranged from 200nm to 450 nm in diameter, which represented an approximate 5-fold increase in size compared to dendrimer only solutions, indicating strong complexation between oppositely charged dendrimers (**Fig 4a**). PIC formulations prepared using fructose functionalized anionic dendrimers were significantly smaller than those prepared with anionic dendrimers without fructose (**Fig 4a**). These data are consistent with reduced numbers of dendrimer molecules aggregating together with the fructose functionalized dendrimers, since they had an overall reduced charge on the molecule (**Fig 3f**).

To characterize the loading of model biomacromolecule drug cargo into dendrimer-based PICs, we first used a fluorescein conjugated bovine serum albumin (F-BSA) for initial proof-of-concept studies. We measured quenching of detected fluorescein intensity as an indicator of dendrimer-protein complexation. Irrespective of formulation, fluorescence intensity decreased as the mass loading ratio (LR) of dendrimer relative to protein increased, indicating enhanced interactions between the dendrimer PICs and the protein and greater cargo loading (**Fig S14**). We tested protein-dendrimer complexation with two other FITC-labeled proteins to assess robustness of the PIC cargo loading capacity, namely cytochrome C (pI = 10-10.8) which has a net cationic charge at pH=7.4, and FITC-labeled Human IgG (pI = 6.5–9.5) which has a relatively neutral charge under physiological conditions. We found that the negatively charged F-BSA cargo complexed more efficiently into PICs than either of the other two proteins, as both the cytochrome C and IgG required higher loading ratios of dendrimer to protein for fluorescence quenching to be detected. Despite this, formulating PICs with cytochrome C or IgG resulted in increased PIC size suggesting protein loading was readily achieved (**Fig S14**). Using F-BSA as an efficiently complexed model protein cargo, we formulated four PICs for initial size characterization: three different fructose compositions (using 25/75, 50/50, 75/25 C/F functionalized anionic dendrimers complexed with D functionalized cationic dendrimer), and one non-fructose containing naked-PIC (100/0 C/F functionalized anionic dendrimers complexed with D functionalized cationic dendrimer). Upon addition of F-BSA, all four PIC formulations showed increased effective diameters, ranging from 600-800nm when prepared at 0.1mg/ml (**Fig 4b**). Similarly to the sizing results of unloaded PICs, the fructose functionalized PICs loaded with F-BSA generated significantly smaller complexes than the naked-PIC without fructose, not exceeding 710nm in diameter (**Fig 4b**). Even though prepared at calculated stoichiometric ratios of 1:1 C:D end cap, all PICs showed a net cationic charge prior to protein loading, likely due to a higher efficiency in the functionalization of the dendrimer with D endcaps compared to C endcaps for the Michael addition reaction.

Specifically, when reacted for the same amount of time and under the same conditions, the D endcap reaction reaches higher completion than the C endcap reaction (**Fig 4c, S10**). When loaded with the negatively charged BSA cargo, the overall charge on the PIC remained cationic for low fructose or naked formulations, but was neutral or slightly net negative for formulations containing greater than or equal to 50% conjugated fructose on the anionic dendrimer (**Fig 4c**).

For initial *in vitro* cytocompatibility and intracellular protein delivery testing, we evaluated PIC formulations on an MCF-7 human breast cancer cell line, since it highly expresses the Glut5 fructose transporter^79^ and is commercially available^80^. We first tested PICs without any loaded cargo and identified two main formulation parameters that negatively affected MCF-7 viability: (i) a significant charge excess in either the cationic or anionic direction, or (ii) PIC concentrations substantially greater than 1 mg/ml. The concentration dependent cytotoxicity correlated with larger diameter macroscopic droplets, whereas lower concentration cytocompatible PICs were sized in the range of 100-300 nm (**Fig S15**). Based on these insights, we restricted intracellular delivery evaluations to PIC formulations prepared at 1:1 C:D ratios and at concentrations ≤1 mg/ml (**Fig S15**). Notably, increasing the amount of fructose conjugated onto the anionic dendrimer as well as complexing PICs with protein cargo improved cytocompatibility, likely due to further cationic charge shielding (**Fig S15**).

### Fructose-PICs permit fructose dependent intracellular protein delivery

We next tested intracellular delivery of F-BSA to MCF-7 cells using PICs prepared from anionic dendrimers with different extents of fructose conjugation and benchmarked them against a no-fructose containing naked-PIC (100/0 C/F) and an F-BSA solution condition as control groups. After 24 hours of treatment, the F-BSA solution control showed negligible delivery of protein to the cytoplasm of cultured MCF-7 cells consistent with previous studies^77,81^. Naked-PICs showed high intracellular delivery of F-BSA protein that was comparable to both the lowest and highest fructose-PICs (75/25 C/F and 25/75 C/F) (**Fig 4d,e**). By contrast, the intermediately fructose functionalized PIC (50/50 C/F) showed minimal F-BSA intracellular delivery (**Fig 4d,e**). Based on these results and factoring in the net surface charge on each of these PIC formulations as determined by Zeta potential (**Fig 4c**), we hypothesized that there were two main factors influencing intracellular delivery: (1) net cationic charge driven delivery, consistent with prior findings for similar non-viral lipid and polymer-based formulations^82,83^, and (2) fructose mediated delivery. High cationic surface charge likely enabled effective protein delivery for the naked and the low fructose-PICs whereas fructose mediated delivery could explain the elevated intracellular F-BSA levels seen for the neutral, high fructose-PIC formulation. The PIC formulation made using 50/50 C/F had limited F-BSA delivery, potentially because it was insufficiently equipped to enable either of the two delivery modes.

To further probe these potential intracellular delivery mechanisms, we focused next on testing fructose-PIC formulations prepared using only one of the fructose anionic dendrimers (25C/75F), since it showed the best delivery outcomes in the prior experiment and contained the greatest amount of conjugated fructose. To confirm the fructose dependent mechanism of delivery, we modulated the net charge presented by the fructose-PIC when formulated with F-BSA by increasing the relative proportion of cationic dendrimer from 1:1 to 1:4 C:D (anionic:cationic) and testing the differentially charged fructose-PIC formulations in media prepared with and without soluble fructose (100mM) to competitively inhibit the Glut5 transporter. If intracellular delivery directed by fructose-PICs is mediated through direct fructose and Glut5 interactions, competition with free soluble fructose should readily block F-BSA from being delivered into the cell. Indeed, for fructose-PICs with a net-negative or near neutral charge, competition with soluble fructose dramatically attenuated the intracellular delivery of F-BSA (**Fig 4f,g**). By contrast, F-BSA delivery permitted by the highly cationic fructose-PICs was unperturbed by the soluble fructose competition, consistent with a cationic charge mediated mechanism of delivery (**Fig 4f,g**). To strengthen our confidence in these findings, we tuned the anionic:cationic dendrimer ratio for both naked-PICs and fructose-PICs such that we established formulations that were either near neutral (0-2mV) or highly cationic (12-15mV) (**Fig 4i-j, S15**). For the near neutral formulations, fructose-PICs but not naked-PICs enabled F-BSA delivery into MCF-7 cells, and delivery was readily blocked by competition with soluble fructose. By contrast, both naked-PICs and fructose-PICs when prepared with high net cationic charge effectively delivered F-BSA in a manner that was unperturbed by fructose competition. Thus, taken together these results show that fructose-PICs, when formulated with negative or near neutral surface charge permit fructose-mediated intracellular delivery of protein cargo to cultured Glut5 expressing cells *in vitro*.

To test the specificity of fructose-PIC based delivery, we next evaluated protein delivery outcomes when Fructose-PICs were applied to cells that do not express high levels of Glut5. Mouse neural progenitor cell derived astrocytes that we have used previously^77,84^ express high levels of the glutamate transporter *Scl1a2* but do not express *Slc2a5* transcripts, so we considered these cells to be an appropriate candidate for probing delivery specificity (**Fig 5a**). We confirmed minimal protein expression of the encoded Glut5 transporter in mouse astrocyte lysates, with it being at least 3.5-fold less abundant than in MCF-7 cells (**Fig 5b**). To evaluate intracellular delivery in a higher throughput manner, we used an automated fluorescence cell counter to measure the relative intensity of F-BSA across individual cells (**Fig 5c**), as well as the proportion of cells within the population that had intracellular F-BSA levels above a background threshold (**Fig 5d**), following PIC treatment in both MCF-7 cells and astrocytes. This evaluation method enabled improved detection of intracellular delivery since cells were washed and trypsinized prior to reading and increased the number of evaluated cells 30-fold compared to the prior immunocytochemistry approach. Like before, neutral Fructose-PICs facilitated excellent F-BSA delivery to MCF-7 cells such that an average of 67% of cells had detected F-BSA delivery when treated with 1 mg/ml Fructose-PICs and delivery was abolished with soluble fructose competition (**Fig 5c,d**). By comparison, neutral naked-PICs showed no delivery of F-BSA to MCF-7 cells as before. Most notably, both neutral fructose-PICs and naked-PICs failed to deliver F-BSA cargo to cultured astrocytes consistent with fructose mediated delivery mechanisms being absent in these cells (**Fig 5c,d**).

**Figure 5:**
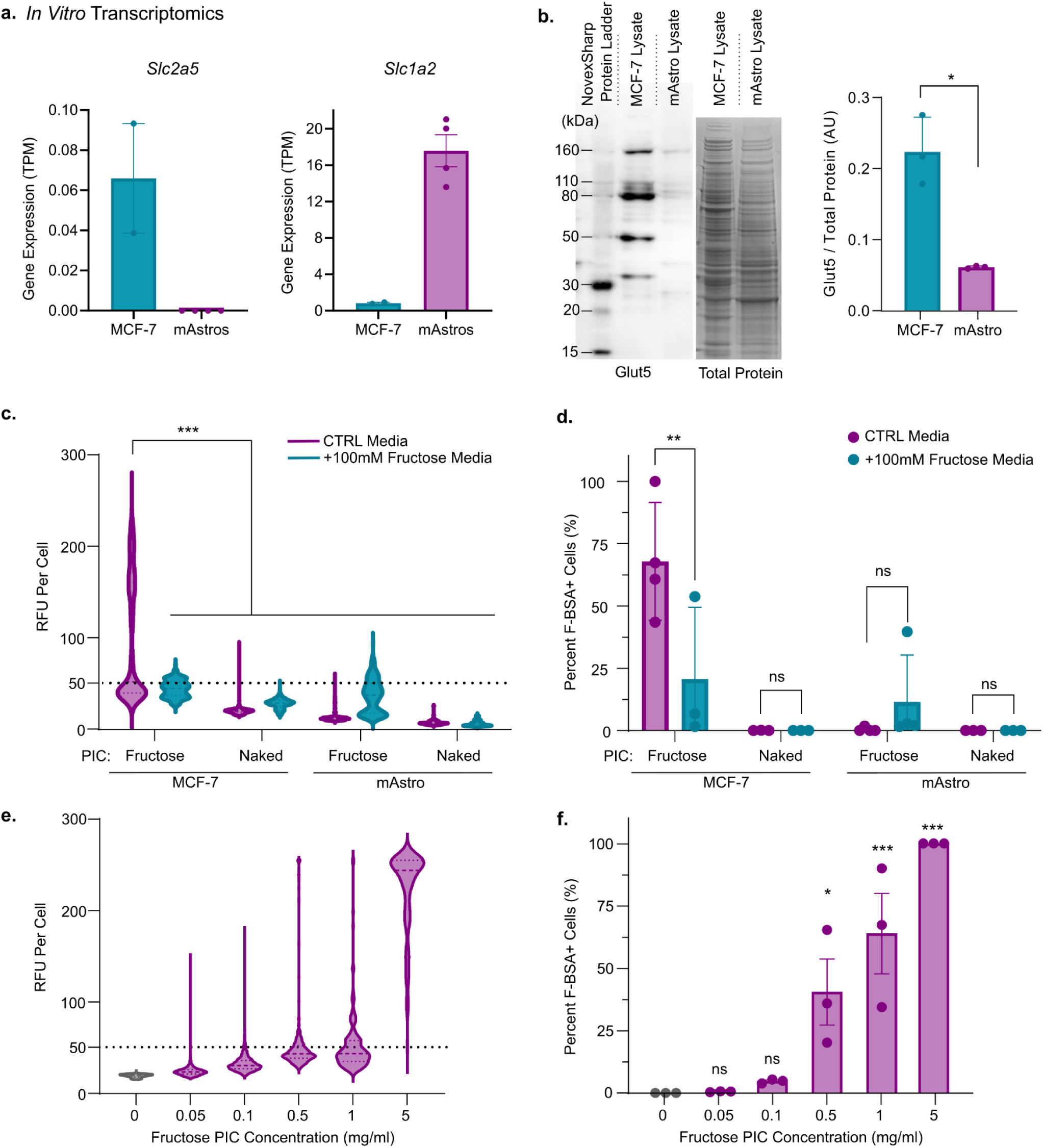
High throughput delivery analysis demonstrated on Glut5 expressing MCF-7 cells, and non-Glut5 expressing derived astrocytes. (a) In vitro transcriptomics from either published MCF-7 data or transcriptomics of our mouse NPC-derived astrocytes differentiated with 100ng/mL CNTF and 2% FBS for 48 hours, showing *Slc2a5* fructose transporter gene and *Slc1a2* canonical astrocyte glutamate transporter gene. (b) Western blot of MCF-7 and mAstro cell lysate, stained with ProteinTech Glut5 antibody and Coomassie G-250 total protein stain, with quantification of the 80kDa band intensity normalized to total protein, n=3, unpaired t-test comparison. All cell culture experiments in this figure were done with PICs of a neutral charge, complexed with Fluorescein BSA at LR 10, all RFU measures were done with a 488nm light filter on the Countess for high throughput quantification. (c) RFU distribution in both MCF-7 and mAstro cells treated with PICs and Fructose-PICs, with and without competition of 100mM free fructose media. (d) Percentage of cells with an RFU above 50, qualifying them to be considered positive for intracellular delivery, same sample groups as panel d, n=3. Panels c-d were analyzed with two way ANOVA and Tukey’s test. (e) RFU distribution of MCF-7 cells treated with Fructose-PICs in control media over increasing concentration. (f) Percentage of cells with an RFU above 50, qualifying them to be considered positive for intracellular delivery, same sample groups as panel e, n=3, analyzed with one-way ANOVA, with Dunnett’s multiple comparisons test compared against the 0mg/ml control group. All quantifications show mean +/-SEM (standard error of mean), *p-value <0.05, **p-value <0.01, ***p-value <0.001.

Delivery permitted to astrocytes by both carriers was unaltered by the presence of soluble fructose in the culture media and there was no change to Glut5 expression in astrocytes under these conditions (**Fig S16)**. An evaluation of dose response for Fructose-PICs on MCF-7 cells also revealed a positive correlation between the Fructose-PIC concentration and the total F-BSA delivered into each cell as well as the proportion of cells with detectable F-BSA. Essentially 100% of cells showed high F-BSA levels when treated with fructose-PICs at 5 mg/ml, whereas only 40% of cells had any detectable F-BSA at 0.5mg/ml (**Fig 5e,f**).

Overall, these data demonstrate that effective intracellular delivery by neutral fructose-PICs correlates strongly with Glut5 transporter expression such that cells with minimal Glut5 do not take up the fructose-PICs or its cargo, supporting the idea that fructose-PICs enable highly effective targeted delivery to Glut5 expressing cells.

### Fructose-PICs permit intracellular delivery of functional proteins but not mRNA

To determine if fructose-PICs can deliver functional protein cargo intracellularly, we loaded an engineered fusion protein of Cre recombinase covalently linked to a negatively supercharged Green Fluorescent Protein (GFP) with a theoretical net charge of -27 (hencefore referred to as (–27)GFP-Cre) and treated Glut5 expressing tail tip fibroblasts derived from Ai14 Cre reporter mice. These Ai14 fibroblasts contain a Lox-Stop-Lox-tdTomato construct under a ubiquitous CAG promoter such that Cre-Lox recombination and resultant tdTomato reporter expression occur in cells receiving functional Cre protein delivery^85^. The highly negative (–27)GFP-Cre enabled efficient loading into net neutral Fructose-PICs using a dendrimer to protein mass loading ratio (LR) of 10. The (–27)GFP-Cre loaded Fructose-PICs had a mean diameter of 430nm which meant complexes were 43% larger than those formed with F-BSA at the same 1mg/ml concentration (**Fig 6a**). As the LR used to prepare the (–27)GFP-Cre loaded Cre Fructose-PICs increased, the size of the complexes decreased such that a LR of 100 showed 300nm complexes (**Fig S17**). While multiple loading ratios of (–27)GFP-Cre loaded Fructose-PICs produced nanoscale complexes, a mass loading ratio of 10 was sufficient to form a complex that was neutrally charged (**Fig 6b**). To assess functional delivery *in vitro,* we applied a series of (–27)GFP-Cre loaded neutral fructose-PICs with LR of 10, 50 and 100 onto floxed Ai14 fibroblasts at 1mg/ml and evaluated cells for tdTomato expression 4 days post treatment (**Fig 6c-e**). These primary fibroblasts expressed the Glut5 transporter across essentially all cells and at comparably high levels to that detected in MCF-7 cells. While treating fibroblasts with the (–27)GFP-Cre protein only control promoted negligible tdTomato expression, all three Fructose-PIC formulations mediated improved functional (–27)GFP-Cre delivery, with the LR of 10 resulting in the most effective delivery with tdTomato expression detected in approximately 31% of cells (**Fig 6d,e**). While the LR of 10 formulation had the highest functional protein delivery, we also found that the LR of 100 had the next highest total delivery of approximately 22% of cells, a higher efficiency of delivery with 22% of cells per 10ug of protein, compared with 3.1% of cells per 10ug of protein in the LR10 condition (**Fig 6d,e**).

**Figure 6:**
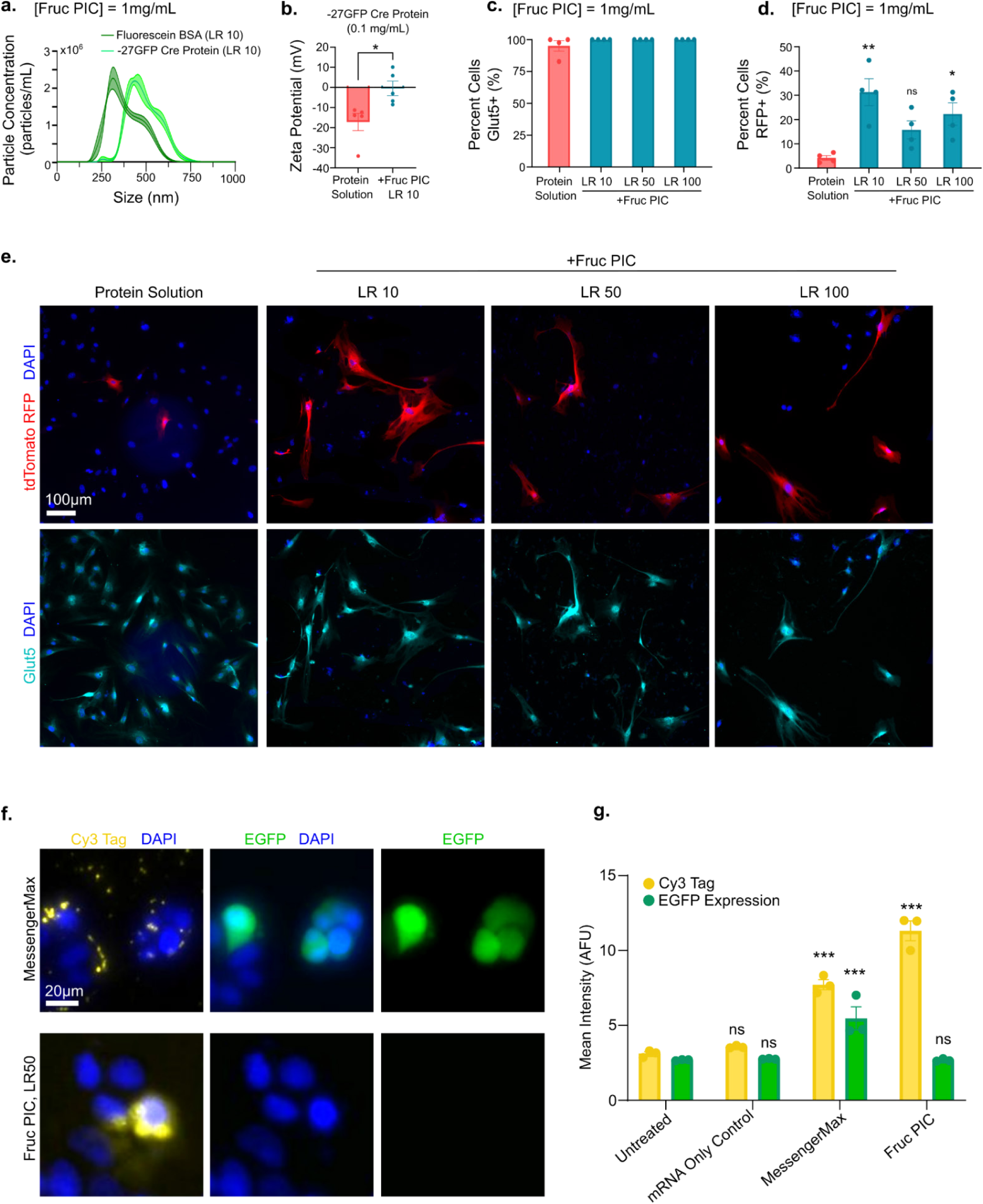
Functional delivery of Cre protein to Ai14 mouse derived fibroblasts in vitro. (a) Nanoparticle tracking analysis (NTA) size distribution comparison between the F-BSA loaded Fructose-PIC used in previous experiment and the Cre protein loaded Fructose-PIC used in this figure. (b) zeta potential of charge neutralization of -27GFP Cre protein upon complexation with dendrimers with a C:D ratio of 1:2 using 25C/F75 fructose dendrimer, dendrimer to protein mass loading ratio of 10, and concentration of 0.1 mg/mL of protein. Analyzed with unpaired t-test. (c) Quantification of immunocytochemical staining of Glut5 on Ai14 (tdTomato Cre reporter) mouse tail derived fibroblasts in vitro, normalized to total cell population, no significant difference across groups. (d) Quantification of immunocytochemical staining of tdTomato indicating functional intracellular delivery of Cre protein, to Ai14 fibroblasts in vitro across Fructose-PIC loading ratios 10, 50, and 100, where LR10 and LR100 treatments had significantly higher tdTomato expression compared to protein solution control, normalized to total cell population. (e) Immunocytochemical staining of tdTomato, Glut5, and DAPI on Ai14 fibroblasts treated with either Cre protein solution in PBS or Cre protein loaded Fructose-PICs at LR 10, 50, 100 for 4 days. The scale bar is 100 micron, n=4, analyzed with one-way ANOVA, with Dunnett’s multiple comparisons test compared against the protein solution control group. (f) Live-imaged MCF-7 cells after treatment with a MessengerMax positive control, and Fructose-PIC formulated at LR50 and 1:2 C:D ratio, concentration at 0.5mg/mL, all complexed with Cy3-tagged EGFP coding mRNA, such that green EGFP signal is indicative of functional mRNA intracellular delivery, and stained with NucBlue live DAPI. (g) Quantification of mean intensity of signal from live imaging the Cy3 mRNA tag and expressed EGFP, analyzed with two way ANOVA and Sidak’s multiple comparisons test, n=3. All quantifications show mean +/-SEM (standard error of mean), *p-value <0.05, **p-value <0.01, ***p-value <0.001.

Following effective functional protein delivery *in vitro*, we next tested whether Fructose-PICs could deliver functional mRNA, as this would expand the potential applications of the carrier to nucleic acid-based therapies. Using a Ribogreen assay, typically employed to characterize mRNA encapsulation efficiency for lipid nanoparticles (LNP), we determine that both the Fructose-PICs and naked-PICs could effectively complex mRNA at a LR of 10 (**Fig S18**). Using Cy3-tagged EGFP mRNA we treated MCF-7 cells *in vitro* with neutral Fructose-PICs at a concentration of 0.5mg/mL and evaluated functional expression of the EGFP protein at 2 days after treatment, using the standard reagent MessengerMax as a control mRNA carrier for comparison. While we detected a high abundance of Cy3-positive mRNA on, and potentially in, cultured MCF-7 cells when delivered using the Fructose-PIC, we ultimately detected no translated EGFP protein using the Fructose-PIC carrier. By contrast, effective translation and expression of EGFP protein was robustly detected using the MessengerMax carrier (**Fig 6f,g**). The source of mRNA delivery failure, whether an issue of inappropriate particle size since PICs are much larger than standard lipid nanoparticles, failure of mRNA to sufficiently de-complex from the PIC carrier, or insufficient mRNA stability with PIC complexation are uncertain. Ultimately, further work to refine PIC formulations so they are specifically tailored for mRNA delivery would likely lead to improved outcomes.

### Fructose-PICs injected into mouse striatum induce a concentration dependent inflammatory response

To assess the *in vivo* foreign body response to Fructose-PICs, we injected formulations with increasing concentrations into the mouse striatum (**Fig 8a**). The striatum is a suitable anatomical site for initial biocompatibility and delivery assessments in the CNS, since it is easily surgically accessible and permits consistent and reproducible injections of materials^60^. To track the dispersion of Fructose-PICs throughout the striatum, we loaded 10kDa biotinylated dextran amine (BDA) into the dendrimer complex, which is a bioinert tracing agent that can be readily detected by immunohistochemical methods applied to fixed tissue sections (**Fig 8a**). PICs were formulated with a constant 1mg/mL BDA solution while increasing dendrimer to BDA cargo LRs serially from 0 (BDA solution only) to 64 to probe Fructose-PIC concentration dependent biocompatibility. A 1uL volume of fructose-PICs loaded with BDA was injected into healthy mouse striatum and mice were perfused for tissue collection at 7 days post injection. The toxicity profile of the Fructose-PICs *in vivo* largely mirrored that observed for the various *in vitro* cell viability assays, with formulations at 1mg/mL inducing minimal neural tissue responses equivalent to that provoked by the BDA solution only control. At 4 and 8 mg/ml, Fructose-PICs elicited increasing neuroinflammation as evident by elevated Gfap-positive astrocyte reactivity as well as increasing numbers of CD45- and Iba1-positive reactive microglia, but notably minimal neuronal toxicity. Fructose-PICs at concentrations of 16 mg/ml or higher caused severe neural tissue toxicity resulting in neuronal loss and the formation of CNS injury-like compartmentalized lesions at the injection site which contained non-neural cores densely filled with peripherally-derived CD13-positive macrophages (**Fig 7b-e S18**). At the injection site of the 8mg/ml Fructose-PIC dose, we detected increased Glut5 expression, mostly restricted to astrocytes that were forming borders around the region of tissue damage, in an equivalent manner to that detected at ischemic stroke lesions (**Fig S20, 2d,g**). While the Glut5 expression in reactive astrocytes was striking, there was notably minimal upregulation of Glut5 in Iba1-positive microglia.

**Figure 7:**
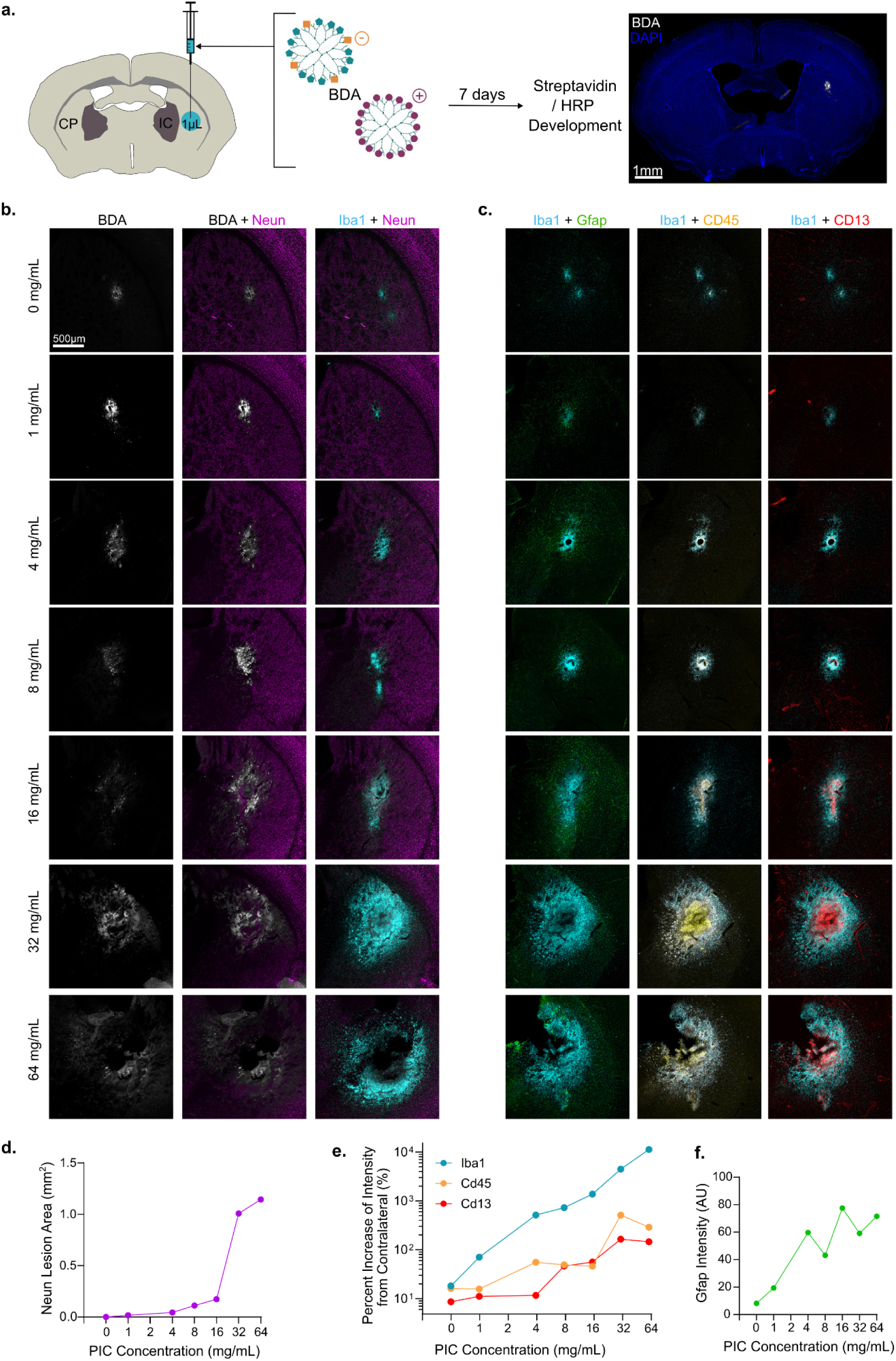
Biocompatibility and tissue response to PIC injection *in vivo.* (a) Experimental schematic showing unilateral striatal injection in the caudate putamen of Fructose-PICs complexed with BDA, and tissue collection and staining for BDA at 7 days post injection. (b) IHC images of striatal injections sites for PICs across concentrations ranging from 0mg/ml (a BDA solution only control) to 64mg/ml, stained for BDA via streptavidin conjugation (white), neuron bodies via NeuN (magenta), and activated microglia and immune cells via Iba1 (cyan). (c) IHC images for the same groups as panel b, stained for activated microglia and immune cells via Iba1 (cyan), reactive astrocytes via Gfap (green), recruited leukocytes via CD45 (yellow), and endothelial cells as well as recruited myeloid cells via CD13 (red). (d) Quantification of area at injection site void of NeuN+ neurons over the increasing concentration of PIC injected. (e) Percent increase in reactive microglial marker, Iba1, and recruited immune cell markers CD13 and CD45 at the injection site, normalized to expression of these markers in the healthy contralateral tissue. (f) Quantification of reactive astrocyte marker Gfap at the injection site over the increasing concentration of PIC injected.

**Figure 8:**
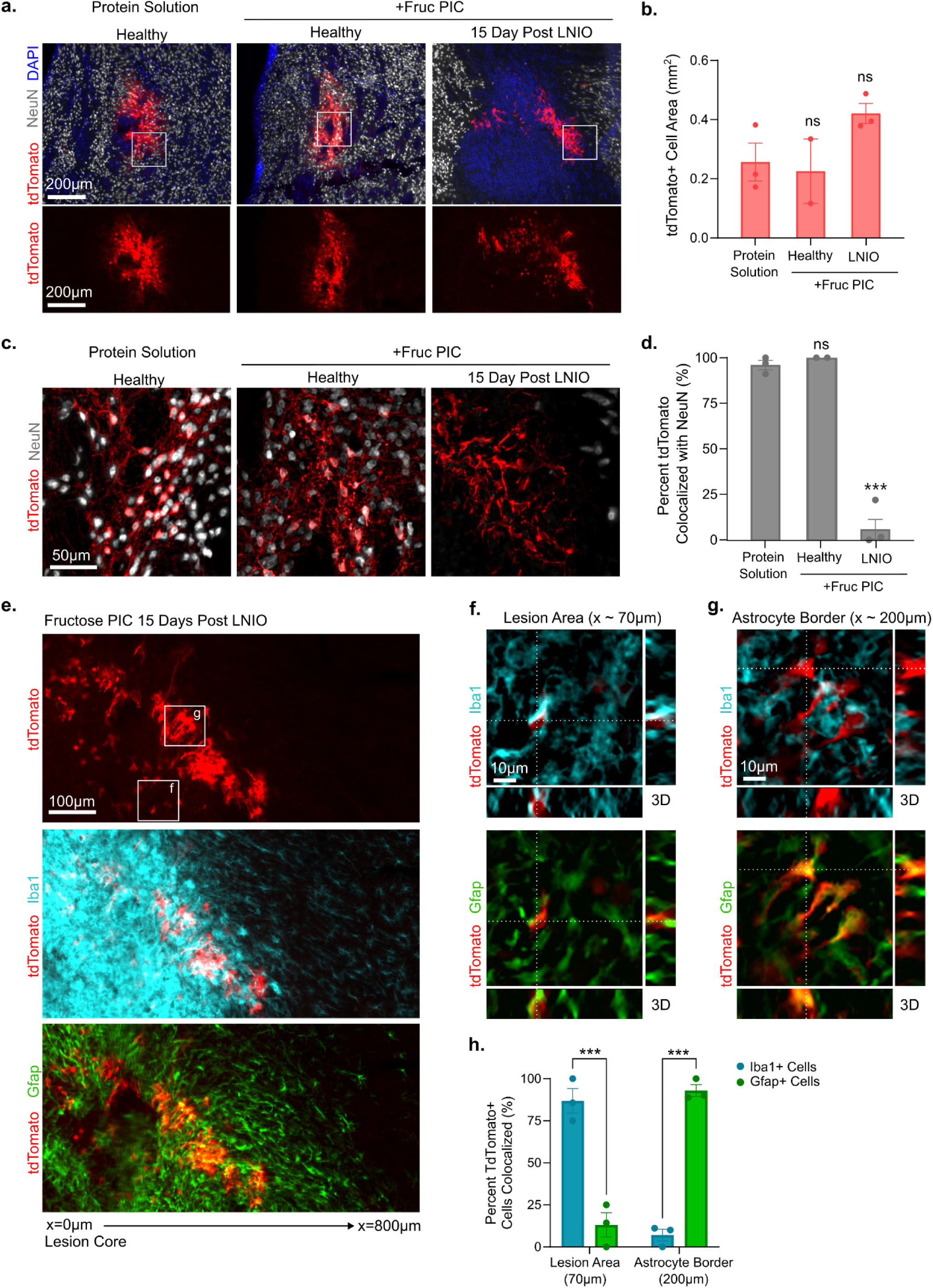
Functional Cre protein delivery to Glut5 expressing reactive glia demonstrated in Ai14 mice. (a) 10x IHC images of cells expressing TdTomato upon functional delivery of Cre in the mouse striatum from injection of either Cre protein in saline solution or Cre protein complexed with a neutral charged Fructose-PIC, in either healthy tissue or 2 days after ischemic stroke, all conditions received 0.1mg/mL of Cre protein, stained for TdTomato (red), NeuN (gray), and Dapi (blue). (b) Quantification of TdTomato positive cell area in all three conditions depicted in panel a, showing no significant difference in the total amount of expression, analyzed with one-way ANOVA. (c) 20x IHC images of TdTomato expression at the injection site co-stained with NeuN (gray) showing neural uptake in the healthy tissue conditions. (d) Quantification of percentage of TdTomato+ cells that are colocalized with NeuN staining, indicating expression in a neuron, analyzed with one-way ANOVA and Dunnett’s multiple comparisons test compared against the protein solution control group. (e) 10x IHC images of TdTomato expression in the border of an ischemic stroke lesion following injection of Cre-complexed Fructose-PICs 2 days post injury, costained with reactive glial markers for microglia (Iba1, cyan) and astrocytes (Gfap, green). (f) 20x z-stack maximum intensity orthogonal projections of Iba1+ microglia, Gfap+ astrocytes, and TdTomato in the lesion region (approximately 70 micrometers radially from the center of the lesion). (g) 20x z-stack maximum intensity orthogonal projections of Iba1+ microglia, Gfap+ astrocytes, and TdTomato in the astrocyte border region (approximately 200 micrometers radially from the center of the lesion). (h) Quantification of the percentage of TdTomato+ cells that colocalize with either Iba1 positive microglia or Gfap positive astrocytes, analyzed with two-way ANOVA and Sidak’s test for multiple comparisons, (n=3). All quantifications show mean +/-SEM, *p-value <0.05, **p-value <0.01, ***p-value <0.001.

For the 1mg/ml concentration Fructose-PIC formulations that induced minimal neural tissue disruption, BDA delivery was detected exclusively in NeuN-positive neurons just like it was for the BDA solution control. Thus it did not appear that the Fructose-PICs meaningfully altered the delivery of the BDA cargo beyond its natural cell uptake path (**Fig 7b, S19**). Based on these results, and since BDA is essentially non-ionic and neutral at a physiological pH, it is likely that there was insufficient loading of BDA into the Fructose-PICs for any fructose mediated delivery mechanisms to be enabled. With increasing Fructose-PIC dose and the associated neural tissue toxicity, BDA was consumed preferentially by microglia and macrophages and was not taken up neurons, suggesting that BDA cargo was being phagocytosed along with the Fructose-PIC material. Ultimately, these data show that at concentrations of 1mg/ml Fructose-PICs are non-toxic and well tolerated but that beyond this concentration Fructose-PICs are noxious to neural tissue such that they provoke increased neuroinflammation and neuronal toxicity at escalating doses.

### Fructose-PICs permit functional protein delivery to healthy and stroke injured mouse brain

To improve our capacity to assess Fructose-PIC mediated delivery *in vivo*, we loaded (–27)GFP-Cre protein into Fructose-PICs and injected these formulated particles into the striatum of Ai14 Cre reporter mice. Unlike BDA, the (–27)GFP-Cre protein is highly negatively charged, so essentially all (–27)GFP-Cre protein is delivered to neural tissue in the form of a complex with Fructose-PICs, thereby improving our capacity to distinguish Fructose-PIC mediated delivery outcomes over those achieved by soluble protein contained within the formulated solution (**Fig S17**). We injected (–27)GFP-Cre loaded Fructose-PICs at 1mg/ml with an LR of 10 into healthy and ischemic stroke injured brains, comparing outcomes with those achieved using the (–27)GFP-Cre protein solution alone. When injected into the healthy striatum, Fructose-PICs delivered functional (–27)GFP-Cre protein exclusively to neurons and did not substantially alter the volume extent or nature of the delivery beyond what was permitted by the protein solution (**Fig 8a-d, S21**). Injecting Fructose-PICs loaded with (–27)GFP-Cre at 2 days after inducing an ischemic stroke by L-NIO shifted protein delivery to microglia and astrocytes locally at the lesion such that we detected minimal numbers of tdTomato-positive neurons in these animals (**Fig 8c-h**). Remarkably, the cell type receiving functional (–27)GFP-Cre delivery was varied by lesion compartment in the ischemic stroke treated mice. Specifically, within the lesion core area, over 80% of detected TdTomato-positive cells were Iba1-positive reactive microglia (**Fig 8f,h**) whereas at the lesion border region essentially all tdTomato-positive cells were Gfap-positive reactive astrocytes (**Fig 8g,h**). The cell type distribution of functional (–27)GFP-Cre delivery mimics the upregulated Glut5 expression pattern noted at 2 day ischemic stroke lesions (**Fig 2e,g,h**), indicating potential fructose-mediated delivery mechanisms that warrant further investigation in the future.

## Discussion

Glial cells are critical players in the pathophysiology of CNS disorders, yet delivering targeted therapies to these cells, especially microglia, remains an unresolved challenge. Achieving effective and cell-type specific drug delivery to glial cells would enable new therapies to treat CNS injury and neurological diseases while minimizing off-target effects that plague current molecular therapies. Here, we leveraged various existing neural cell specific RNA-seq data sets to identify unique surfaceome targets for microglia, astrocytes, and neurons and investigated in detail the potential for targeting one such microglia-specific transporter, Glut5, using nanoparticles with multivalent presentation of the transporter’s primary ligand, fructose. We demonstrated that formulated fructose functionalized particles achieve robust fructose mediated delivery *in vitro* selectively to Glut5-expressing cells but that Glut5 is not an appropriate target for specific delivery to microglia *in vivo*. These findings have broad implications for using transcriptomic information for identifying cell specific targets for drug delivery *in vivo*, for the use of dendrimer-based drug delivery carriers, and for designing microglia specific therapies.

Use of transcriptomic data to identify cellular targets is a growing investigative space with applications ranging from neurodegeneration to autoimmune disorders and oncology. These datasets are often interpreted with the goal of further understanding disease mechanisms, partitioning different cellular states, and identifying cell types and functions that may be altered for therapeutic purposes^86–89^. Significant research in the use of RNA-seq data for cell surface marker identification for drug delivery has come from identifying cellular antigen presentation on cancer cells and studying how these antigens change in different tissues and tumor environments for use in immunotherapy and induction of tumor-specific cell apoptosis^87,88,90,91^. There are two key differences between these studies and our approach: (i) these studies rely heavily on single cell RNA sequencing (scRNA-seq) for their target discovery, which may offer a more specific analysis of the transcriptomics in a focal tissue location but at the cost of losing the depth of reads and larger picture information that bulk sequencing provides, and (ii) in many of these applications, the goal of targeting a particular cell type is to inhibit or modify a signaling cascade by accessing the associated surface-presented receptor or altering extracellular cytokine secretion^89,90,92^, rather than intracellular functional protein delivery.

While our identification of fructose as a specific ligand for targeted delivery to Glut5-expressing cells *in vitro* showed success, the environment for delivery *in vivo* has a multitude of factors that need to be considered when applying an *in silico* identified targeting modality. In this scenario it is important to consider the robustness of expression of the target in the tissue. We found conflicting evidence of our target expression in healthy tissue, with RNA sequencing and in-situ hybridization showing expression by microglia, but staining showing that Glut5 protein localized on astrocytes in the striatal tissue. This, along with the fact that the commercially available antibodies for immunostaining Glut5 were highly variable, hindered our ability to verify our target spatially in the tissue prior to attempted targeted delivery. Additionally, as we demonstrated with the comparison of Cre recombinase delivery via PIC in healthy tissue and ischemic stroke, the injury type, glial reactivity phenotype, and level of Glut5 transporter expression all highly affect the efficacy of delivery. Likely due to the low and diffuse profile of Glut5 expression in healthy striatal tissue, we did not see specificity in the cargo delivery, suggesting that this method may not be an appropriate target and that the use of ligand-transporter interaction based delivery may only be viable in injury pathophysiologies that show cell type specific upregulation of the transporter, such as in acute ischemic stroke lesions. While similar glial reactivity is present in many neurological disorders such as Parkinson’s, Alzheimer’s and Huntington’s disease, further studies would need to assess the expression levels and specificity of potential cell surface transporters in these pathologies, as well as the various glial phenotypes. In future studies, we would likely select targets from the *in silico* RNA-seq analysis in combination with verification of robust protein expression not only in healthy tissue, but also disease and injury states of interest. While Fructose-PICs may not be applicable for targeted drug delivery in the CNS, it may be applicable in the context of cancer cell targeting as breast, lung, renal, and ovarian cancer cells have all been shown to upregulate Glut5^79,93–96^. In fact, high Glut5 expression within tumor cells is often associated with poor prognosis due to fructose intake being linked to high metabolic activity and cellular proliferation^94^, and these highly proliferative cells might be the perfect target to slow or stop tumor growth.

Dendrimers offer a tunable structure that allows for multivalent presentation of ligands for cell surface targeting. In this paper, we implemented a solvent-free series of thiol-ene radically and enzymatically catalyzed reactions to synthesize a biodegradable dendrimer of controllable size via reaction generations, with endcap presentation that is controllable by modulating stoichiometric ratios of endcaps that undergo thiol-ene Michael addition reactions. This synthesis pipeline allowed us to produce dendrimers that were decorated with multivalent presentation of our targeting ligand, fructose, and these dendrimers were able to be complexed with a drug cargo through charge interactions. Dendrimer based drug carriers have been applied to the brain previously, but in these instances have been engineered for systemic delivery by crossing the blood brain barrier, reducing tissue type and cell type targeting specificity^43,93^. Further, they have a common issue of high toxicity due to an excess of cationic charge which is key for endosomal escape upon cellular uptake^97^.

Consistent with this, we discovered that toxicity is a limitation of our drug delivery platform *in vivo*, where the working range of concentrations that do not cause high levels of cellular reactivity and toxicity upon injection in the striatum was relatively small. During *in vitro* testing we found a minimum concentration for efficacy at 0.5 mg/mL, and *in vivo* there was toxicity from the material injection at 8 mg/mL, which gave us a single order of magnitude range in which the Fructose-PICs are usable. This would limit the use of the material to target larger areas of tissue in which a higher concentration may be needed. One way to improve compatibility of the Fructose-PICs would be to modulate the size. We found that formulations of higher concentrations resulted in larger complexes that correlated with higher toxicity. Therefore, using a lower generation dendrimer to reduce molecule size while maintaining a high percentage of fructose presentation could be a way to improve not only biocompatibility but also delivery, since many studies have established that an optimal particle size for intracellular delivery is 50-200nm^98^, a size range that our current Fructose-PICs surpass. A reduction in size via use of a lower generation dendrimer could additionally improve chances of functional mRNA delivery, however the change in size would also change the amount of charged endcap functionality and so cargo loading efficiency would have to be evaluated.

That being said, we demonstrated that by functionalizing our dendrimers with a combination of fructose and ionizable anionic carboxylate endcaps as well as ionizable cationic tertiary amine endcaps, we formulated polyionic complexes that maintained fructose presentation while neutralizing individual dendrimer charges, such that the net charge of the cargo loaded PIC was neutral and cationic charge-based cytotoxicity was reduced. While most nanoparticles in the field depend on cationic charge for intracellular delivery, net neutral nanoparticles have been studied for low toxicity drug delivery, but they are limited to extracellular delivery, such as into the blood stream^99^, or to lipid-based particles that are administered systemically and delivered in a non-specific manner^100^. The efficacy of functional protein delivery via a neutrally charged PIC is an exciting innovation as it allows us to leverage the ligand-transporter interaction to facilitate delivery, resulting in a protein delivery system that is transporter specific and functional without introducing cationic charge-based toxicity. As such, this technology has the potential to be applied to numerous cell targets for different cell types, since the dendrimer model is modular and the fructose endcaps that we synthesized and functionalized onto the dendrimer branches could be replaced with a different ligand for facilitating interactions with a different cell surface target.

While we showed Glut5 specificity of functional protein delivery *in vitro*, this result did not translate *in vivo*, which poses the question of how to best design microglia specific delivery vehicles in the future. Selection of microglia specific targets can be done using RNA-seq analysis, however transcriptomic datasets should be collected from not only healthy neural tissue but also from the disease state of interest. Additionally, transcriptomic analysis would need to be corroborated at the protein expression level as identifiable, quantifiable, and specifically colocalized to microglia in the diseased or injured tissue prior to incorporation on a delivery vehicle. This could be achieved with the use of natural ligands, as we attempted to do in this study, or by using receptor specific nanobodies or antibodies functionalized onto the delivery vehicle. Dendrimers synthesized with the thiol-ene Michael addition click chemistry functionalization would allow for tunable reaction with nanobody proteins, but this targeting technology could be broadened to use with other non-viral drug delivery vehicles such as lipid nanoparticles (LNPs)^101^, coacervates^102^, or engineered virus like particles (VLPs)^103^. In future studies, in order to encourage cellular uptake by microglia, we will consider using nanobodies targeting canonical markers of ramified cell states, such as P2RY12 or CX3CL1, as well as surface receptors known to be upregulated in activated and inflammatory cells states, such as Iba1, CD45, or CD68, which may be key to the use of this targeting method in the context of disease.

## Methods

### In silico RNA-Seq Analysis

A literature search was conducted for studies utilizing ribosome tagged (ribo-tagged) immunoprecipitation (IP) together with bulk RNA sequencing on brain tissue samples that have both input data and enriched data for astrocytes, microglia, oligodendrocytes, and neurons. Datasets were found from studies that used ALDH-Cre ERT mice (GSE84540)^47^, P2Y12-Cre ERT mice (GSE138333)^46^, Olig1-Cre mice (GSE118451)^104^, and Camk2-Cre ERT mice(GSE142678)^45,105^ for ribo-tagged enrichment of astrocytes, microglia, oligodendrocytes, and neurons, respectively. Immunopanned data was sourced from a single study (GSE73721)^106^. The fastq raw data files were then downloaded from Gene Expression Omnibus (GEO) database, trimmed, aligned with Galaxy, and the resulting count files were used to calculate a fragments per kilobase of exon per million mapped (FPKM) value and Z-score for each gene in each sample group. All genes analyzed for Z-scores were then filtered using “the *in silico* human surfaceome” database, created by the Wollscheid lab at ETH Zurich, to keep only surfaceome genes, genes that code for proteins expressed on the exterior cell surface, in our list.

The delta expression, input values subtracted from IP values, was then used in a principal component analysis (PCA) with two components. The same analysis was repeated with an immunopanned dataset^48^. Resulting factor scores and factor loading values allowed for the calculation of the cosine similarity between any given gene and a designated cell type. The eigenvector for a given gene was then calculated in the PCA space, and the angle between the gene vector and the cell type vector was labeled theta. Taking the cosine of theta allowed us to quantify the alignment between the gene and the cell type. The more aligned the gene is with a cell type, the closer the cosine value is to 1. If a gene is orthogonal to a cell type the cosine value is closer to zero. For this reason, the threshold that we assigned to consider a gene to be specific to a cell type is a cosine similarity value of 0.9 or higher. Then, by using a 0.9 threshold for the cosine similarity values and finding genes that were expressed in both the ribo-tagged and immunopanned datasets, the field of available surfaceome genes was reduced to between 130 and 190 per enriched cell type. When genes were sorted from highest to lowest FPKM value, the top 25 genes were selected for further analysis. For each of these genes, a ligand for mode of targeting the encoded protein was identified.

### In Situ RNA Hybridization

Multiplexed FISH was performed as described previously in Moya et al^107^. Briefly, 200 µm coronal sections of striatum were collected from 8-week-old C57BL/6J mice transcardially perfused with 4% PFA in PBS. Sections were pre-treated with 90% DMSO in PBS, 1% NaBH_4_ in PBS, and 8% SDS in PBS to reduce background autofluorescence and increase permeability. They were then treated according to the Molecular Instruments HCRv3 protocol for staining a “generic sample in solution”^108^ and imaged on a Nikon CSU-W1 SoRa spinning disk confocal microscope. All sections were stained for pan-neuronal markers Snap25 and Atp1b1 (B1 – Alexa 488), pan-astrocytic marker Aldoc (B4 – Alexa 594), and fructose transporter Slc2a5 (B5 – Alexa 647). Half the sections were stained for pan-microglial marker Selplg (B3 – Alexa 546), and the other half were stained for pan-microglial marker Tmem119 (B3 – Alexa 546).

### Dendrimer Synthesis

The first reaction in the dendrimer synthesis was completed in a 20mL scintillation vial (Fisherbrand FS74504-20); 0.8mL (or 1.92mMoles) of pentaerythritol tetrakis (3-mercaptopropionate) (PETMP, Sigma 381462-100ML) was combined with 1.6mL (or 8.06mMoles) of trimethylolpropane ally ether (TMPAE, Sigma 416118-500ML) and stirred thoroughly until mixture appears homogeneous. Then 30mg (or 0.12mMoles) of 2,2-Dimethoxy-2-phenylacteophenone (DMPA, Sigma 196118-5G) is added as a reaction photoinitiator. The solution is then stirred continuously while exposed to 365nm UV light (AloneFire SV005 10W LED, X002M6TKY3) in a blackout reaction chamber for 1 hour. At the end of 1 hour, thin layer chromatography (TLC, MilliporeSigma HX45371841) confirmed that the first reaction had run to completion. For the second reaction, 4mL (or 31.43mMoles) of ethyl 3-mercaptopropionate (E3MP, TCI Chemicals M1038-25G) was added to the reaction vial, followed by 200mg of *Candida antarctica* lipase B (CALB, Novozym 435) as a reaction catalyst and 1g of 4Å molecular sieves (4AMS, ThermoScientific L05454.0B). The reaction solution was then mixed and placed on a shaker (IKA KS 4000i Control) at 50°C with 250rpm orbital shaking for 96 hours. At the conclusion of the second reaction, TLC was again used to confirm the reaction had run to completion. The CALB and 4AMS were then filtered out of the product by running product through cotton in a glass pipette under air pressure into a new 20mL scintillation vial. To facilitate recovery of product from the reaction vial and allow for easier filtration, ethyl acetate (Fisherbrand E145-4) was added to the product before transferring to the filter. The product was then rotovapped (IKA RV8, IKA Vacstar control, IKA HB digital) to remove solvent. For synthesis of higher generation dendrimers, the entire process was repeated, replacing the PETMP starting material with the purified first generation dendrimer, and scaling molar ratios accordingly. Once the desired generation was complete, crude material was purified by repeatedly washing in 50mL of hexanes (Fisherbrand H292-4) until TLC showed that all unreacted species were removed. Material was then rotovapped until dry to remove any remaining solvent, and the final product was stored in a 20mL scintillation vial, sealed with tape, at 4°C until characterization or functionalization.

### Dendrimer Scaffold Characterization

Unfunctionalized dendrimer scaffolds were characterized by nuclear magnetic resonance (NMR) and Fourier transform infrared spectroscopy (FTIR). For NMR characterization, a portion of the final dendrimer product was put into solution at approximately 10mg/mL in deuterated dimethyl sulfoxide (dDMSO, ThermoScientific 166290500) and transferred to an NMR tube (FisherScientific 664000585). NMR spectroscopy was a 32 scan proton test using a 500MHz frequency (Agilent 500 MHz VNMRS). Analysis of NMR spectra was completed using MestReNova version 14.1.0-24037.

For FTIR characterization, the material was applied to the quartz window of the Nicolet 4700 FTIR-ATR, undiluted and in a viscous liquid form. Analysis of FTIR spectra was completed using MATLAB version R2021a and GraphPad Prism version 10.6.

### Fructose Acrylate Synthesis

The fructose acrylate reaction was completed in 100mL of 2-methyl-2-butanol (2M2B, Alfa Aesar A18304) solvent in a 200mL Erlenmeyer flask. Firstly, 1g of fructose anhydrous (fructose, Sigma F0127-100G) was added to the 2M2B and heated to 50°C on a shaker (IKA KS 4000i Control) with 250rpm orbital shaking for 30 minutes. Following the 30-minute period, the flask was removed from the shaker, and 1g of 4AMS, 20mg of 4-methyloxyphenol (MEHQ, TCI Chemicals M0123), and 250mg of CALB were added to the reaction flask. This was heated for 10 minutes on the orbital shaker at 50°C. After 10 minutes, the flask was removed from the shaker and 780uL of 2,2,2-trifluoroethyl acrylate (TFEA, TCI Chemicals A18235.14) was added to the reaction. The flask was then returned to the shaker at 50°C and reaction was allowed to proceed for 72 hours, covered with aluminum foil. TLC was used to monitor reaction progression at 24-hour intervals. At the 72-hour time point, the reaction flask was removed from the shaker to stop the reaction. Reaction contents were vacuum filtered with 11um cellulose filtration paper (Whatman 1001-055), and rotovapped down to approximately 10mL volume. This material was dried with Celite and purified with flash chromatography using a two-solvent gradient of ethyl acetate (Fisherbrand E145-4) and methanol (Fisherbrand A412-4). Resulting fractions were run on TLC to identify the location of the purified product, and the product-containing fractions were pooled into a 500mL round-bottom flask and rotovapped down to remove solvent. The final product was stored in a 20mL scintillation vial, sealed with tape, at 4°C until characterization or functionalization.

### Fructose Acrylate Characterization

Fructose monoacrylate was characterized with nuclear magnetic resonance (NMR) and Fourier transform infrared spectroscopy (FTIR) as described above.

### Dendrimer Functionalization Reaction

Dendrimer functionalization was completed with the assumption that the second reaction of dendrimer synthesis proceeded to completion. Thus, for a single one-generation dendrimer molecule, there are 8 available thiol groups for functionalization reactions. The functionalization reaction was completed in a 10mL round-bottom flask, in which 100mg of purified dendrimer product was solubilized in 1mL of N,N-dimethylformamide (DMF, Sigma D158550-1L). After solubilization, a molar ratio of endcap (fructose monoacrylate, 2-Carboxyethyl acrylate (“C”, Sigma 55234), 2-(Dimethylamino)ethyl acrylate (“D”, Sigma 330957), 3-Sulfopropyl acrylate potassium salt (“S”, Sigma 251631), 2-(Acryloyloxy)ethyl trimethylammonium chloride solution (“A”, Sigma 496146)) was added to the reaction flask, followed by N,N-diisopropylethylamine (DIEA, TCI Chemicals D1599) at a ratio of 1mole DIEA to 3mole of reactable thiols on the dendrimer. This reaction was stirred at room temperature for 1 hour; TLC at the 1-hour time point confirmed that the reaction had proceeded. After reaction completion, the product was rotovapped to remove excess DMF and DIEA from the solution. Once solvents were removed, water was added to the product, and if functionalized with ionizable endcaps, then either hydrochloric acid (for protonation of D) or sodium hydroxide (for deprotonation of C) was added to the suspended material to put it into a charged state; the volume of acid or base correction was based on the molar amount of charged groups added to the dendrimer. Product solubilized in water was then transferred to a 50mL conical tube containing ethyl acetate, and the tube was shaken to wash the product with ethyl acetate and remove any excess reactants. This was then centrifuged for 10 minutes at 4.4krpm to form a liquid-liquid separation, and the ethyl acetate supernatant was removed. The solution was then transferred to a 20mL scintillation vial and rotovapped to ensure removal of all ethyl acetate and resolubilized in water. The water-soluble material was then lyophilized for 3 days to achieve a dry material stock, which was stored sealed with tape at 4°C until characterization or use for in vitro and in vivo studies.

### Fructose Functionalized Dendrimer Material Characterization

Bulk fructose-functionalized dendrimer material was characterized with nuclear magnetic resonance (NMR) and Fourier transform infrared spectroscopy (FTIR), as described in section 5.4 above, and matrix-assisted laser desorption/ionization-time of flight (MALDI-TOF).. Dry functionalized dendrimer was solubilized in either DI water or PBS, for cationic and anionic functionalizations respectively, at highly concentrated stock solutions of 200 mg/mL. For characterization of zeta potential and dynamic light scattering (DLS) (Brookhaven NanoBrook Particle Size and Zeta Potential Analyzer), material was diluted to 10mg/mL in PBS and filtered with a 0.22um PES syringe -driven filter. For MALDI-TOF characterization, the material was solubilized in methanol, mixed with DBH matrix (Sigma-Aldrich 50862-10MG-F) solubilized in 50/50 methanol and water, and spotted onto a polished steel 384-spot plate to dry at room temperature. Material was scanned with a reflective protocol (RP) test, set to a range of 900-4500 Da, on the Bruker Autoflex TOF instrument. Analysis of MALDI-TOF spectra was completed using FlexControl version 3.4.

### TNS Assay

A buffer was prepared consisting of 10mM HEPES (Fisherbrand BP310-100), 10mM MES (Fisherbrand BP300-100), 10mM sodium acetate (Fisherbrand BP333-500), and 140mM sodium chloride (Sigma 71376-5KG). This solution was then separated into different tubes and pH adjusted with hydrochloric acid (Honeywell 258148-500ML) and sodium hydroxide (Sigma 72068-100ML) to create a set of buffers ranging in pH from 2 to 11 at 0.5 intervals. Functionalized dendrimers were then solubilized or suspended in each buffer at a concentration of approximately 0.5mg/mL. TNS solution was prepared by dissolving 1mg of TNS (Abcam ab275049) into 160mL of deionized water and mixing thoroughly, resulting in a final concentration of 20uM. For each dendrimer at each pH 100uL of sample solution was added to a polystyrene, non-treated 96-well plate in triplicate wells. After all samples were added to the plate, 10uL of TNS solution was added to each well for a working concentration of 2uM TNS per well. The plate was covered with aluminum foil to prevent photobleaching of fluorescence and incubated for 10 minutes. The fluorescence was then quantified on the Molecular Devices SpectraMax i3X Microplate Detection Platform (plate reader) at an excitation of 321nm and emission of 445nm. Raw fluorescence values were plotted in GraphPad Prism, averaging over the technical replicates. Each dataset was fitted with a three-parameter sigmoidal curve, and the EC50 value determined for that fit was taken as the pKa value of that dendrimer. For each functionalized dendrimer, this assay was repeated at least three times to calculate experimental replicate pKa values for each material, and these were averaged for the final comparison of pKa between different dendrimer functionalities.

### PIC Formation and Cargo Loading

G5-D dendrimers were solubilized in deionized water to make high-concentration stock solutions at 200mg/ml and 50mg/ml. G5-C/F materials were solubilized in pH 7.4 PBS to make high-concentration stock solutions at 200mg/ml and 50mg/ml. All stock solutions were sterile-filtered with a 0.22 micron PES membrane. Delivery cargos (fluorescein BSA (MilliporeSigma A9771-1G), FITC-IgG (Sigma-Aldrich F9636-1ML), Cytochrome C (Sigma-Aldrich C2506-250MG), BDA (Invitrogen D1956), or Cre-protein (GenScript neg27GFP-GGS9-Cre-6xHis)) were solubilized at 10mg/mL in pH 7.4 PBS and filtered. Ionic complexes were formed using two different complexation orders: cargo + anionic G5-C/F + cationic G5-D, or anionic G5-C/F + cationic G5-D + cargo, to test the effect of order on complexation and delivery efficiency. All complexations were done at 10mg/mL of total dendrimer material in PBS and then diluted to the desired concentration.

### MCF-7 Culture

MCF-7 cells were kindly provided by the Grinstaff lab at Boston University. MCF-7 cells were cultured from frozen stock in a T75 flask with growth medium: Eagle’s Minimum Essential Medium (MEM, Corning 10-009-CV) with 10% fetal bovine serum (FBS, Gibco A52567-01) and 0.01mg/mL of human insulin (MP Biomedical 193900). Cells were maintained in a 37C incubator with 5% CO2, and media was changed every two days while expanding cells. Cells were passaged at 80-90% confluency, typically every 4-5 days. To passage, growth medium was aspirated from the flask, cells were washed once with sterile PBS, and trypsinized with 0.05% Trypsin EDTA (cat no) for 3-5 minutes in the incubator. After cells had lifted off the flask, trypsin was neutralized with equal volume of unsupplemented MEM, cells were transferred to a sterile 50mL conical tube and centrifuged at 100 RCF for 8 minutes. Supernatant was aspirated without disturbing the cell pellet, and the pellet was resuspended in 5mL of growth medium. To count the cells, 10uL of thoroughly mixed cell suspension was combined with 10uL of trypan blue (ThermoFisher 15250-061), and 10ul of this was transferred to a hemocytometer chip to be counted either manually on a brightfield microscope or with the Countess automatic cell counter (Invitrogen AMQAF2000). All experiments were performed with cells at a passage number between 22 and 29.

### mAstrocyte Culture

Mouse neural progenitor cells (NPCs) were cultured from frozen stock in a T75 flask in neural expansion (NE) media consisting of Advanced DMEM/F12 (ThermoFisher 12634-010), 1x B27 (ThermoFisher 12587010), 1% Non-essential amino acids (ThermoFisher 11140050), 1% Nucleosides (EMD Millipore ES-008-D), 1% Antibiotic/Antimyotic (ThermoFisher 15240062), 100ug/mL of heparin, 100ng/mL fibroblast growth factor (FGF, Peprotech 100-18B-100UG), and 100ng/mL of epidermal growth factor (EGF, Peprotech AF-100-15-100UG). Cells were maintained in a 37C incubator with 5% CO2, and media was changed every two days while expanding cells. Cells were passaged at 80-90% confluency, typically every 4-5 days. To passage, NE was aspirated from the flask, cells were washed once with sterile PBS, and trypsinized with 0.05% Trypsin EDTA (cat no) for 3 minutes in the incubator. After cells had lifted off the flask, trypsin was neutralized with equal volume of unsupplemented DMEM/F12, cells were transferred to a sterile 50mL conical tube and centrifuged at 1000 rpm for 5 minutes. Supernatant was aspirated without disturbing the cell pellet, and the pellet was resuspended in 5mL of NE medium. To count the cells, 10uL of thoroughly mixed cell suspension was combined with 10uL of trypan blue, and 10uL of this was transferred to a hemocytometer chip to be counted either manually on a brightfield microscope or with the Countess automatic cell counter. Once NPCs were plated and at desired confluency, NE media was removed and replaced with astrocyte differentiation media consisting of 1% FBS and 100ng/mL CNTF (Peprotech 450-13-100UG). Cells were incubated in astrocyte differentiation media for 48 hours prior to use in assays. All experiments were performed with cells at a passage number between 14 and 25.

### Cytotoxicity Assay

Cells were seeded in a 96-well polystyrene tissue culture-treated plate at a density of 50,000 cells per well in 100uL of growth medium. Cells were incubated for 24 hours after seeding to allow for adherence to the plate. At 24 hours, cells were treated with PICs at desired concentrations, formulated as described in section 5.10, including an untreated condition as a live cell control. All groups were run in triplicate. Cells were incubated in treatment conditions for 24 hours at 37C and 5% CO2. After 24 hours, cells for a negative dead cell control were treated with 80% methanol in water for 30 minutes. Following this, treatment media was aspirated, wells were washed with 200uL of sterile 1x PBS, and 100uL of 2μM Calcein AM (Corning 354217) solution in PBS was added to each well. Cells stayed in Calcein solution for 30 minutes at room temperature, then the plate was read on a plate reader with fluorescence settings of 488nm excitation and 520nm emission. Cell viability percentages were calculated by normalizing treatment group raw fluorescence values to the fluorescence values of live cell and dead cell controls.

### MCF-7 In Vitro Competition Assay

Cells were cultured in growth medium as described in section 5.11, and seeded in either a 6-well polystyrene tissue culture-treated plate or on 0.1% gelatin-coated (Sigma G1890-100G) 12mm round coverslips (Fisherbrand 22293232) in a 24-well polystyrene plate. Cells on coverslips were seeded at a lower density of 44,000 cells/cm^2^ to allow for clear visualization of individual cells with cytochemical staining, whereas cells in the 6-well plate were seeded at a higher density of 90,000 cells/cm^2^ close to confluency to be used for higher-throughput quantitative fluorescence reading via Countess. Cells were incubated for 24 hours after seeding in a growth medium to allow for adherence to the plate. At 24 hours post-seeding, growth medium was removed, cells were washed once with sterile PBS, and treatments were applied. All treatments, including untreated control groups, were done in a serum-free media consisting of DMEM/F12 (ThermoFisher 11320-033) with 1x B27 and 100ng/mL FGF supplemented. High-fructose media conditions consisted of solubilizing fructose at 100mM in serum-free media and sterile filtering with a 0.22 micron PES membrane before use. PICs were formed and loaded with fluorescein BSA cargo as described in section 5.10 in low volumes of sterile PBS and then diluted out with media. After treatment conditions were applied, all groups were incubated for 24 hours, after which cells on coverslips were fixed for cytochemical staining or analyzed with the Countess.

### Astrocyte In Vitro Competition Assay

NPCs were cultured in NE medium as described in section 5.12 and seeded in a 6-well polystyrene tissue culture-treated plate at a higher density of 90,000 cells/cm^2^, close to confluency, to be used for higher-throughput quantitative fluorescence reading via Countess. Cells were incubated for 24 hours after seeding in NE medium to allow for adherence to the plate. At 24 hours post-seeding, NE medium was removed, cells were washed once with sterile PBS, and astrocyte differentiation media was applied for 48 hours as described in section 5.12. After 48 hours, the differentiation medium was removed, cells were washed once with sterile PBS, and treatments were applied. All treatments, including untreated control groups, were done in a serum-free media consisting of NE media with CNTF at 100ng/mL supplemented. High fructose media conditions consisted of solubilizing fructose at 100mM in serum-free media and sterile filtering with a 0.22 micron PES membrane before use. PICs were formed and loaded with fluorescein BSA cargo as described in section 5.10 in low volumes of sterile PBS and then diluted out with media. After treatment conditions were applied at 1mg/mL of total dendrimer material and at a dendrimer:protein mass loading ratio (LR) of 10, all groups were incubated for 24 hours, after which cells were analyzed with the Countess.

### Cytochemical Staining

Coverslips seeded with cells were washed with PBS and fixed in 4% paraformaldehyde (Electron Microscopy Sciences 15714-S) for 30 minutes. Coverslips were then washed with PBS 3 times and stored in PBS with 0.01% sodium azide in a parafilm-sealed plate at 4°C until use. Coverslips were stained by first washing six times with TBS, then incubating in a phalloidin stain solution at 10uL phalloidin / 1mL of TBS solution for 25 minutes. At 25 minutes, DAPI stain was added to the phalloidin solution for a total DAPI concentration of 4uL DAPI / 1mL, and incubated for an additional 15 minutes. Coverslips were then washed 6 times with TBS and mounted on slides with ProLong Gold mounting media and stored away from light as the slides dried overnight.

### Countess Hight Throughput Fluorescence Quantification

At the end of treatment incubation, treatment media is removed from the 6 well plate, and cells are washed with 2 mL of sterile PBS. Cells are then trypsinized with 1.5mL of 0.05% trypsin for 3 minutes. Trypsin was neutralized with unsupplemented MEM or DMEM/F12 for MCF-7 cells or astrocytes respectively, transferred to 15mL conical tubes, and centrifuged according to cell type protocols. Media was aspirated without disturbing the cell pellet and cells were resuspended in 0.5mL of sterile PBS. For each condition, 10uL of cell suspension was transferred to a Countess hemocytometer chip and the chip was inserted for automatic counting. Light gating was held constant for both brightfield (value of 66) and green fluorescence (value of 90) across all samples. Three separate preparations and measurements were performed for each condition. Measurement settings collected the position, radius, size, circularity, brightfield brightness, fluorescence brightness (RFU), and area of each individual cell in the field, as well as an overall cell count. These data were compiled and exported in a .JSON file, which was then extracted and analyzed using MATLAB. RFU value thresholds to consider a cell to be positive for fluorescein BSA were established in the manual processing of the data in MATLAB, not in the Countess software.

### In Vitro Transcriptomics

In vitro transcriptomic datasets were sourced from the NCBI GEO database. Mouse neural progenitor cell derived astrocyte data were acquired from accession number GSE194319. MCF-7 data were acquired from accession number GSE240542.

### Western Blotting

Cell lysate was collected from a confluent T75 flask using RIPA buffer (ThermoScientific XL359231) for each cell type and media condition. Total protein concentration was measured using a Pierce BCA Total Protein kit (ThermoScientific 23225) prior to loading a gel for blotting. Electrophoresis was run on Tris-Bis 4-12% 10 well 1mm mini protein gels (Invitrogen NP0321BOX) using an Xcell SureLock (Invitrogen EI0001) system and NuPAGE Running Buffer (Invitrogen NP0001). For each sample, 30ug of total protein of cell lysate was added to a 0.6mL tube, combined with sample buffer (Invitrogen NP0007) and DI water for a total volume of 20uL. These samples were heated at 70C for 10 minutes and briefly centrifuged to collect all liquid at the bottom. 20uL of sample was added to each well, except for protein ladder wells, which received 10uL of Novex Sharp prestained protein standard (Invitrogen LC5800). After all wells were loaded, electrophoresis was run at a constant 150mV for approximately 1.5 hours. Following electrophoresis, gels were transferred onto PVDF membranes (ThermoScientific XJ3570311) which had been activated in methanol, rinsed in DI water, and saturated in NuPAGE transfer buffer (Invitrogen NP0006-1), run at a constant 30mV for 1 hour. Following transfer, total protein was stained using SimplyBlue SafeStain (Invitrogen 465034) following provided product protocol. The membrane was then de-stained, washed for 10 minutes three times with TBS + 0.1% Tween20 (TBST) and blocked in a solution of 5% milk in TBST for 1 hour at room temperature. Following blocking, the membrane was incubated in 3% milk in TBST solution with rabbit Glut5 ProteinTech primary antibody (ProteinTech 27571-1-AP) at 1:1,000 dilution, overnight at 4C. On the second day, the membrane was washed three times with TBST and incubated in 3% milk in TBST solution with anti-rabbit HRP-conjugated secondary antibody for 1 hour and washed 3 times with TBST; all incubations were done with gentle shaking. The membrane was then laid flat and left stationary while incubated with SuperSignal WestFemto (ThermoScientific 34094) for 5 minutes. Finally, the membrane was placed between two transparencies and imaged with optimal exposure chemiluminescence (BioRad ChemiDoc MP Imaging System). Intensity of total protein stain as well as chemiluminescence was quantified using ImageJ.

### Ai14 Fibroblast Harvesting and Culture

B6.Cg-*Gt(ROSA)26Sor^tm^*^14^(CAG–tdTomato)*^Hze^*/J (Ai14) mice (Jax #007914) were perfused with PBS containing heparin, and tails were removed and cleaned with 70% isopropyl alcohol, then submerged in cold PBS. Using sterile forceps and surgical scissors in a sterile biosafety cabinet, skin was removed from the mouse tail and the tail tissue was cut into 1cm long pieces, which were added to a gelatin-coated well of a 6-well polystyrene plate containing 2mL of fibroblast growth medium (DMEM/F12 (ThermoFisher 11320-033) with 10% FBS (Gibco A52567-01) and 1% antibacterial/antimicrobial (ThermoFisher 15240062)). Tissue was removed from the plate after 5 days of incubation at 37C with 5% CO2. Cells were passaged upon reaching confluency and seeded on gelatin-coated 12mm coverslips for ICC as described in section 5.14 and treated with Cre protein loaded PICs at 1mg/mL in serum-free medium for 4 days before fixation. Coverslips seeded with cells were washed with PBS and fixed in 4% paraformaldehyde (Electron Microscopy Sciences 15714-S) for 30 minutes. Coverslips were then washed with PBS 3 times and stored in PBS with 0.01% sodium azide in a parafilm-sealed plate at 4°C until use. Coverslips were stained by first washing six times with TBS, then incubating in a blocking and permeabilization solution using 1× TBS containing 5 % Normal donkey serum (NDS, Jackson ImmunoResearch Laboratories) and 0.5 % Triton x-100 for 1 h. Coverslips were then stained with primary antibodies overnight in 1× TBS containing 0.5 % Triton x-100. The following primary antibodies were used in this study: anti-RFP (guinea pig, 1:500) and anti-Glut5 (rabbit, 1:200). Coverslips were then washed 6 times with TBS and incubated in secondary antibodies in 1× TBS containing 0.5 % Triton x-100 and 5 % NDS for 2 hours at room temperature. All secondary antibodies were affinity purified whole IgG (H + L) purchased from Jackson ImmunoResearch Laboratories, with donkey host and target specified by the primary antibody. Cell nuclei were counter stained with 4′,6′-diamidino-2-phenylindole dihydrochloride (DAPI; 2 ng/ml; Molecular Probes) prior to mounting onto glass slides with ProLong Gold (Invitrogen, Cat# P36934) anti-fade reagent. Stained cell coverslips were imaged using epifluorescence and deconvolution epifluorescence microscopy on an Olympus IX83 microscope.

### In Vitro mRNA Delivery Assay

MCF-7 cells were cultured as described in section 5.11 and seeded in a 96 well tissue culture-treated polystyrene plate at 100,000 cells per well in 100uL of growth medium, and incubated for 24 hours to allow for cell adhesion. Dendrimers were complexed with Cy3 tagged GFP mRNA (APE Bio R1008, 996 nucleotides, arca cap, 5-moUTP modified) in RNAse and DNAse free PBS. Lipofectamine 3000 (Invitrogen L3000001) was used as a positive control and was loaded with Cy3 tagged GFP mRNA as per product instructions. Cells were washed once with sterile PBS, then mRNA loaded PICs and Lipofectamine were diluted in serum-free MCF-7 media and added to the cells and allowed to incubate for 24 hours. After 24 hours, NucBlue live nuclear stain (Invitrogen R37605) was added to each well as per product instructions, and the plate was live imaged using epifluorescence microscopy on an Olympus IX83 microscope to determine the amount of Cy3 tagged mRNA present, the amount of GFP expression, and the number of cells in each well. Images were analyzed using ImageJ.

### Surgical Methods

All surgical procedures were approved by the BU IACUC (Protocol number: PROTO20200045) and conducted within a designated surgical facility. All procedures were performed on either C57/BL6 female mice (Cat#000664, JAX) or Ai14 (Cat #007914, JAX) female mice, all were aged 8–12 weeks at the time of craniotomy. Mice were positioned into a rodent stereotaxic apparatus using ear bars (David Kopf, Tujunga, CA) and maintained under general anesthesia achieved through inhalation of isoflurane (1–3%) in oxygen-enriched air. Using the aid of a stereo microscope, a small rectangular shaped craniotomy was performed over the left coronal suture using a high-speed surgical drill. A small flap of bone encompassing sections of the frontal and parietal bone was removed to expose the surface of the brain prior to injection. BDA solution (1mg/mL), Cre-protein solution (0.1mg/mL), BDA loaded neutral Fructose-PICs (1mg/mL - 64 mg/mL), or Cre loaded neutral Fructose-PICs (1mg/mL) in a solution of sterile PBS were loaded into pulled borosilicate glass micropipettes (WPI #1B100-4) that were ground to a 35° beveled tip with 150–250 μm inner diameter after formulation with sterile-filtered components. Glass micropipettes were mounted to the stereotaxic frame and high-pressure polyetheretherketone tubing and connectors were used to attach micropipettes to a 10 μL syringe (Hamilton, Reno, NV, #801 RN) that was mounted into a syringe pump (Pump 11 Elite, Harvard Apparatus). Loaded BDA, Cre protein or PICs were injected into the caudate putamen nucleus of the striatum at 0.15 μL/min using target coordinates relative to Bregma: +1.0 mm A/P, +2.5 mm L/M, and −3.0 mm D/V. A total volume of 1 μL was used for all conditions. Once the complete volume had been injected, the micropipette was allowed to dwell in the brain at the injection site for an additional 4 min before being slowly removed from the brain incrementally over a 2-min period. After injection, the surgical incision site was sutured closed with polypropylene sutures and the animals were allowed to recover.

### Tissue Fixation and Immunohistochemistry

At 7 or 13 days after striatal injections, mice underwent terminal anesthesia by overdose of isoflurane and were then perfused transcardially with heparinized saline and 4 % paraformaldehyde (PFA) using a peristaltic pump at a rate of 7 ml/min. Approximately 10 ml of heparinized saline and 50 ml of 4 % PFA were used per animal. Brains were dissected and post-fixed in 4 % PFA for 6–8 h followed by cryoprotection in 30 % sucrose with 0.01 % sodium azide in Tris-Buffered Saline (TBS) for at least 3 days at 4 °C prior to cryosectioning. Coronal brain sections (40 μm thick) were cut using a cryostat (Leica CM1950 Cryostat). Tissue sections were stored in 96well plates immersed with 1× TBS and 0.01 % sodium azide solution at 4 °C. For immunohistochemistry (IHC), tissue sections were stained using a free-floating staining protocol outlined in detail previously [24,55]. Following 1 N HCl antigen retrieval and serial washes, the tissue sections were blocked and permeabilized using 1× TBS containing 5 % Normal donkey serum (NDS, Jackson ImmunoResearch Laboratories) and 0.5 % Triton x-100 for 1 h. Tissue sections were then stained with primary antibodies overnight in 1× TBS containing 0.5 % Triton x-100. The following primary antibodies were used in this study: anti-Rabbit Glut5 (ProteinTech 27571-1-AP, 1:200), anti-Guinea pig Neun (Synaptic Systems, 266 004, 1:500), anti-Rat Gfap (Invitrogen, 13–0300, 1:1000); anti-Guinea pig Iba1 (Synaptic Systems, 234,004, 1:500); anti-Goat CD13 (R&D systems, AF2335, 1:500); anti-Rat CD45 (R&D Systems AF14, 1:500), anti-Goat Sox9 (R&D Systems AF3075, 1:500), anti-Rat PU1 (Biolegend 681302, 1:500), anti-RFP (Abcam Ab62341, 1:500). Tissue sections were washed three times and incubated with appropriate secondary antibodies diluted at 1:250 in 1× TBS containing 0.5 % Triton x-100 and 5 % NDS for 2 h. All secondary antibodies were affinity purified whole IgG (H + L) purchased from Jackson ImmunoResearch Laboratories, with donkey host and target specified by the primary antibody. BDA was visualized using streptavidin-HRP plus Tyr-Cy5 (PerkinElmer). Cell nuclei were counter stained with 4′,6′-diamidino-2-phenylindole dihydrochloride (DAPI; 2 ng/ml; Molecular Probes) prior to mounting onto glass slides and then cover slipped with ProLong Gold (Invitrogen, Cat# P36934) anti-fade reagent. Stained sections were imaged using epifluorescence and deconvolution epifluorescence microscopy on an Olympus IX83 microscope utilizing the 10x, 20x, and 40x objectives and the following fluorescent channels: 488nm (FITC), 555nm (TRITC), 647nm (Cy5), 790nm (Cy7), 405nm (DAPI).

### Quantitative analysis of immunohistochemistry

Immunohistochemical staining intensity quantification for the various antibody markers or BDA was performed on tiled images prepared on the Olympus IX83 epifluorescence microscope that were taken at a standardized exposure time across groups and using the raw/uncorrected intensity settings. Quantification of total antibody staining or BDA accumulated in tissue was performed using NIH ImageJ version 1.53k, and orthogonal projections were done with Zeiss ZEN software.

## Statistical analysis

Statistical evaluations were conducted by one-way or 2way ANOVA with post hoc independent pairwise analysis via Tukey’s or Sidak’s multiple comparisons test where appropriate using Prism 10 (GraphPad Software Inc, San Diego, CA). Across all statistical tests, significance was defined as p-value <0.05. All graphs show mean values plus or minus standard error of the means (S.E.M.) as well as individual values overlaid as dot plots.

## Acknowledgements

We would like to thank the Grinstaff Lab at Boston University for generously providing an initial cryostock of MCF-7 cells, and thank undergraduate researchers Han Ren Leong and Mincheol Kim for their help with initial material synthesis troubleshooting. This research is supported by the Biomedical Engineering Core Facilities at Boston University. A special thanks to Xin Brown and the Biointerface Technologies (BIT) core, as well as the Micro and Nano Imaging (MNI) core at BU. This research was also supported by the BU Chemistry Department Chemical Instrumentation Center (CIC) and staff.

## Funding

We are grateful to the National Science Foundation for the purchase of the NMR (CHE0619339) and MALDI-TOF mass spectrometer (CHE1337811) used in this work. This research was supported by funding from Boston University Start-Up Funds (T.M.O), Boston University’s Undergraduate Research Opportunities Program (UROP) (G.A), Boston University BUNano PhD Fellowship (P.S.W), National Institute of Neurological Disorders and Stroke (NIH NINDS) R21NS133525 (T.M.O).

## Notes

### Competing Interest Statement

The authors have declared no competing interest.

