## Supplementary Information for "Fructose Dendrimer-Based Poly Ionic Complexes for Glut5-Specific Intracellular Drug Delivery in Murine Brain"

* Corresponding author

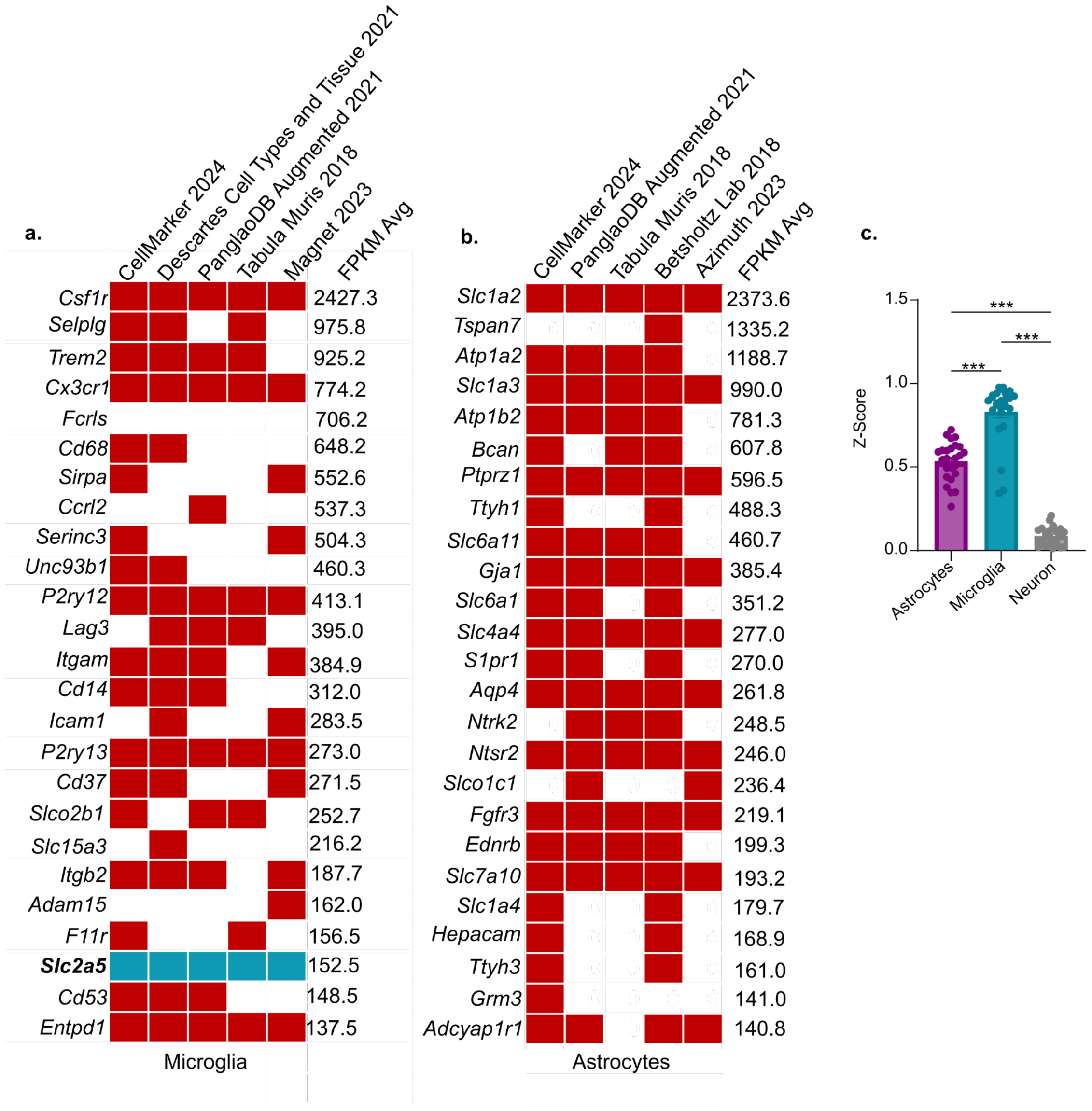


**Supplemental Figure 1: Single cell sequencing corroboration of cell specific surfaceome genes.** (a) Binary heatmap of whether an identified microglia specific surfaceome gene is established as microglia specific via single cell sequencing across five different single cell sequencing databases, slc2a5 highlighted in teal. (b) Binary heatmap of whether an identified astrocyte specific surfaceome gene is established as astrocyte specific via single cell sequencing across five different single cell sequencing databases, for both panels (a and b) a red box indicates a positive result, that the gene is present in the database as cell type specific, a white box indicates a negative result. (c) Comparison of the z scores of the top 25 cell specific surfaceome genes for astrocytes, microglia, and neurons, with the microglia z scores being significantly higher indicating a greater separation from the general population distribution. Panel C shows mean +/- SEM (standard error of mean) and was analyzed with one way ANOVA and Tukey's test, ***p-value <0.001.

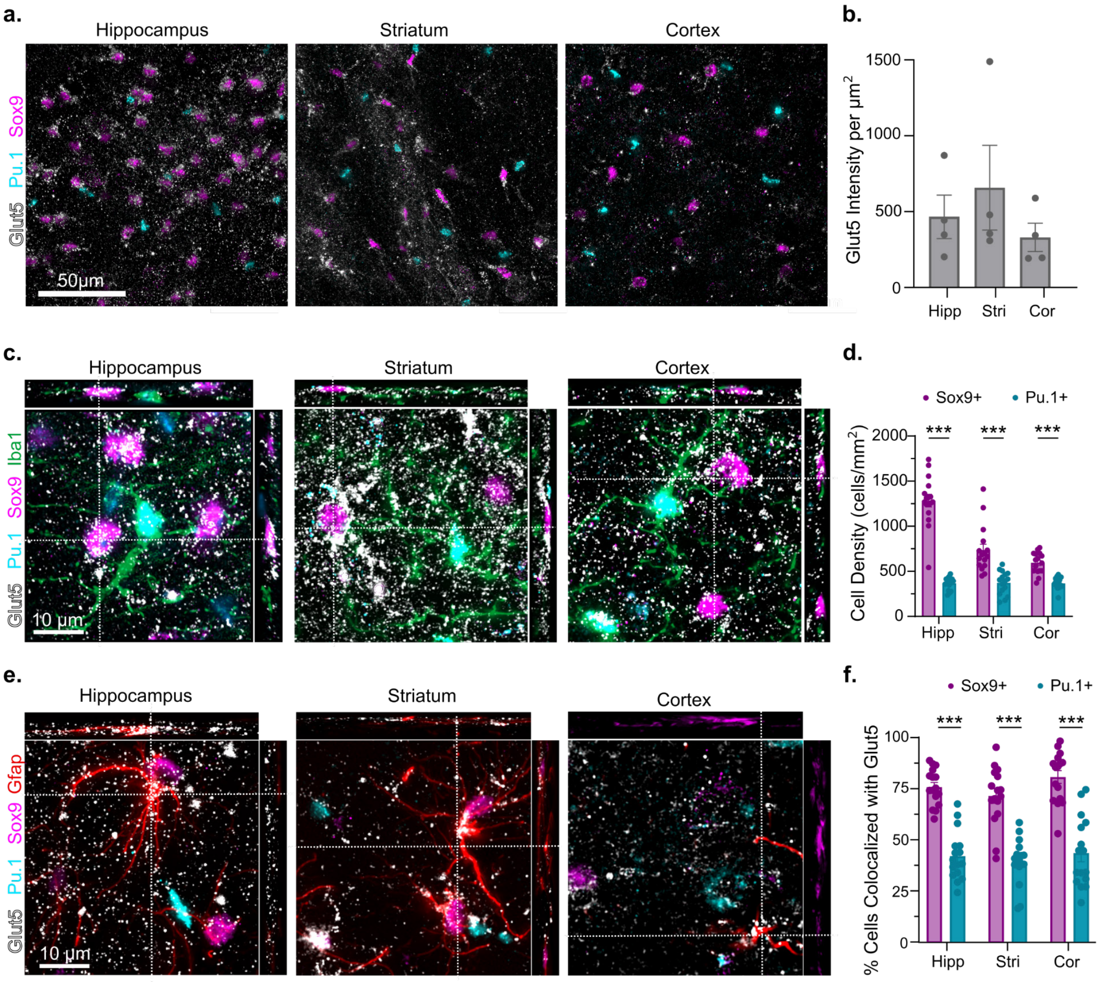


**Supplemental Figure 2: IHC staining of Glut5 across brain regions.** (a) IHC images of Glut5 staining (white) in the hippocampus, striatum, and cortex brain regions, costained with Pu.1 (cyan) as a nuclear stain for microglia, and Sox9 (magenta) as a nuclear stain for astrocytes, scale bar is 50µm. (b) Quantification of Glut5 staining intensity normalized to area for all three tissue regions depicted. (c) IHC images with orthogonal projections showing lack of alignment of Glut5 staining (white) on Pu.1+ (cyan) and Iba1+ (green) microglia compared to Sox9+ (magenta) astrocytes. (d) Quantification of Pu.1+ and Sox9+ cell density in each tissue region, showing a higher density of astrocytes. (e) IHC images with orthogonal projections showing Glut5 staining (white) on Gfap+ (red) and Sox9+ (magenta) astrocytes. (f) Quantification of the percentage of Pu.1+ microglia or Sox9+ astrocytes with Glut5 staining, demonstrating that Glut5 staining occurs on astrocytes. Quantifications, panels b, d, and f, show mean +/- SEM (standard error of mean) and were analyzed with one way and two way ANOVA, ***p-value <0.001.

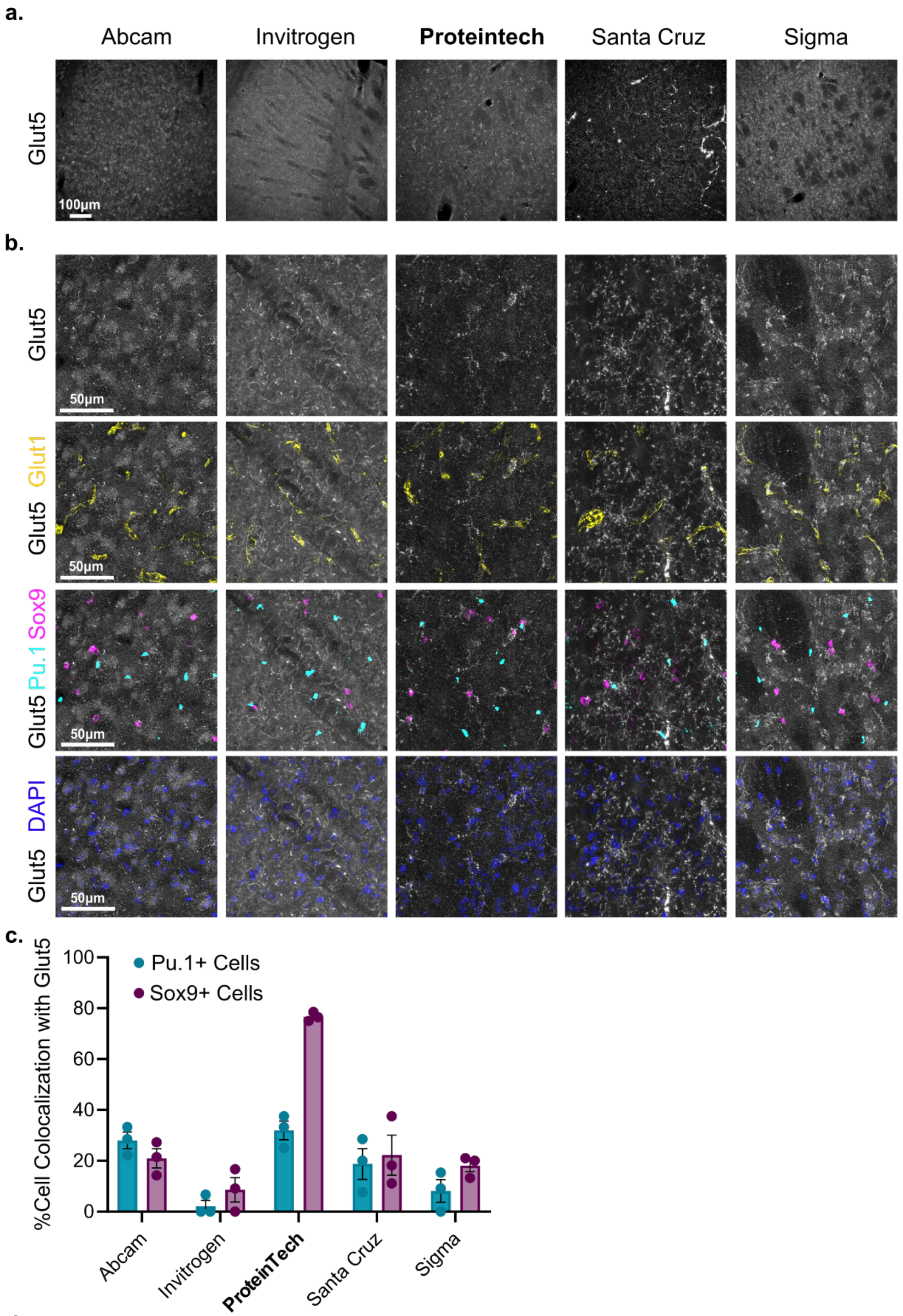


d.

| Vendor: | Abcam | Invitrogen | **Proteintech** | Santa Cruz Biologicals | Sigma |
|---|---|---|---|---|---|
| Catalog #: | ab41533 | PA5-80024 | 27571-1-AP | sc-271055 | AV42096 |
| Origin | Polyclonal | Polyclonal | Polyclonal | Monoclonal | Polyclonal |
| Species: | Rabbit | Rabbit | Rabbit | Mouse | Rabbit |
| Dilution used: | 1:200 | 1:200 | 1:200 | 1:100 | 1:200 |

**Supplemental Figure 3: Comparison of available Glut5 antibodies in IHC.** (a) IHC images of healthy mouse striatum of each Glut5 antibody in 1mm square areas. (b) IHC images of healthy mouse striatum of each Glut5 antibody, costained with Glut1 glucose transporter, Pu.1+ microglia nuclei, Sox9+ astrocyte nuclei, and DAPI nuclear labeling. (c) Quantification of the amount of the percentage of Pu.1+ and Sox9+ cells that colocalize with the staining of each Glut5 antibody. (d) Product details of each antibody tested.

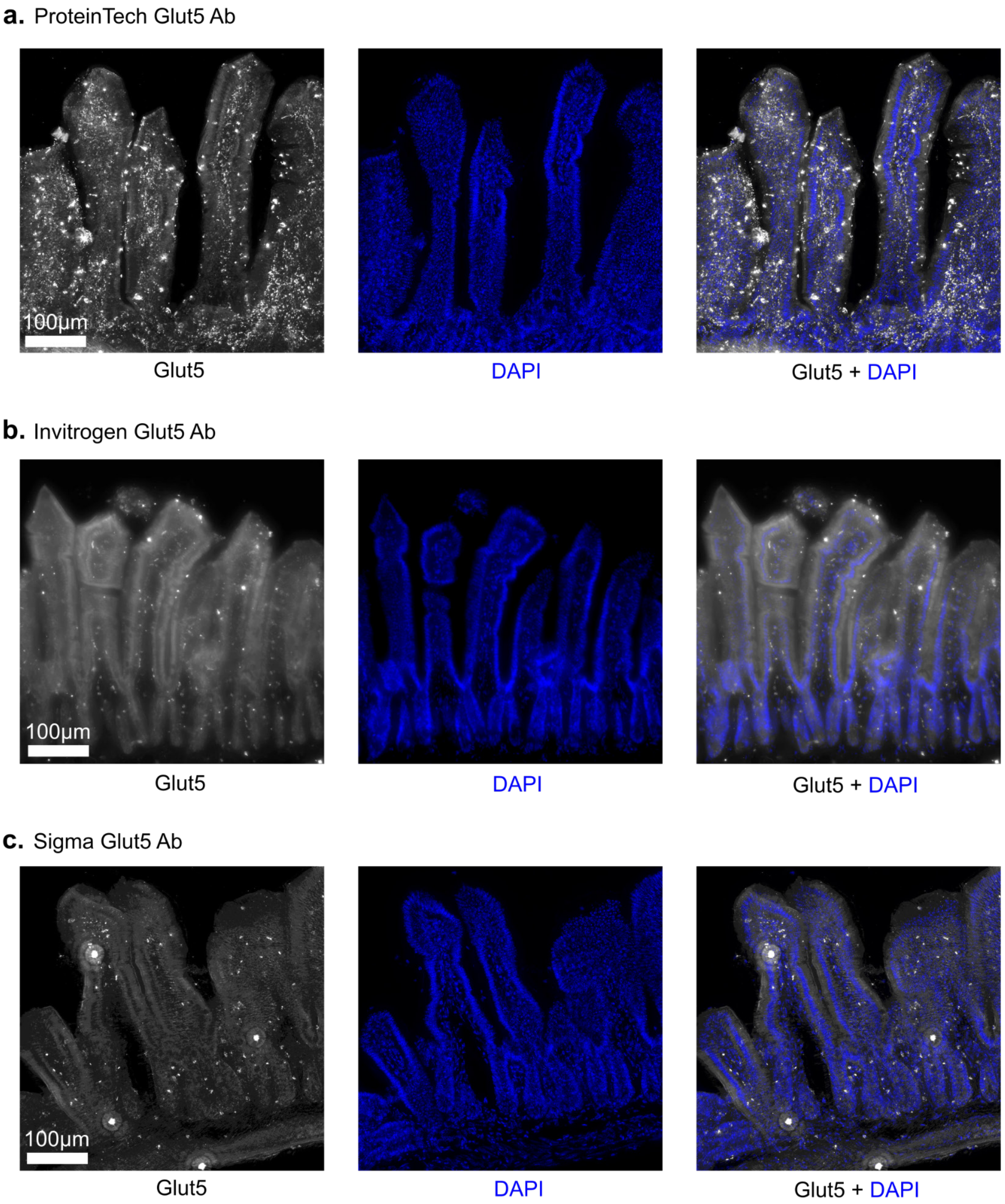


**Supplemental Figure 4: Glut5 antibodies tested in mouse small intestine.** IHC images of mouse jejunum stained with the (a) ProteinTech Glut5 primary antibody, (b) Invitrogen Glut5 primary antibody, (c) Sigma Glut5 primary antibody, all using anti-rabbit Cy5 labeled secondary antibodies and costained with DAPI nuclear labeling.

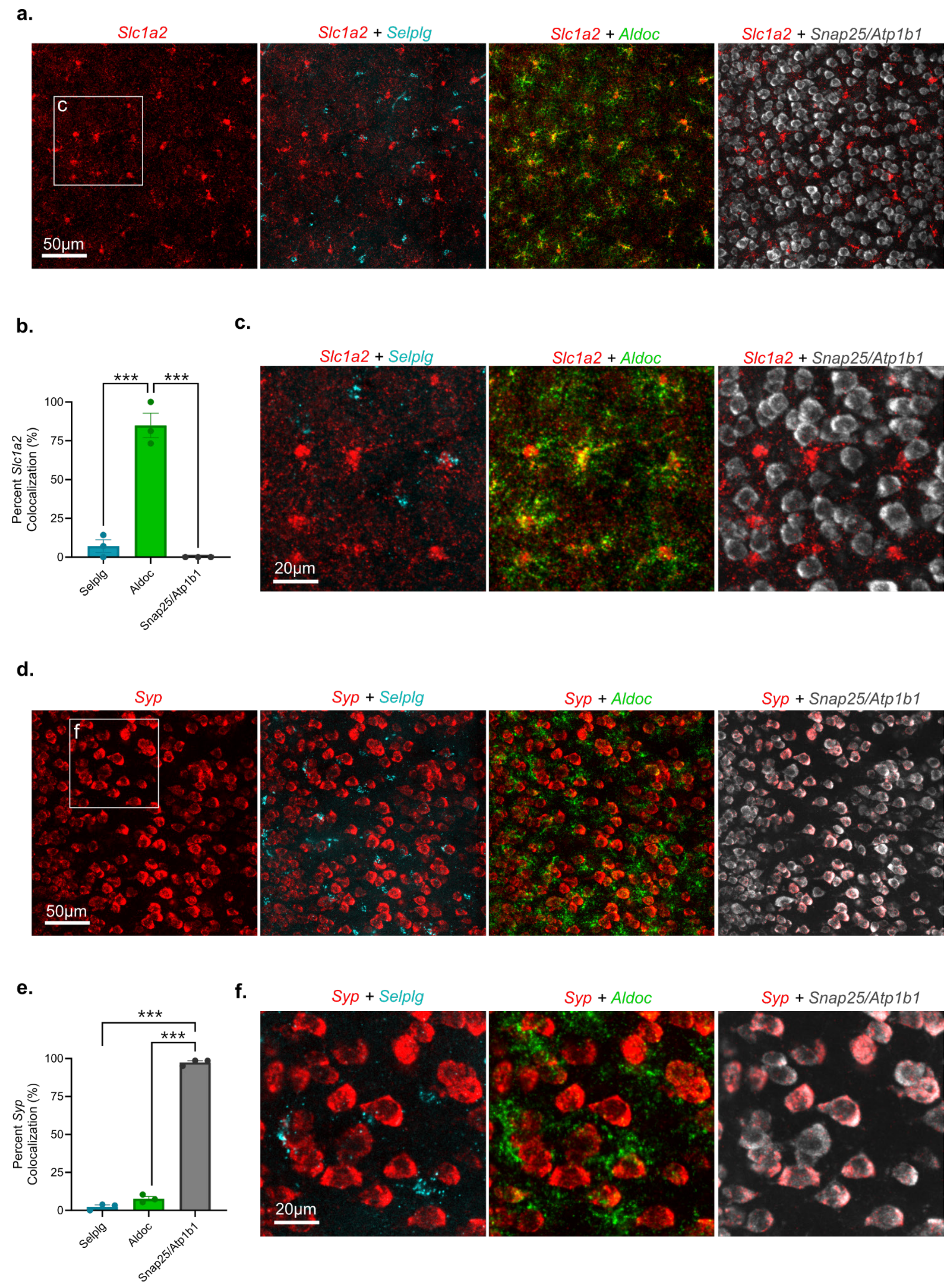


**Supplemental Figure 5: In Situ RNA of astrocyte and neuron transporters.** (a) In situ RNA probe images in healthy striatum of astrocyte specific glutamate transporter gene Slc1a2 (red), co-probed with Selplg to show microglia (cyan), Aldoc to show astrocytes (green), and a combination of Snap25 and Atp1b1 as a pan neuron marker (gray). (b) Quantification of colocalization of in situ hybridization Slc1a2 spots with each of the cell markers, showing high and specific expression in Aldoc positive astrocytes. (c) Closer cropped images from panel a. (d) In situ RNA probe images in healthy striatum of neuron specific synaptic vesicle protein gene Syp (red), co-probed with Selplg to show microglia (cyan), Aldoc to show astrocytes (green), and a combination of Snap25 and Atp1b1 as a pan neuron marker (gray). (e) Quantification of colocalization of in situ hybridization Syp spots with each of the cell markers, showing high and specific expression in Snap25 and Atp1b1 positive neurons. (f) Closer cropped images from panel d. Quantifications show mean +/- SEM, analysis done with one way ANOVA and Tukey's test, ***p-value <0.001.

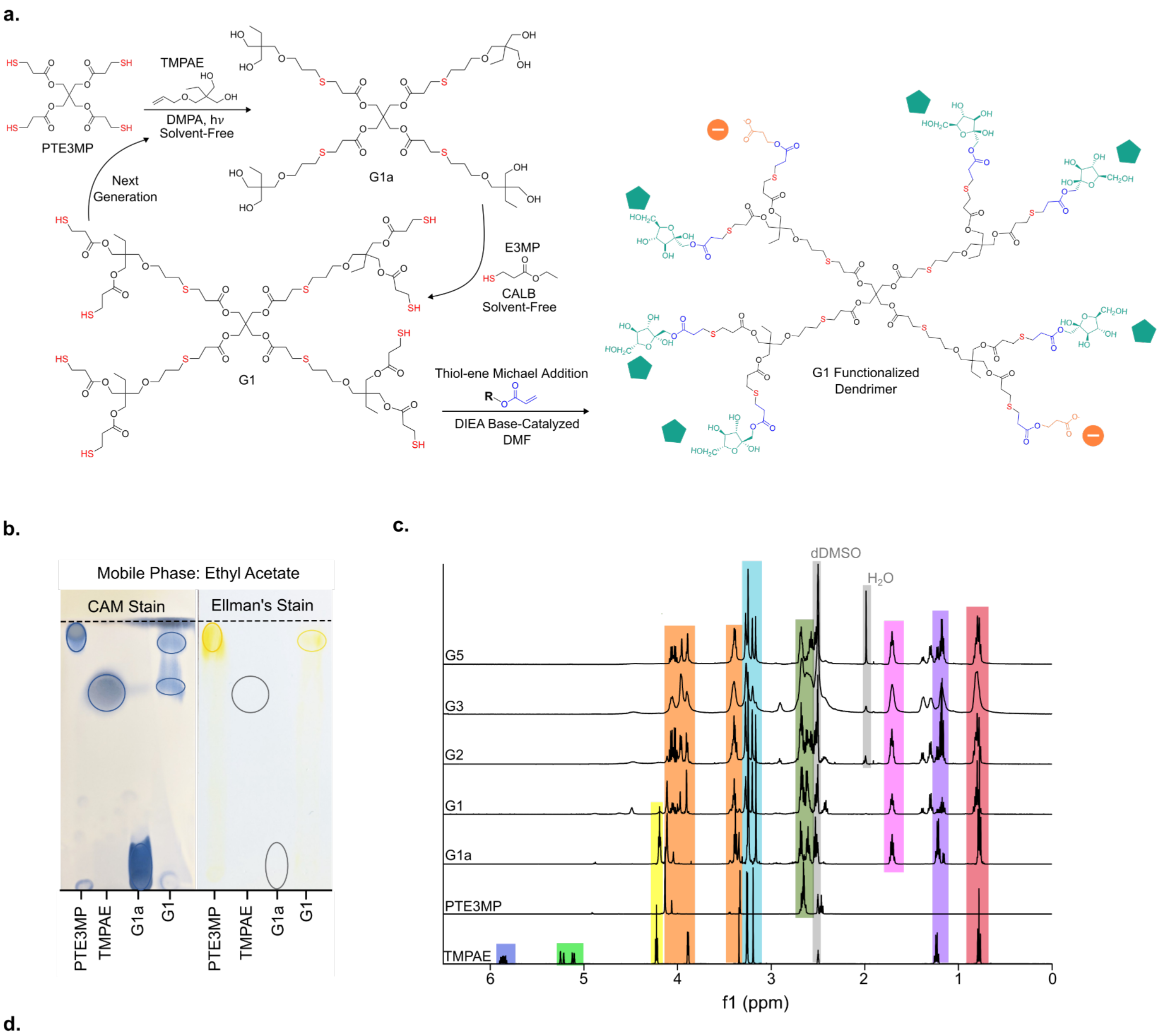


d.

| Shift (ppm): | 5.9-5.8 | 5.3-5.1 | 4.2 | 4.1-4.0 | 3.4 | 3.3-3.1 | 2.8-2.6 | 1.7 | 1.2 | 0.8 |
|---|---|---|---|---|---|---|---|---|---|---|
| G5 | 0.00 | 0.00 | 0.15 | 2.82 | 2.40 | 4.76 | 4.22 | 1.71 | 5.99 | 3 |
| G3 | 0.00 | 0.00 | 0.11 | 3.72 | 4.32 | 3.76 | 6.40 | 2.02 | 4.84 | 3 |
| G1 | 0.00 | 0.02 | 1.75 | 3.73 | 4.16 | 4.66 | 8.01 | 1.63 | 4.82 | 3 |
| G1a | 0.00 | 0.01 | 2.03 | 2.36 | 5.87 | 6.34 | 4.83 | 1.84 | 2.32 | 3 |
| TMPAE | 0.87 | 1.92 | 1.92 | 2.00 | 0.83 | 6.18 | 0.02 | 0.00 | 2.01 | 3 |

**Supplemental Figure 6: Dendrimer scaffold synthesis.** (a) Full scaffold structure reaction scheme, (b) TLC of reactants and both first and second step reaction products stained with CAM to track all material and Ellman's to track the loss and gain of thiols as the reaction progresses, (c) NMR of each reactant and product, including higher generation dendrimers formed with the repetition of the reaction cycle, (d) Integrations of NMR shifts, all normalized to the three protons on the $CH_3$ at 0.8ppm that occur on a single branch of the dendrimer.

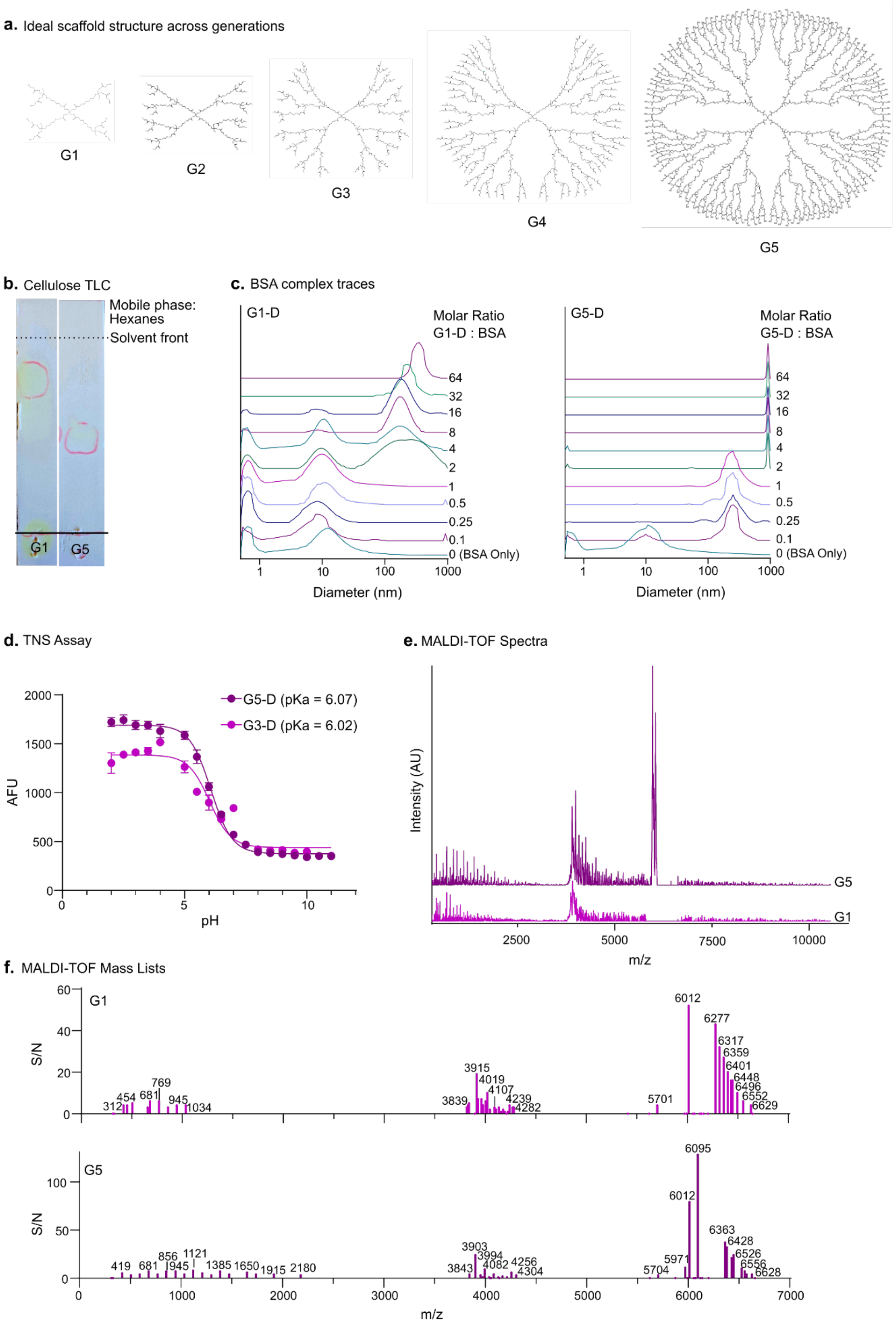


**Supplemental Figure 7: Dendrimer scaffold generational growth.** (a) Structure drawings of generations 1-5 assuming ideally complete reactions,(b) Comparison of run distance of a G1 and G5 on a cellulose TLC plate stained with Ellman's reagent, (c) DLS traces of BSA complexed G1-D and G5-D dendrimers demonstrating increased multivalency in higher generations, (d) TNS fluorescence assay demonstrating an increased total fluorescence of higher generations indicating presentation of more positively charged endcaps, while conserving pKa, (e) MALDI-TOF spectra of G1-D and G5-D material showing increased relative abundance of high molecular weight species in the G5-D, (f) MALDI-TOF mass lists of G1-D and G5-D material showing increased high molecular weight fragments and an increased distribution of fragment species in the G5-D material.

**a.** Reaction scheme

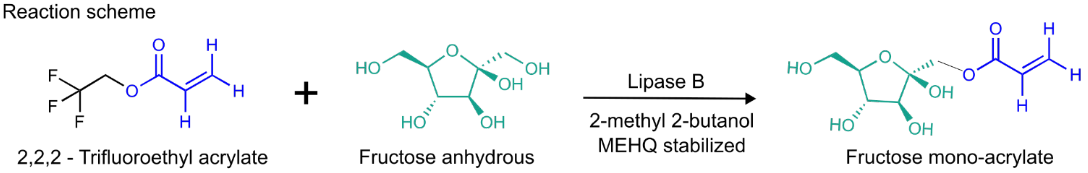


**b.** TLCs: CAM stained, mobile phase is 17:4:1 ethyl acetate:methanol:water, plates are silica on glass

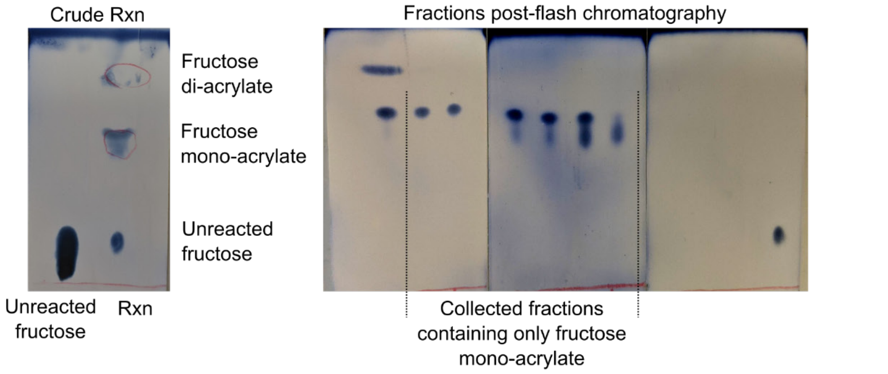


**c.** NMR

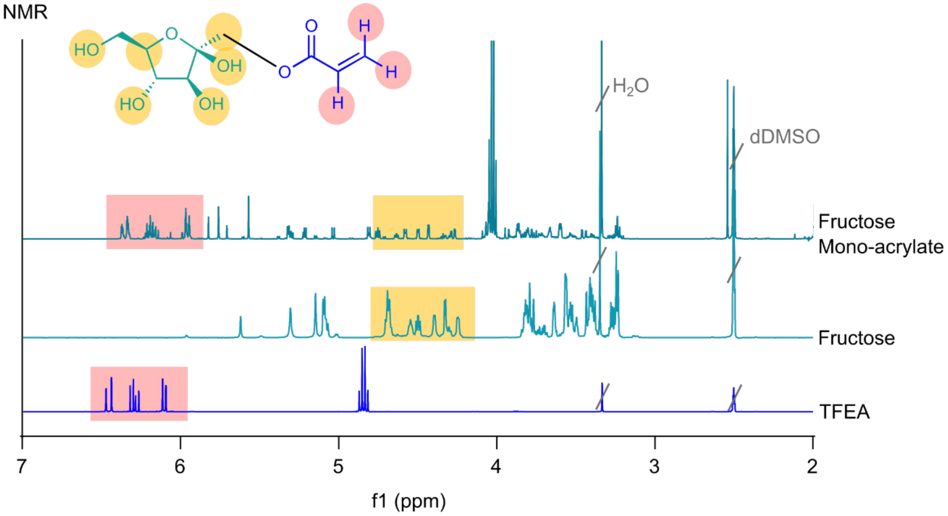


**d.** NMR Shift Integrations

| Shifts: | 6.5 - 6 | 6-5 | 5-4.8 | 4.8-4.2 | 4.2-4 | 4-3.3 | 3.3-3.1 |
|---|---|---|---|---|---|---|---|
| Fructose Mono-Acrylate | 1 | 0.78 | 0.02 | 0.95 | 1.63 | 1.53 | 0.24 |
| Fructose | 0.0 | 0.54 | 0.01 | 1 | 0.0 | 1.74 | 0.44 |
| TFEA | 1 | 0.0 | 0.70 | 0.01 | 0.0 | 0.01 | 0.0 |

**Supplemental Figure 8: Fructose acrylate synthesis.** (a) reaction scheme, (b) TLCs pre and post flash chromatography purification showing separation of di-acrylate, mono-acrylate, and unreacted fructose,(c) NMR reactants and resulting fructose mono-acrylate, (d) integrations of NMR shifts normalized to characteristic acrylate shifts for TFEA and fructose mono-acrylate, and normalized to fructose hydroxyl shifts for the fructose NMR.

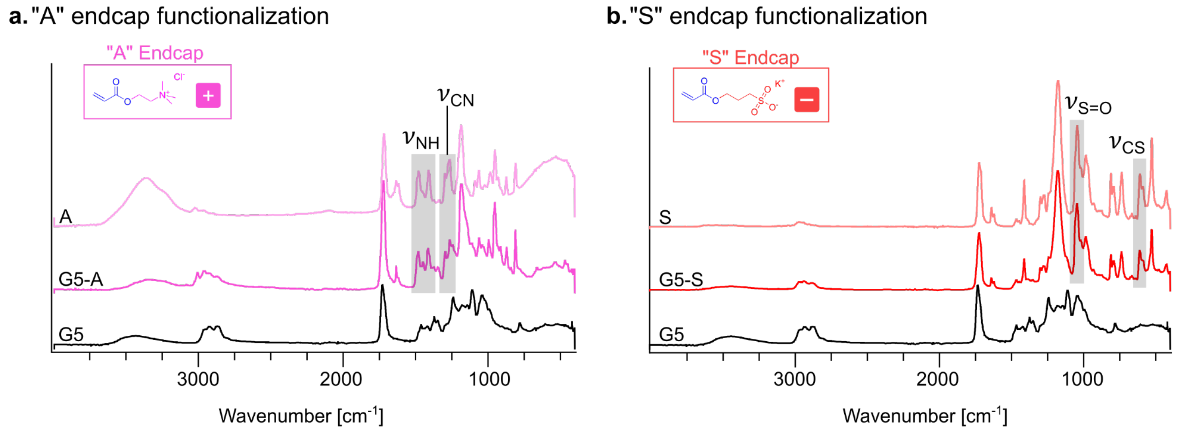


**Supplemental Figure 9: FTIR assessment of dendrimer functionalization with constitutively charged endcaps.** (a) FTIR spectra of unreacted A endcap, A functionalized G5, and unfunctionalized G5 with highlighted stretches corresponding to the N-H bond and C-N bond. (b) FTIR spectra of unreacted S endcap, S functionalized G5, and unfunctionalized G5 with highlighted stretches corresponding to the S=O double bond and C-S bond.

**a.** G5-D H NMR

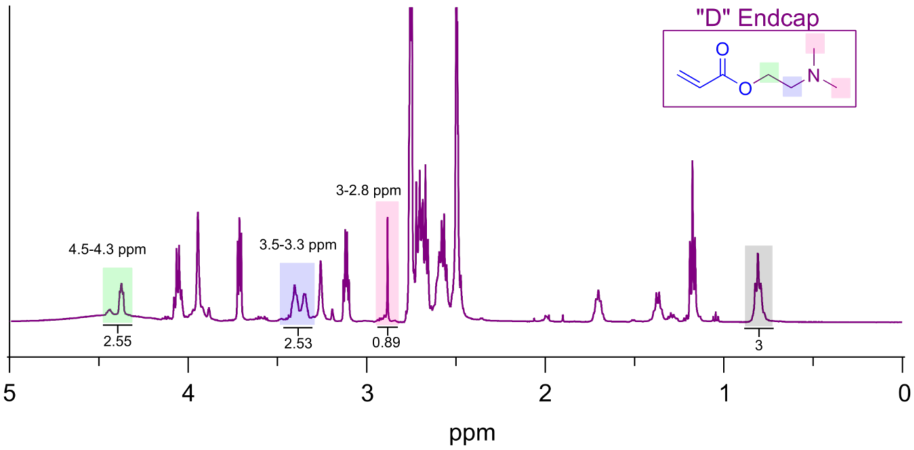


**b.** G5-C H NMR

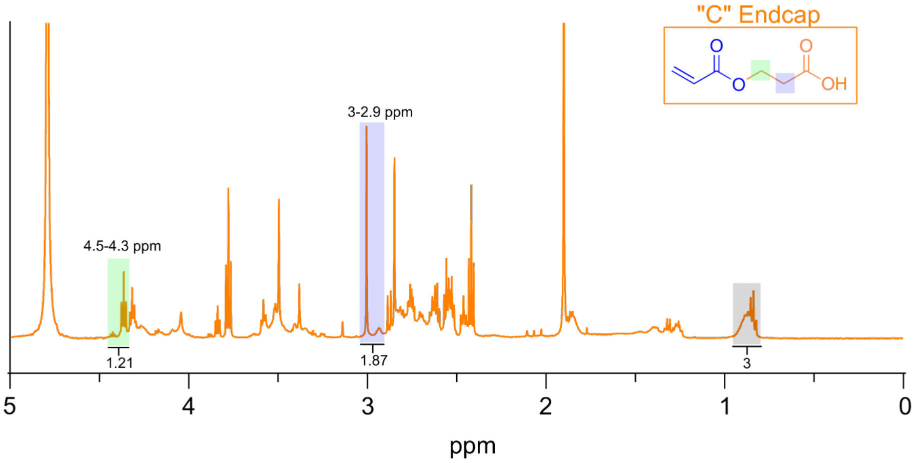


**c.**

| Material | Number of Protons Highlighted | Theoretical | Integrated | % Reaction Completion |
|---|---|---|---|---|
| G5-D | 6 | 6 | 5.97 | 99.5% |
| G5-C | 4 | 4 | 3.08 | 77.0% |

**Supplemental Figure 10: NMR assessment of D endcap vs C endcap functionalization reaction efficiency.** (a) Proton NMR spectra of G5-D material with shifts related to D endcap structure highlighted and integrated. (b) Proton NMR spectra of G5-C material with shifts related to C endcap structure highlighted and integrated. (c) Table showing total integrations of endcap shifts and compared against theoretical integration based on proton number.

**a.** H NMR

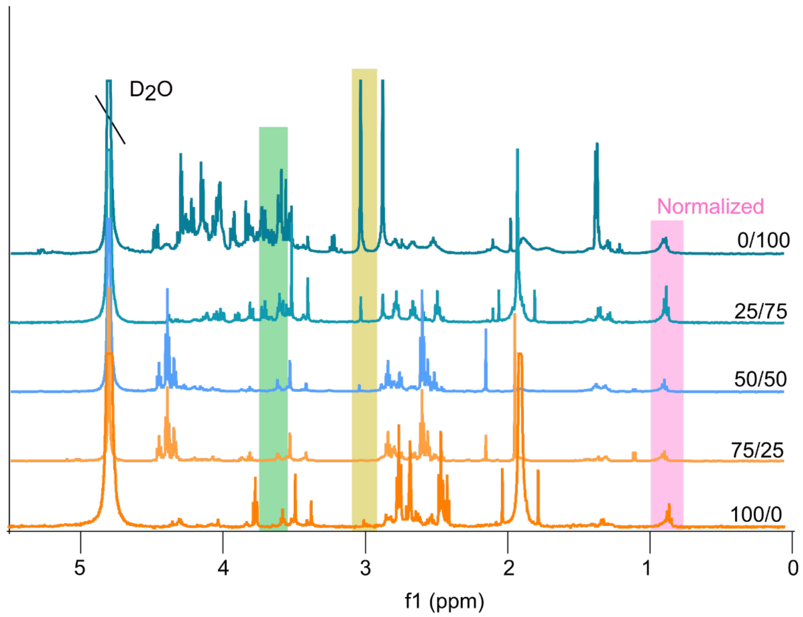


**b.** Predicted Protons

| Shift: | 0.75-1 | 2.8-3.1 | 3.5-3.8 |
|---|---|---|---|
| C100/F0 | 3 | 4 | 0 |
| C75/F25 | 3 | 5 | 1.5 |
| C50/F50 | 3 | 6 | 3 |
| C25/F75 | 3 | 7 | 4.5 |
| C0/F100 | 3 | 8 | 6 |

**c.** Baseline Corrected Shift Integrations

| Shift: | 0.75-1 | 2.8-3.1 | 3.5-3.8 |
|---|---|---|---|
| C100/F0 | 3 | 1.87 | 0 |
| C75/F25 | 3 | 5.52 | 1.48 |
| C50/F50 | 3 | 9.41 | 2.99 |
| C25/F75 | 3 | 15.82 | 6.08 |
| C0/F100 | 3 | 23.82 | 15.34 |

**d.**

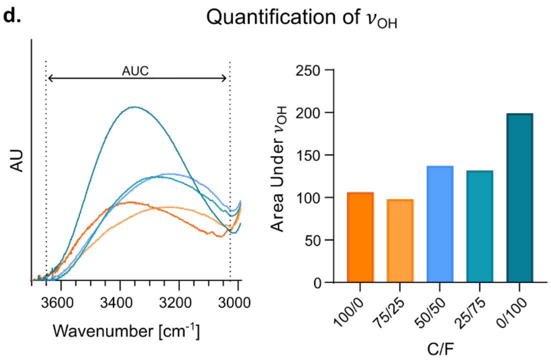


**e.**

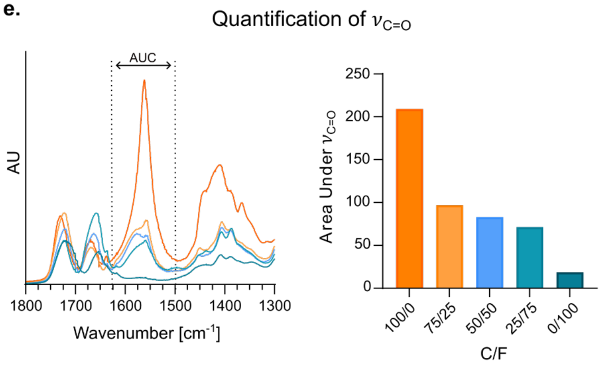


**Supplemental Figure 11: NMR assessment of C/F functionalization ratio.** (a) Proton NMR spectra of G5 dendrimers functionalized with reactant molar ratios of C and F acrylates of 0/100, 25/75, 50/50, 75/25, and 100/0, with spectra normalized to the CH3 protons that occur at 0.8ppm. (b) Predicted number of protons at each of the highlighted shift locations across all formulations. (c) Actual integrations of highlighted shifts when normalized to the CH3 protons and baseline corrected. (d) Quantification of area under the curve (AUC) of the FTIR OH stretch across C/F formulations. (e) Quantification of area under the curve (AUC) of the FTIR C=O double bond stretch across C/F formulations. Panels d and e correspond to the FTIR spectra found in Figure 3, panel g.

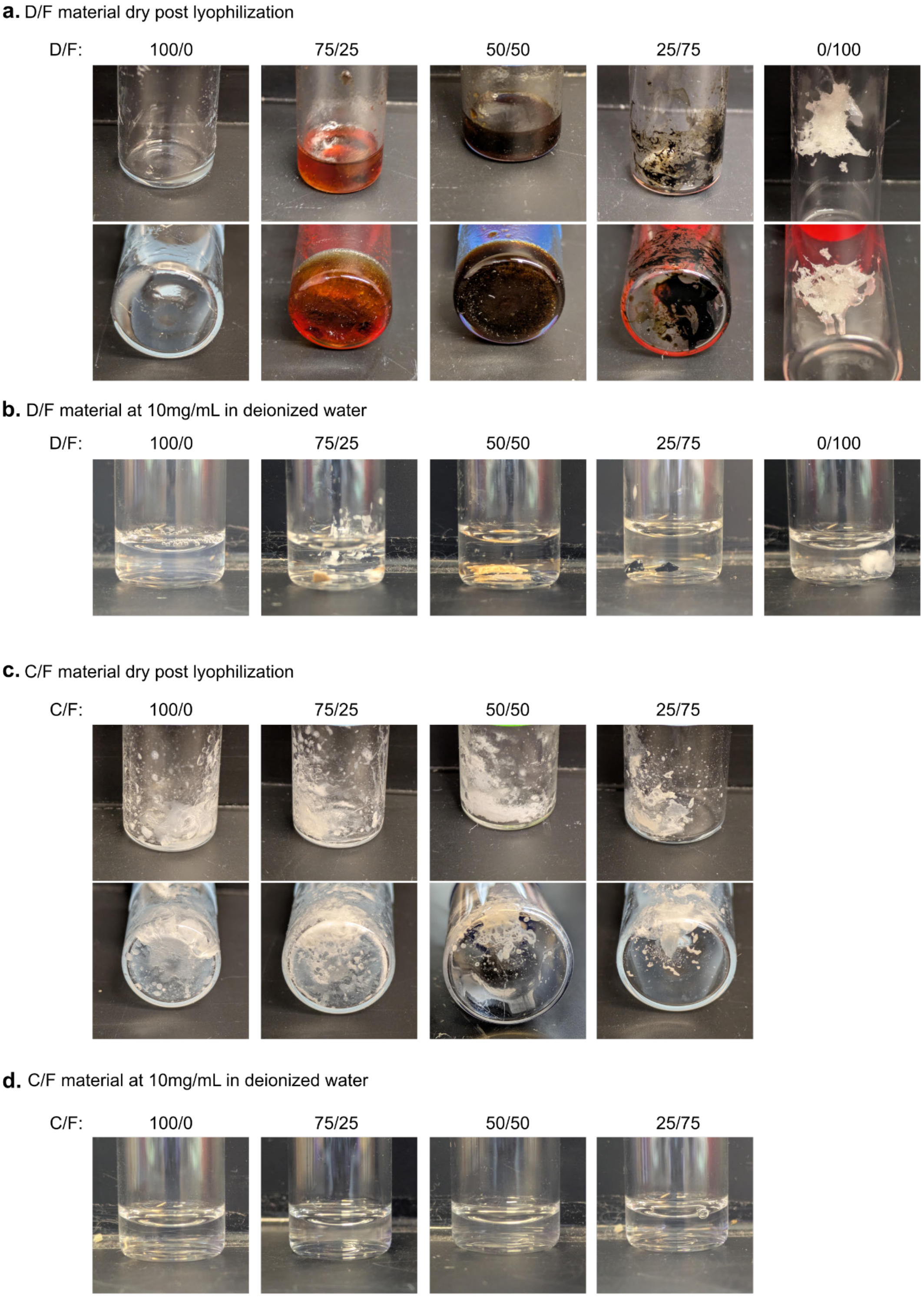


**Supplemental Figure 12: D endcap induced base-catalyzed degradation of the fructose** (a) Images of dry D/F materials demonstrating a color change to dark brown upon the reaction between reducing fructose and D endcap amine groups. (b) Images of D/F material clumping and insoluble in deionized water (c) Images of dry C/F material showing no color change. (d) Images of C/F material fully soluble in deionized water at 10mg/mL.

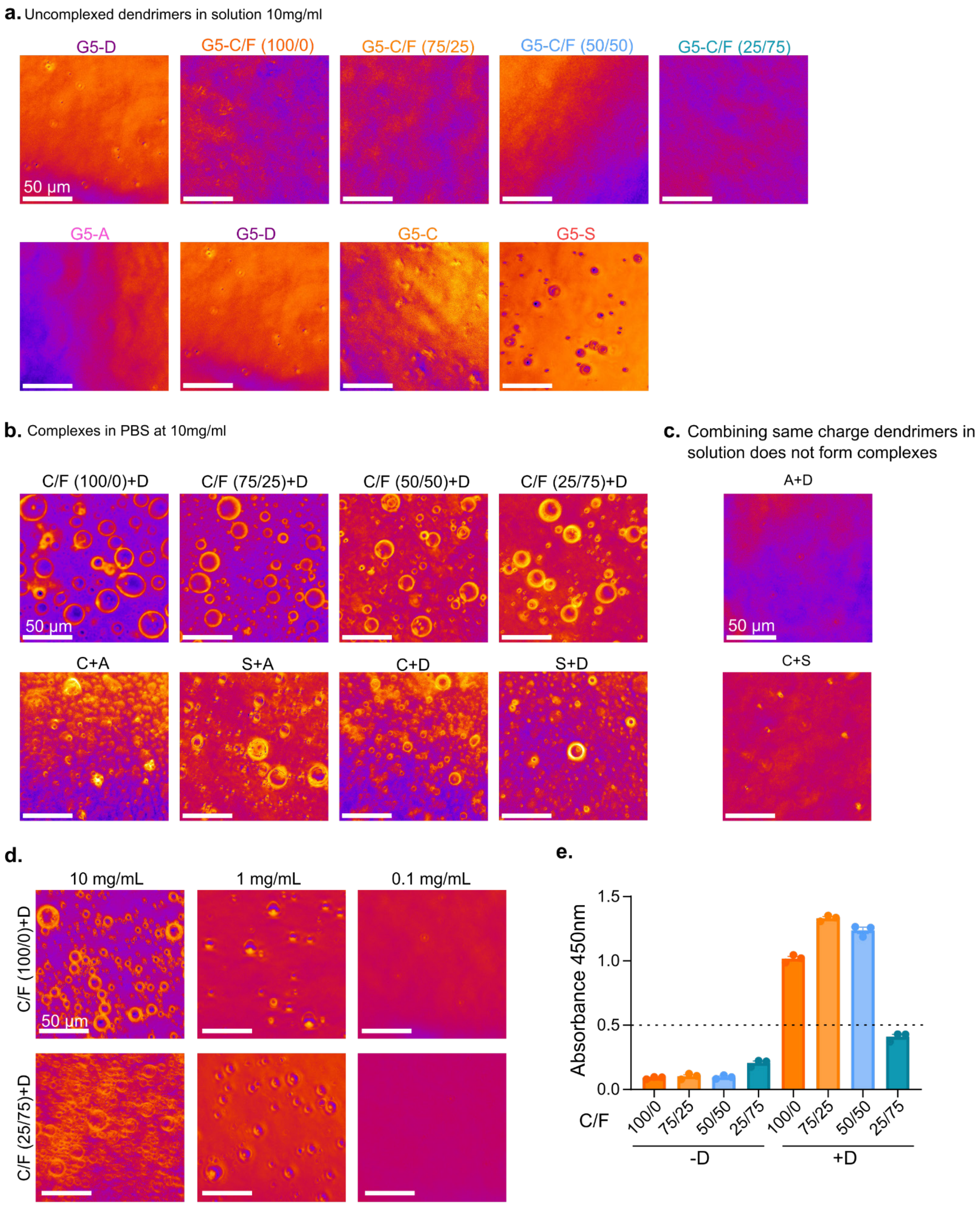


**Supplemental Figure 13: Polyionic complex formation**. (a) Brightfield phase contrasted images of dendrimers functionalized with A, D,C/F, C, and S endcaps solubilized in PBS. (b) Complexes formed due to combinations of oppositely charged dendrimers that are not endcap specific. (c) Brightfield phase contrasted images showing that same-charge combinations do not form PICs. (d) Brightfield phase contrasted images showing that high concentration directly results in larger size droplets, compared to low concentration nanoscale PICs. (e) measure of turbidity via absorbance reading of anionic dendrimers in solution before and after adding a dendrimer of opposite charge, at 1mg/mL, showing mean +/- SEM (standard error of mean). All scale bars are 50 micrometers.

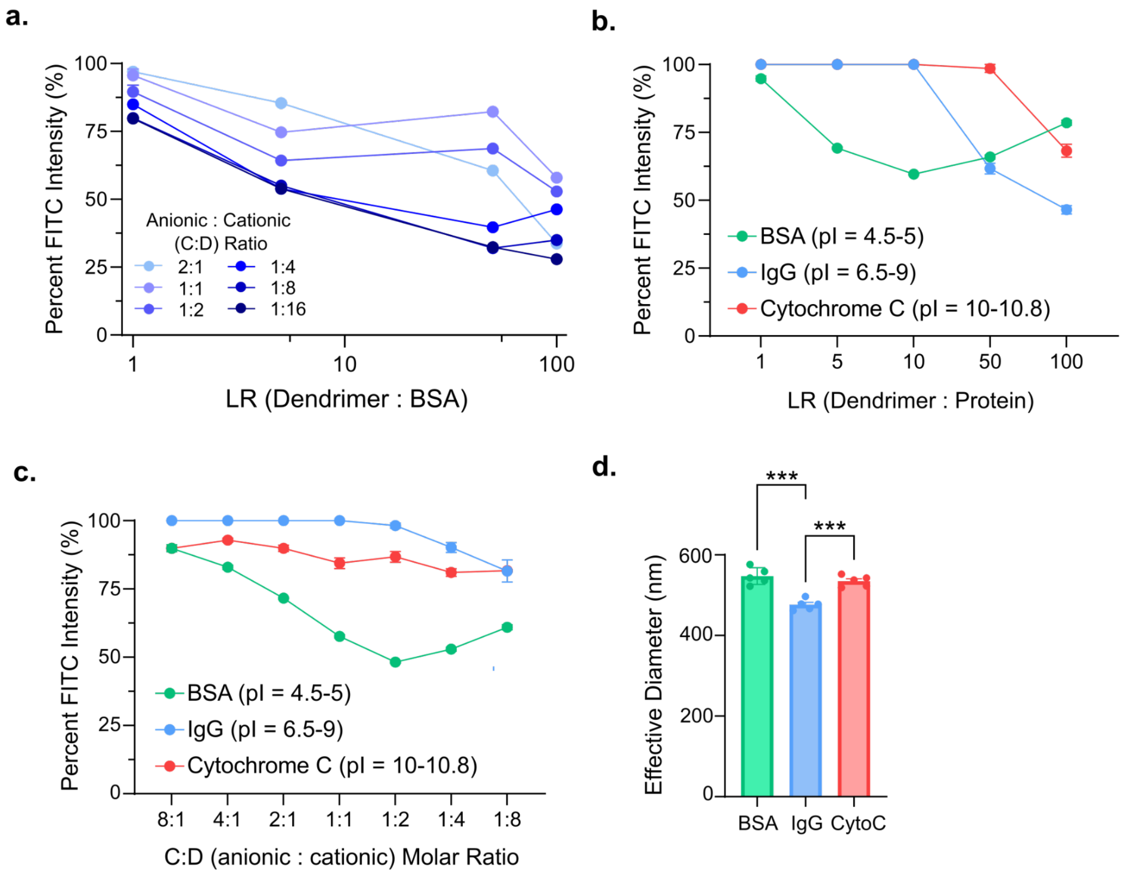


**Supplemental Figure 14: Complexation of protein is dependent on low isoelectric point.** (a) FITC quenching over three orders of magnitude of loading ratio of dendrimer to protein, and over six charge ratios for complexes with F-BSA. (b) FITC quenching over loading ratio of dendrimer to protein for complexes with BSA, IgG, and cytochrome C proteins. (c) FITC quenching over molar ratio of anionic to cationic dendrimers for complexes with BSA, IgG, and cytochrome C proteins. (d) Effective diameter of PICs loaded with BSA, IgG, and cytochrome C proteins at 0.1mg/mL in pH 7.4 PBS. Panel c shows mean +/- SEM (standard error of mean) and was analyzed with one way ANOVA and Tukey's test, ***p-value <0.001.

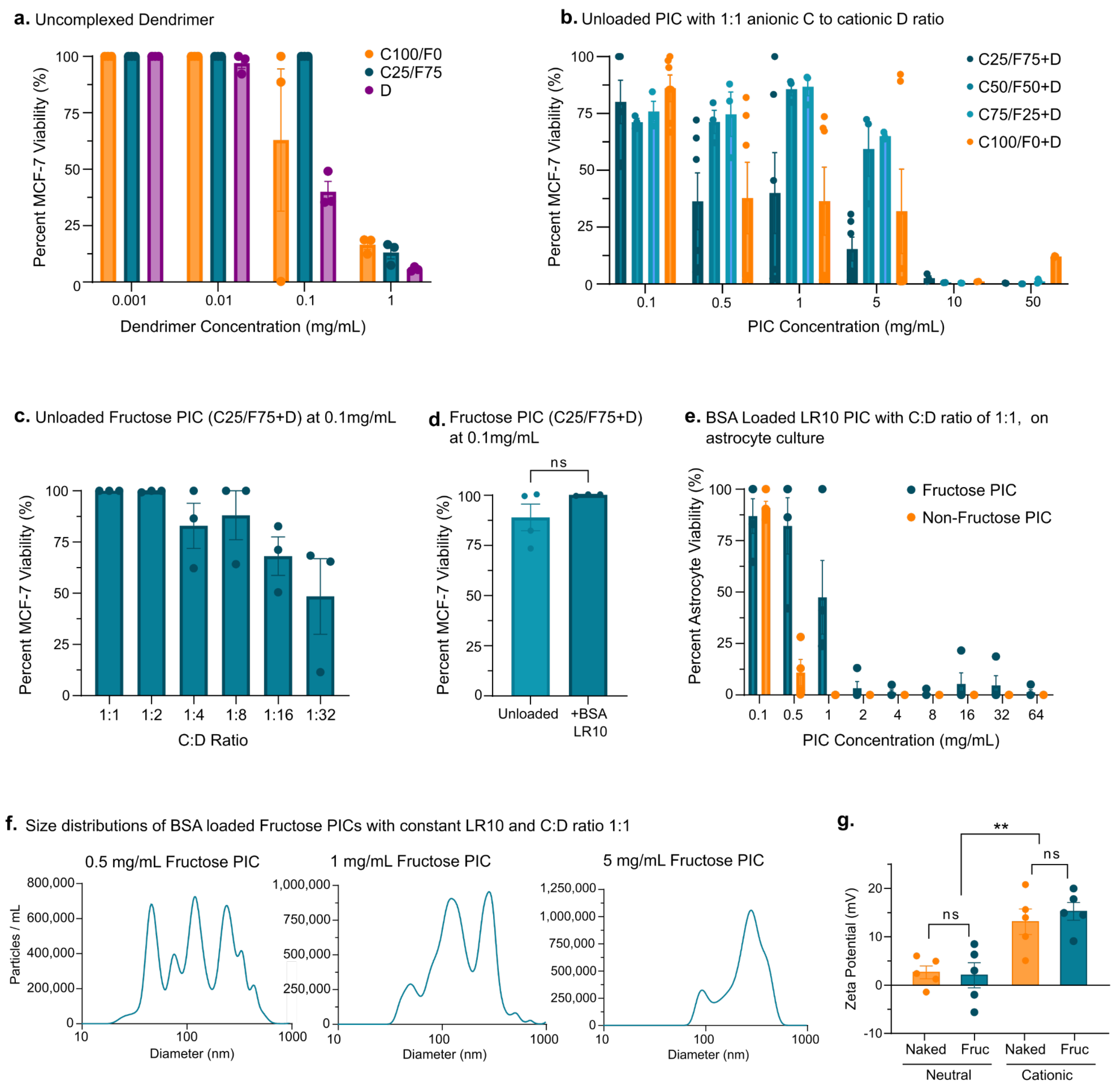


**Supplemental Figure 15: Effects of dendrimers and PICs on viability *in vitro*.** (a) Viability of MCF-7 cells *in vitro* across four orders of magnitude of concentration of uncomplexed dendrimers, the more highly charged materials C100/F0 and D being more toxic than the less charged C25/F75. (b) Viability of MCF-7 cells *in vitro* across concentrations of dendrimers complexed into PICs with varying amounts of fructose presentation. (c) Viability of MCF-7 cells over increasing charge ratios of the fructose PIC with the higher cationic ratios being more toxic than the neutral ratios. (d) Viability of MCF-7 cells *in vitro* when treated with fructose PICs that are either unloaded or loaded with BSA protein at a mass ratio of LR10 (assessed with unpaired t-test). (e) mAstrocyte viability in vitro across concentrations of fructose and non-fructose PICs. (f) NTA evaluation of fructose PIC size distribution across increasing concentrations, showing increasing concentration results in a more uniform population of larger complexes. (g) Zeta potential measured values of neutral and cationic formulations of both naked and fructose PICs.

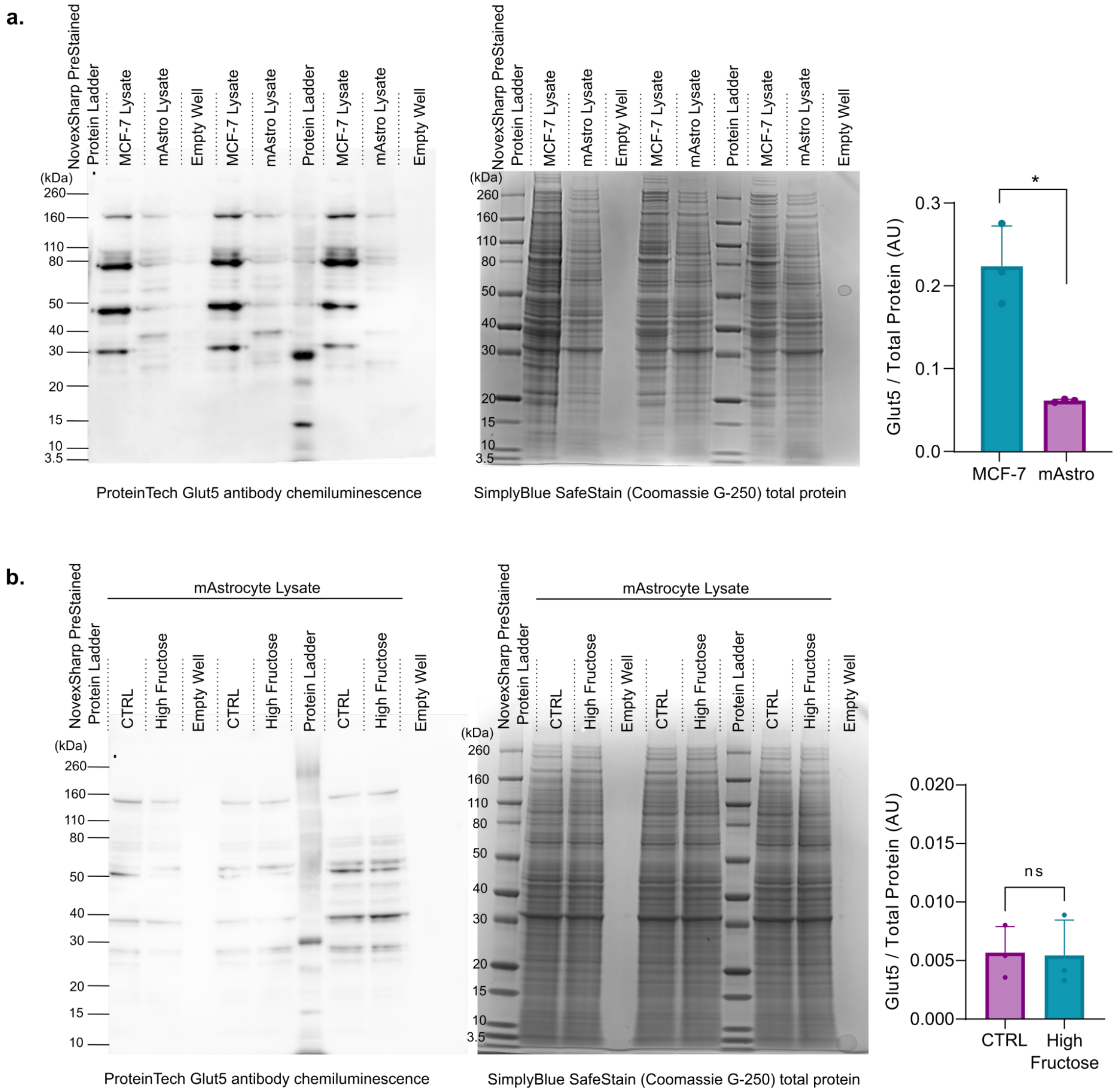


**Supplemental Figure 16: Western blots for evaluation of Glut5 expression *in vitro*.** (a) Glut5 immunostained western blot of MCF-7 and mAstro whole cell lysate with corresponding total protein stain of the same samples. Quantification comparing Glut5 bands intensity normalized over the total protein intensity of the same sample for both MCF-7 and mAstro lysate (n=3). (b) Glut5 immunostained western blot of mAstro whole cell lysate after culturing in either control media or high fructose media, with corresponding total protein stain of the same samples. Quantification comparing Glut5 bands intensity normalized over the total protein intensity of the same sample for mAstro lysate from both media conditions (n=3). Quantifications show mean +/- SEM, analysis done with unpaired t-tests, *p-value <0.05.

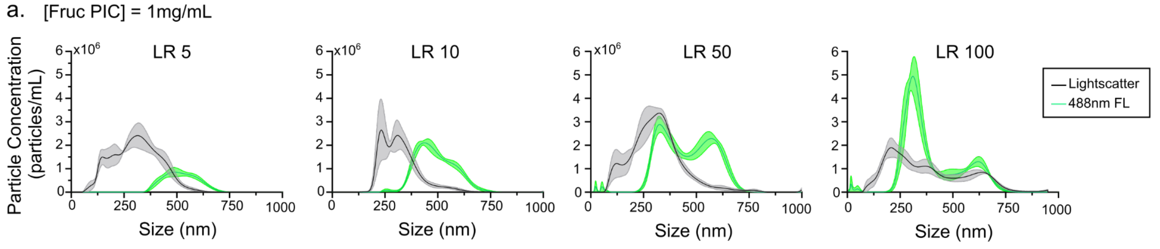

**Supplemental Figure 17: Size characterization of Cre protein loaded fructose PICs** (a) Nanoparticle tracking analysis (NTA) of four different loading ratios of F-BSA loaded PICs, maintaining a constant total dendrimer content concentration at 1mg/mL. Measured with both lightscattering (shown in gray) and 488nm excitation laser module to capture the population of PICs successfully loaded with F-BSA protein (shown in green).

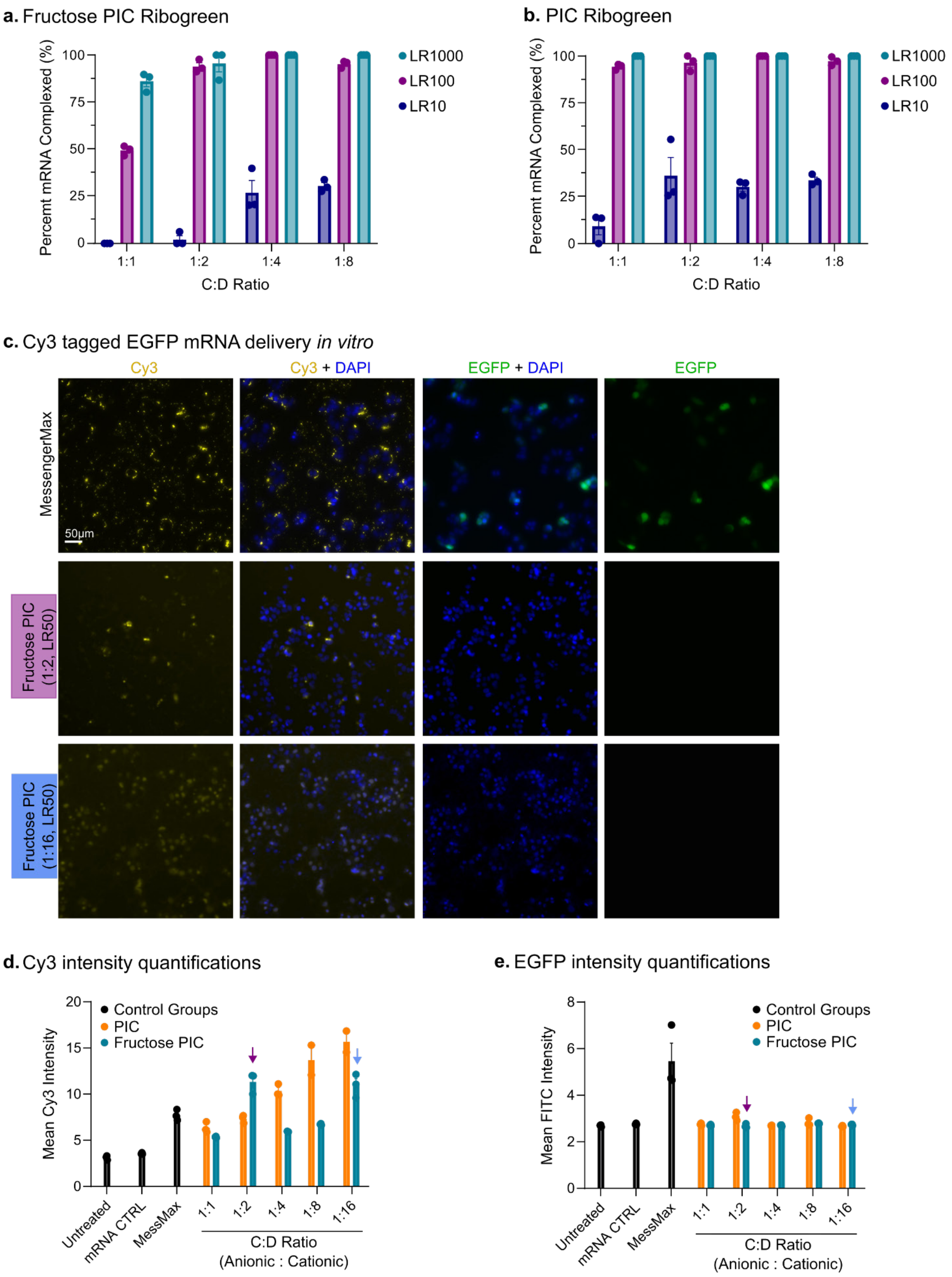


**Supplemental Figure 18: Functional mRNA delivery not achieved *in vitro*.** (a) Assessment of mRNA complexation efficiency with fructose PICs via Ribogreen assay over increasingly cationic ratios of C:D complexation, for three orders of magnitude of dendrimer to mRNA loading ratios. (b) Assessment of mRNA complexation efficiency with non-fructose

PICs via Ribogreen assay over increasingly cationic ratios of C:D complexation, for three orders of magnitude of dendrimer to mRNA loading ratios. (c) Live-imaged MCF-7 cells after treatment with a MessengerMax positive control, and two different charge formulation of fructose PIC, all complexed with Cy3-tagged EGFP coding mRNA, such that green EGFP signal is indicative of functional mRNA intracellular delivery, and stained with NucBlue live DAPI. (d) Quantification of Cy3 signal from live imaging MCF-7 cells post treatment of Cy3-tagged EGFP coding mRNA via MessengerMax, non-fructose PIC and fructose PIC at four different charge ratios with a constant loading ratio of LR50. Arrows indicate quantifications corresponding to images in panel C. (e) Quantification of EGFP signal from live imaging MCF-7 cells post treatment of Cy3-tagged EGFP coding mRNA via MessengerMax, non-fructose PIC and fructose PIC at four different charge ratios with a constant loading ratio of LR50. Arrows indicate quantifications corresponding to images in panel C.

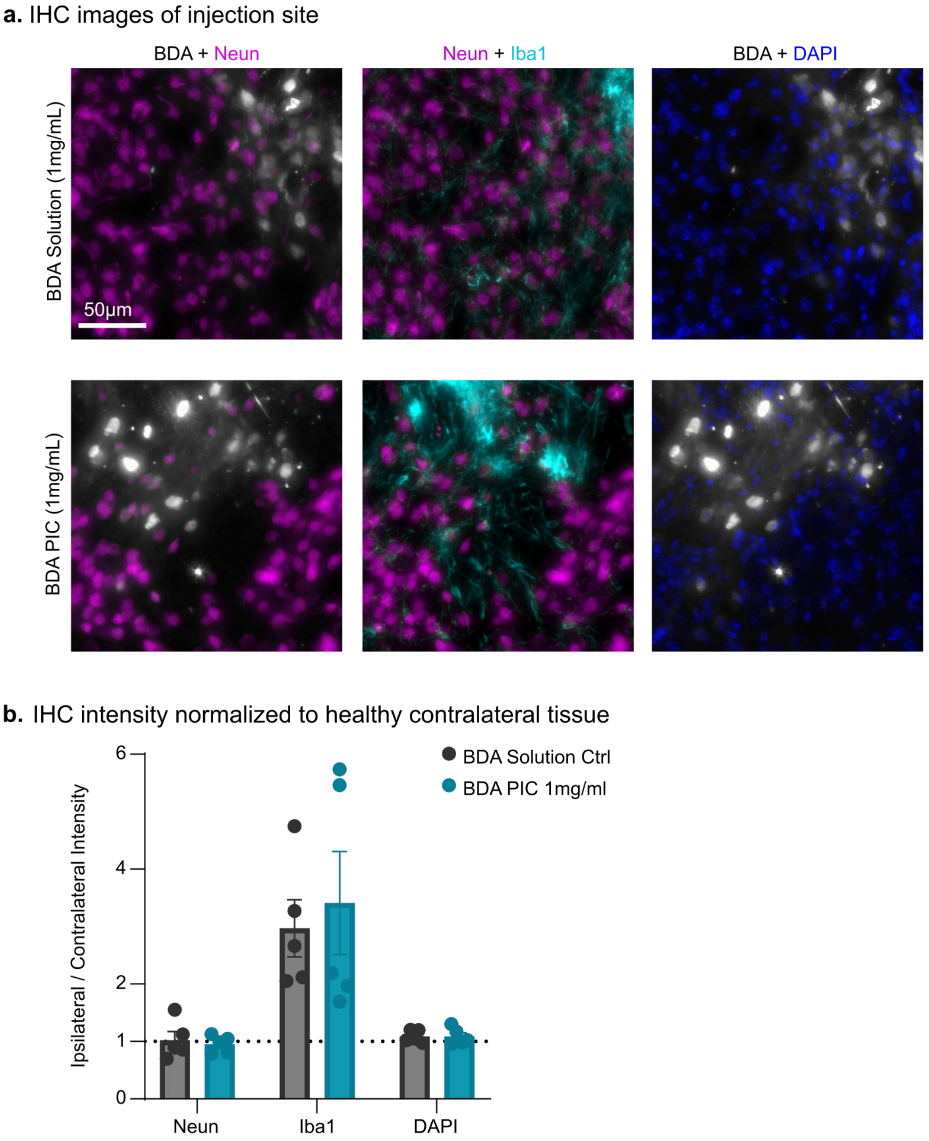


**Supplemental Figure 19: Quantification of BDA-PIC injection site.** (a) 20x magnification IHC images of BDA colocalization on neurons at injection site, comparing the BDA solution control to the 1mg/mL fructose PIC condition. (b) Quantification of NeuN, Iba1, and DAPI of the solution control and 1mg/mL BDA-PIC, normalized to contralateral healthy tissue, demonstrating an increase in microglial reactivity but no loss of neurons or loss of total cell population. A value around 1 indicates that staining at the injection site does not differ in intensity from the contralateral.

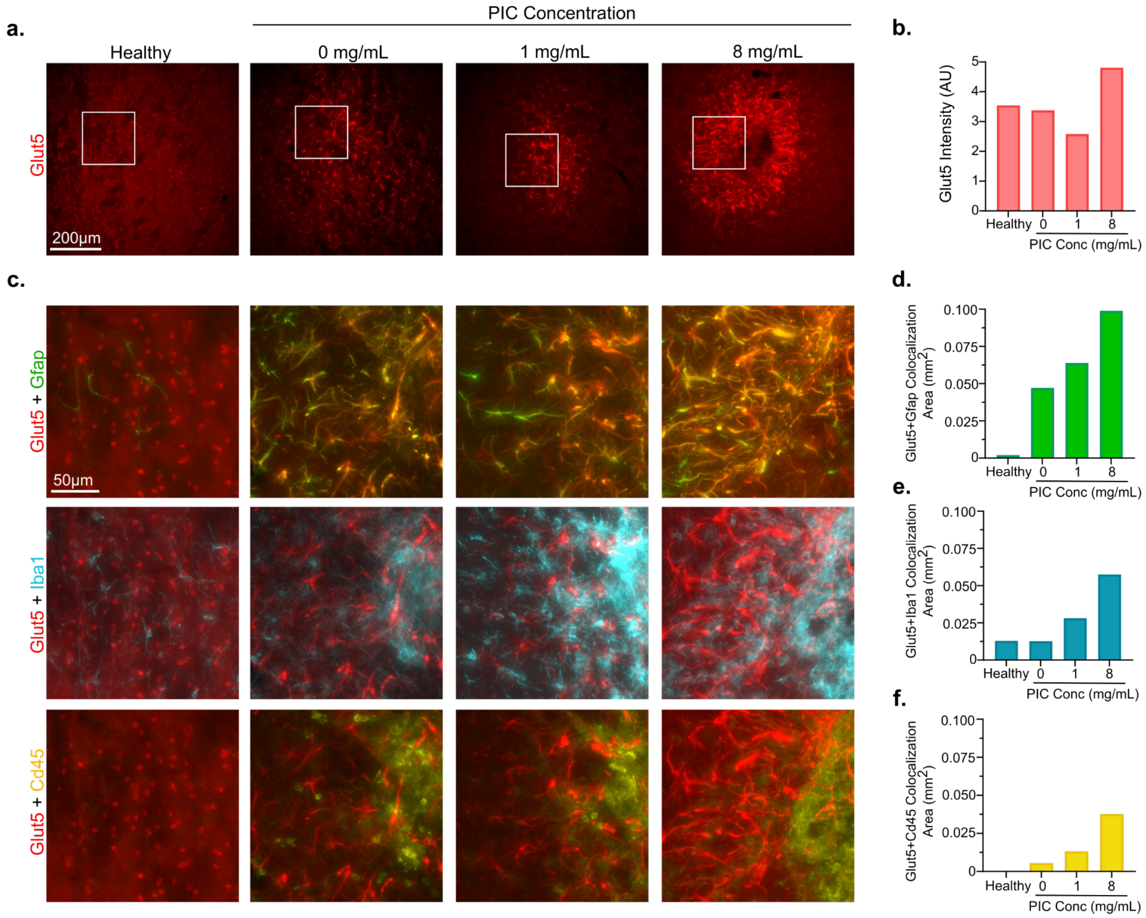


**Supplemental Figure 20: Glut5 expression post BDA PIC injection.** (a) 10x magnification IHC images of Glut5 in healthy striatum tissue and striatum after receiving and injection of either BDA solution control (0mg/mL) or BDA-PIC at a non-toxic concentration (1mg/mL) and a higher concentration that did result in cell loss (8mg/mL). (b) Quantification of the overall staining intensity of Glut5 across the conditions shown in panel a. (c) Higher magnification images corresponding to the white boxes shown in panel a across all four conditions, immunostained for Glut5, Gfap+ reactive astrocytes (green), Iba1+ reactive microglia and recruited monocytes (cyan), and CD45+ recruited leukocytes (yellow). (d) Quantification of colocalization of Glut5 and Gfap staining area across all the conditions shown in panel c. (e) Quantification of colocalization of Glut5 and Iba1 staining area across all the conditions shown in panel c. (f) Quantification of colocalization of Glut5 and CD45 staining area across all the conditions shown in panel c.

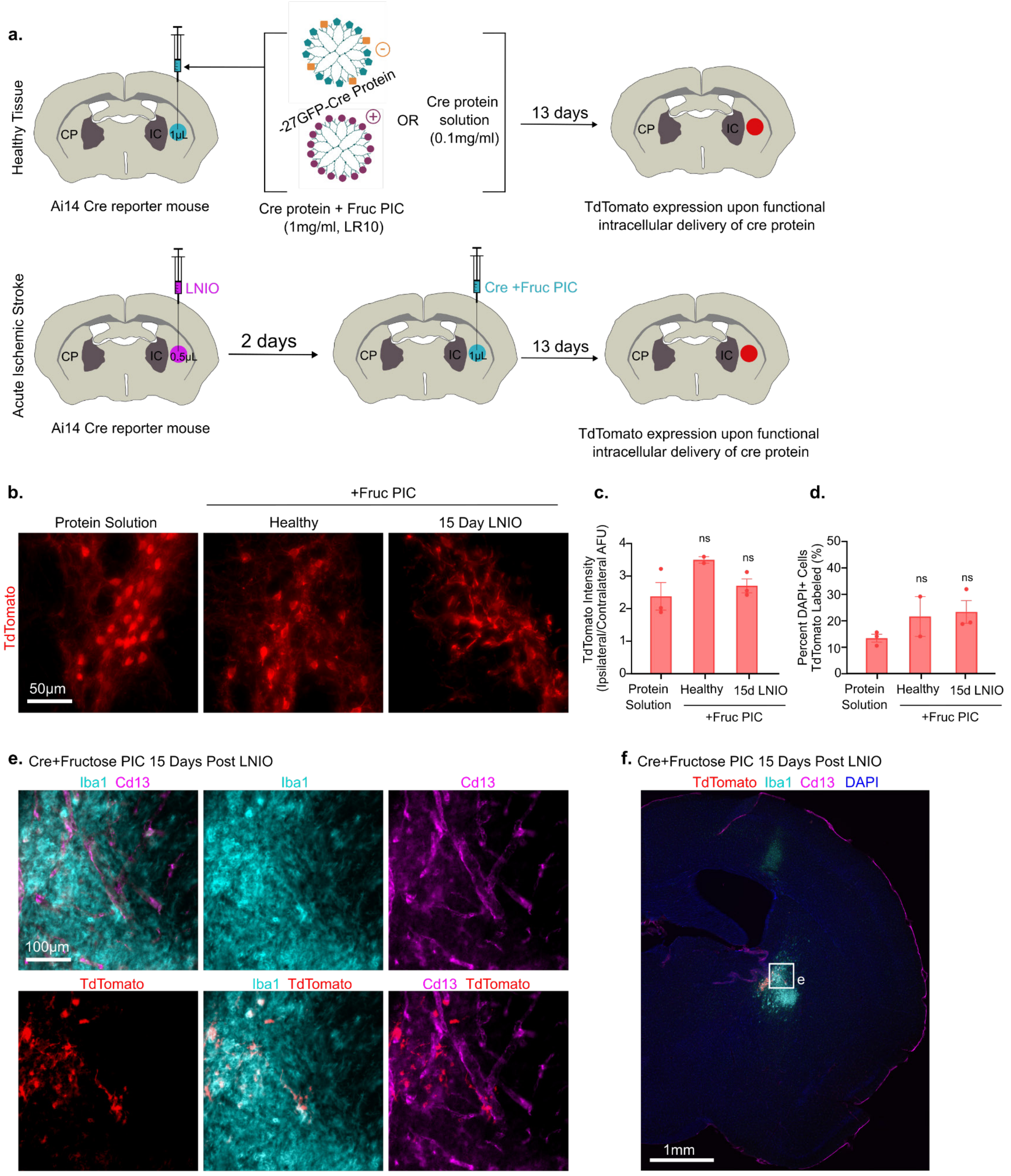


**Supplemental Figure 21: Surgical experimental schematic for Cre loaded Fructose-PICs and IHC.** (a) Schematic of surgical experiment and timeline for healthy and L-NIO animals. (b) IHC images of TdTomato at the injection site for each condition. (c) Quantification of total TdTomato intensity at the injection site normalized to the healthy contralateral tissue for each condition, showing no significant differences. (d) Percentage of DAPI+ cells also labeled with TdTomato in the area of injection, showing no significant differences across conditions.
